# Inference of Single-Cell Phylogenies from Lineage Tracing Data

**DOI:** 10.1101/800078

**Authors:** Matthew G. Jones, Alex Khodaverdian, Jeffrey J. Quinn, Michelle M. Chan, Jeffrey A. Hussmann, Robert Wang, Chenling Xu, Jonathan S. Weissman, Nir Yosef

**Author notes:** Authors Contributed Equally (Matthew G. Jones, Alex Khodaverdian, Jeffrey J. Quinn).

## Abstract

The pairing of CRISPR/Cas9-based gene editing with massively parallel single-cell readouts now enables large-scale lineage tracing. However, the rapid growth in complexity of data from these assays has outpaced our ability to accurately infer phylogenetic relationships. To address this, we provide three resources. First, we introduce Cassiopeia - a suite of scalable and theoretically grounded maximum parsimony approaches for tree reconstruction. Second, we provide a simulation framework for evaluating algorithms and exploring lineage tracer design principles. Finally, we generate the most complex experimental lineage tracing dataset to date - consisting of 34,557 human cells continuously traced over 15 generations, 71% of which are uniquely marked - and use it for benchmarking phylogenetic inference approaches. We show that Cassiopeia outperforms traditional methods by several metrics and under a wide variety of parameter regimes, and provide insight into the principles for the design of improved Cas9-enabled recorders. Together these should broadly enable large-scale mammalian lineage tracing efforts. Cassiopeia and its benchmarking resources are publicly available at www.github.com/YosefLab/Cassiopeia.

## Introduction

The ability to track fates of individual cells during the course of biological processes such as development is of fundamental biological importance, as exemplified by the ground-breaking work creating cell fate maps in *C. elegans* through meticulous visual observation [49, 11]. More recently, CRISPR/Cas9 genome engineering has been coupled with high-throughput single-cell sequencing to enable lineage tracing technologies that can track the relationships between a large number of cells over many generations (Figure 1a, [39]). Generally, these approaches begin with cells engineered with one ore more recording “target sites” where Cas9-induced heritable insertions or deletions (“indels”) accumulate and are subsequently read out by sequencing. A phylogenetic reconstruction algorithm is then used to infer cellular relationships from the pattern of indels. These technologies have enabled the unprecedented exploration of zebrafish [38, 42, 47, 53] and mouse development [31, 7].

**Figure 1:**
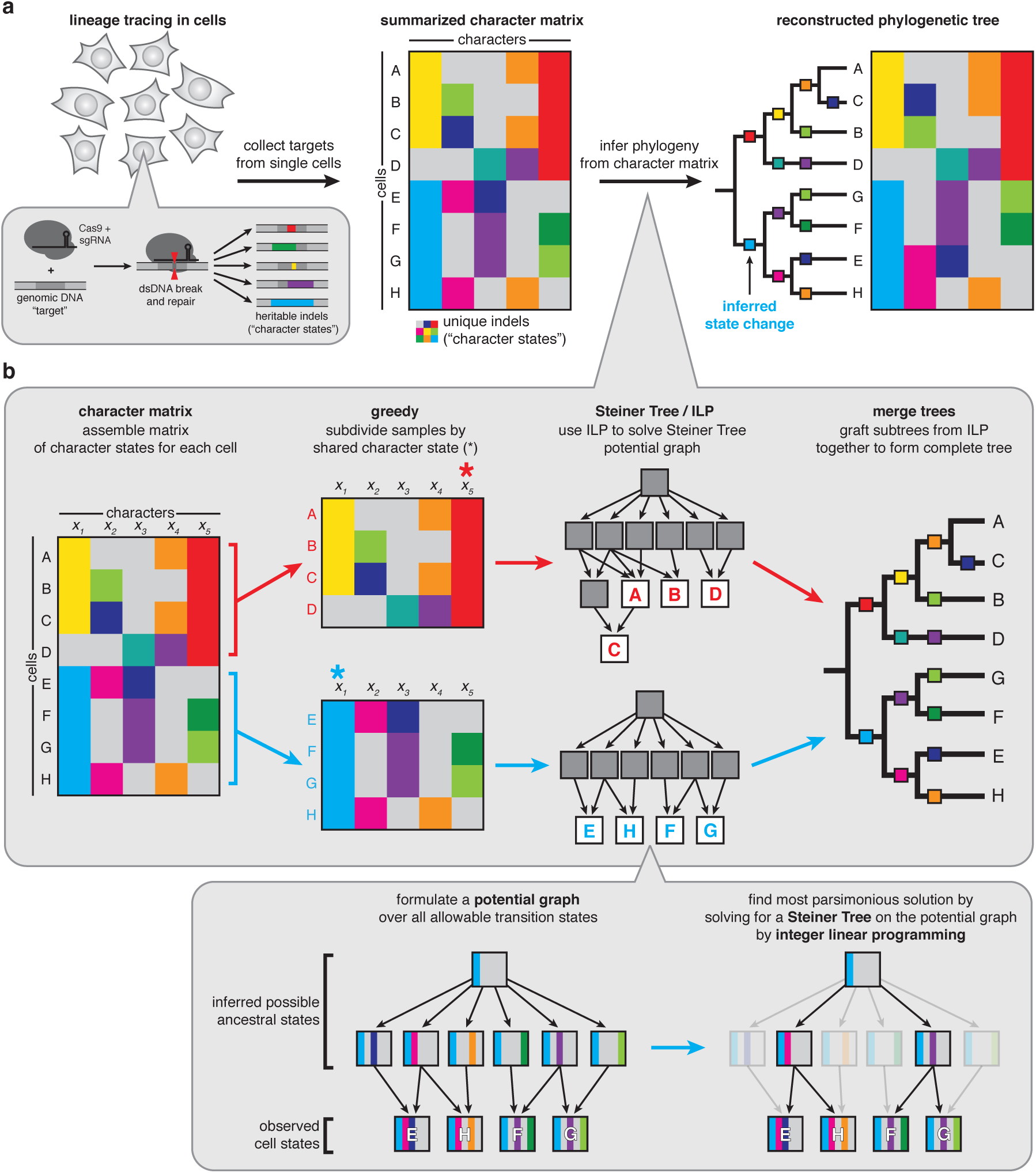
A generalized approach to lineage tracing & lineage reconstruction. (a) The workflow of a lineage tracing experiment. First, cells are engineered with lineage tracing machinery, namely Cas9 that cuts a genomic target site; the target site accrues heritable, Cas9-induced indels (“character states”). Next, the indels are read off from single cells (e.g. by scRNA-seq) and summarized in a “character matrix”, where rows represent cells, columns represent individual target sites (or “characters”) and values represent the observed indel (or “character state”). Finally, the character matrix is used to infer phylogenies by one of various methods. (b) The Cassiopeia framework. Cassiopeia takes as input a “character matrix,” summarizing the mutations seen at heritable target sites across cells. Cassiopeia-Hybrid merges two novel algorithms: the “greedy” (Cassiopeia-Greedy) and “Steiner-Tree / Integer Linear Programming” (Cassiopeia-ILP) approaches. First, the greedy phase identifies mutations that likely occurred early in the lineage and splits cells recursively into groups based on the presence or absence of these mutations. Next, when these groups reach a predefined threshold, we infer Steiner-Trees, finding the tree of minimum weight connecting all observed cell states across all possible evolutionary histories in a “potential graph”, using Integer Linear Programming (ILP). Finally, these trees (corresponding to the maximum parsimony solutions for each group) are returned and merged into a complete phylogeny.

However, the scale and complexity of the data produced by these methods are rapidly becoming a bottleneck for the accurate inference of phylogenies. Specifically, traditional algorithms for reconstructing phylogenies (such as Neighbor-Joining [44] or Camin-Sokal [5]) have not been fully assessed with respect to lineage tracing data and may not be well suited for analyzing large-scale lineage tracing experiments for several reasons. First, traditional algorithms were developed for the cases of few samples (in this case cells) and thus scalability is a major limitation (Supplementary Figure 1). Second, these algorithms are not well suited to handle the amount of missing data from lineage tracing experiments, which can result from either large Cas9-induced resections that remove target sites (“heritable dropout”) or incomplete capture of target sites (“stochastic dropout”). Together, these technical issues necessitate the development of an adaptable approach for reconstructing single-cell phylogenies and an appropriate benchmarking resource that can aid in the development of such algorithms.

Ideally, an algorithm for phylogeny inference from lineage tracing data would be robust to experimental parameters (e.g. rate of mutagenesis, the number of Cas9 target sites), scalable to at least tens of thousands of cells, and resilient to missing data. In this study, we introduce Cassiopeia: a novel suite of three algorithms specifically aimed at reconstructing large phylogenies from lineage tracing experiments with special consideration for the Cas9-mutagenesis process and missing data. Cassiopeia’s framework consists of three modules: (1) a greedy algorithm (Cassiopeia-Greedy), which attempts to construct trees efficiently based on mutations that occurred earliest in the experiment; (2) a near-optimal algorithm that attempts to find the most parsimonious solution using a Steiner-Tree approach (Cassiopeia-ILP); and (3) a hybrid algorithm (Cassiopeia-Hybrid) that blends the scalability of the greedy algorithm and the exactness of the Steiner-Tree approach to support massive single-cell lineage tracing phylogeny reconstruction. To demonstrate the utility of these algorithms, we compare Cassiopeia to existing methods using two resources: first, we benchmark the algorithms using a custom simulation framework for generating synthetic lineage tracing datasets across varying experimental parameters. Second, we assess these algorithms using a new reference *in vitro* lineage tracing dataset consisting of 34,557 cells over 11 clonal populations. Finally, we use Cassiopeia to explore experimental design principles that could improve the next generation of Cas9-enabled lineage tracing systems.

## Results

### Cassiopeia: A Scalable Framework for Single-Cell Lineage Tracing Phylogeny Inference

Typically, phylogenetic trees are constructed by attempting to optimize a predefined objective over characters (i.e. target sites) and their states (i.e. indels) [57]. Distance-based methods (such as Neighbor-Joining [44, 18, 40] or phylogenetic least-squares [6, 17]) aim to infer a weighted tree that best approximates the dissimilarity between nodes (i.e., the number of characters differentiating two cells should be similar to their distance in the tree). Alternatively, character-based methods aim to infer a tree of maximum parsimony [16, 12]. Conventionally, in this approach the returned object is a rooted tree (consisting of observed “leaves” and unobserved “ancestral” internal nodes) in which all nodes are associated with a set of character states such that the overall number of changes in character states (between ancestor and child nodes) is minimized. Finally, a third class of methods closely related to character-based ones takes a probabilistic approach over the characters using maximum likelihood [14, 41] or posterior probability [27] as an objective.

We chose to focus our attention on maximum parsimony-based methods due to the early success of applying these methods to lineage tracing data [42, 38] as well as the wealth of theory and applications of these approaches in domains outside of lineage tracing [35]. Our framework, Cassiopeia, consists of three algorithms for solving phylogenies. In smaller datasets, we propose the use of a Steiner-Tree approach (Cassiopeia-ILP) [59] for finding the maximum parsimony tree over observed cells. Steiner Trees have been extensively used as a way of abstracting network connectivity problems in various settings, such as routing in circuit design [22], and have previously been proposed as a general approach for finding maximum parsimony phylogenies [37, 54]. To adapt Steiner-Trees to single-cell lineage tracing, we devised a method for inferring a large underlying “Potential Graph” where vertices represent unique cells (both observed and plausible ancestors) and edges represent possible evolutionary paths between cells. Importantly, we tailor this inference specifically to single-cell lineage tracing assays: we model the irrereversibility of Cas9 mutations and impute missing data using an exhaustive approach, considering all possible indels in the respective target sites (see methods). After formulating the Potential Graph, we use Integer Linear Programming (ILP) as a technique for finding near-optimal solutions to the Steiner Tree problem. Because of the NP-Hard complexity of Steiner Trees and the difficult approximation of the Potential Graph (whose effect on solution stability is assessed in Figure S1), the main limitation of this approach is that it cannot in practice scale to very large numbers of cells.

To enable Cassiopeia to scale to tens of thousands of cells, we apply a heuristic-based greedy algorithm (Cassiopeia-Greedy) to group cells using mutations that occurred early in the lineage experiment. Our heuristic is inspired by the idea of “perfect phylogeny” [50, 34] - a phylogenetic regime in which every mutation (in our case, Cas9-derived indels) occurred at most once. For the case of binary characters (i.e., mutated yes/ no without accounting for the specific indel), there exists an efficient algorithm [24] for deciding whether a perfect phylogeny exists and if so, to also reconstruct this phylogeny. However, two facets of the lineage tracing problem complicate the deduction of perfect phylogeny: first, the “multi-state” nature of characters (i.e. each character is not binary, but rather can take on several different states; which makes the problem NP-Hard) [4, 48]; and second, the existence of missing data [25]. To address these issues, we first take a theoretical approach and prove that since the founder cell (root of the phylogeny) is unedited (i.e. includes only uncut target sites) and that the mutational process is irreversible, we are able to reduce the multi-state instance to a binary one so that it can be resolved using a perfect-phylogeny-based greedy algorithm. Though Cassiopeia-Greedy does not require a perfect phylogeny, we also prove that if one does exist in the dataset, our proposed algorithm is guaranteed to find it (Theorem 1). Secondly, Cassiopeia-Greedy takes a data-driven approach to handle cells with missing data (see Methods). Unlike Cassiopeia-ILP, Cassiopeia-Greedy is not by design robust to parallel evolution (i.e. “homoplamsy”, where a given state independently arises more than once in a phylogeny in different parts of the tree). However, we demonstrate theoretically that in expectation, mutations observed in more cells are more likely to have occurred fewer times in the experiment for sufficiently small, but realistic, ranges of mutation rates (see Methods; Figure S2), thus supporting the heuristic. Moreover, using simulations, we quantify the precision of this greedy heuristic for varying numbers of states and mutation rates, finding in general these splits are precise (especially in regimes of low mutation rate and realistically large numbers of possible indel outcomes; see Methods and Figure S3). Below, we further discuss simulation-based analyses that illustrate Cassiopeia-Greedy’s effectiveness with varying amounts of parallel evolution (Figure S4).

While Cassiopeia-ILP and Cassiopeia-Greedy are suitable strategies depending on the dataset, we can combine these two methods into a hybrid approach (Cassiopeia-Hybrid) that covers a far broader scale of dataset sizes (Figure 1b). In this use case, Cassiopeia-Hybrid balances the simplicity and scalability of the multi-state greedy algorithm with the exactness and generality of the Steiner-Tree approach. The method begins by splitting the cells into several major clades using Cassiopeia-Greedy and then separately reconstructing phylogenies for each clade with Cassiopeia-ILP. This parallel approach on reasonably sized sub-problems (~ 300 cells in each clade) ensures practical run-times on large numbers of cells (Figure S5). After solving all sub-problems with the Steiner Tree approach, we merge all clades together to form a complete phylogeny.

### A Simulation Engine Enables a Comprehensive Benchmark of Lineage Reconstruction Algorithms

To provide a comprehensive benchmark for phylogeny reconstruction, we developed a framework for simulating lineage tracing experiments across a range of experimental parameters. In particular, the simulated lineages can vary in the number of characters (e.g. Cas9 target sites), the number of states (e.g. possible Cas9-induced indels), the probability distribution over these states, the mutation rate per character, the number of cell generations, and the amount of missing data. We started by estimating plausible “default” values for each simulation parameter using experimental data (discussed below and indicated in Figure 2). In each simulation run, we varied one of the parameters while keeping the rest fixed to their default value. The probability of mutating to each state was found by interpolating the empirical distribution of indel outcomes (Figure S6, see Methods). Each parameter combination was tested using a maximum of 50 replicates or until convergence, each time sampling a set of 400 cells from the total 2^*D*^ cells (where *D* is the depth of the simulated tree).

**Figure 2:**
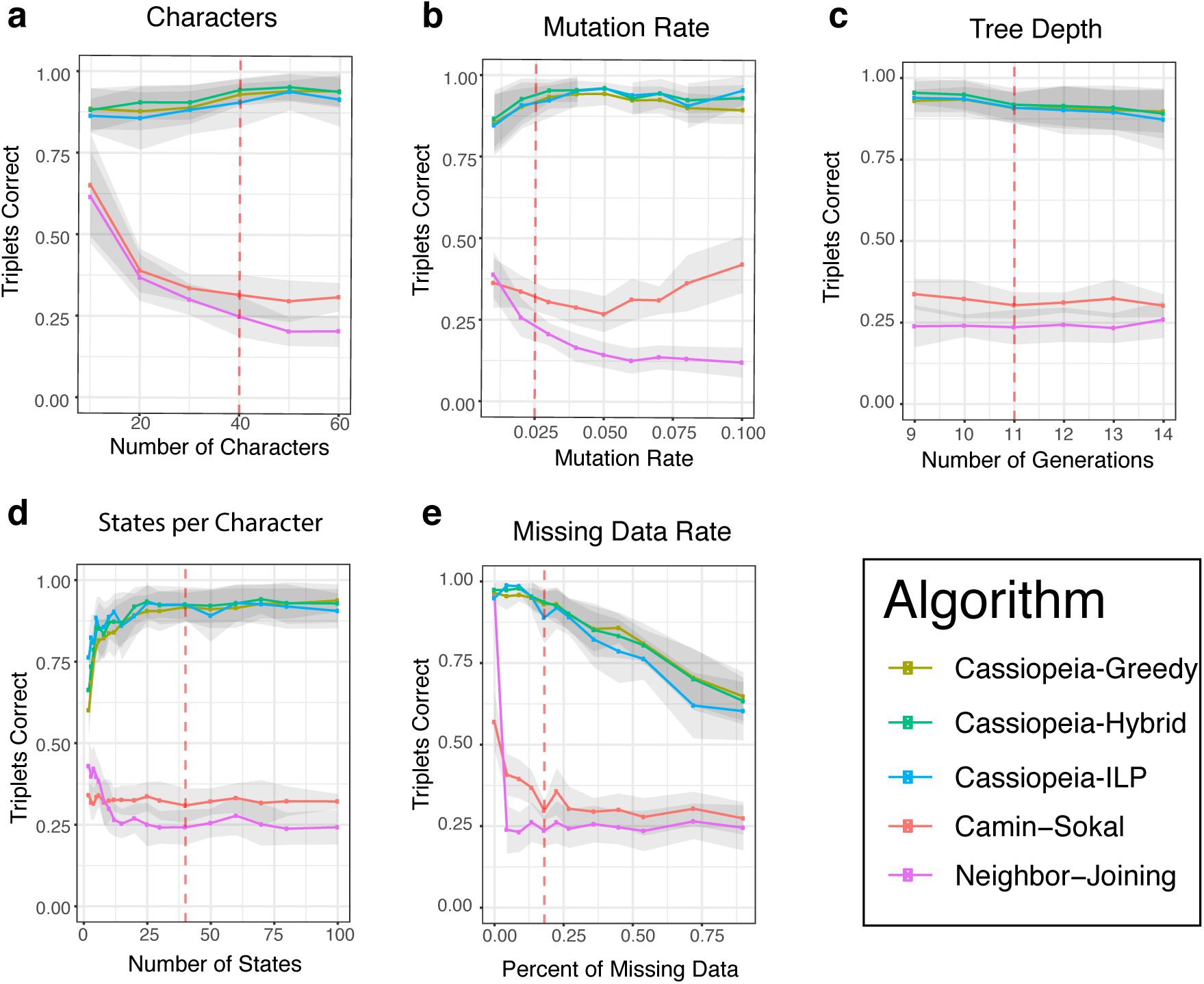
Cassiopeia algorithms outperform other phylogenetic reconstruction methods on simulated lineages. Accuracy is compared between five algorithms (Cassiopeia-Greedy, -ILP, and -Hybrid algorithms as well as Neighbor-Joining and Camin-Sokal) on 400 cells. Phylogeny reconstruction accuracy is assessed with the Triplets correct statistic across several experimental regimes: (a) the number of characters; (b) mutation rate (i.e. Cas9 cutting rate); (c) depth of the tree (or length of the experiment); (d), the number of states per character (i.e. number of possible indel outcomes); and (e) the dropout rate. Dashed lines represent the default value for each stress test. Between 10 and 50 replicate trees were reconstructed, depending on the stability of triplets correct statistic and overall runtime. Standard error over replicates is represented by shaded area.

We compare the performance of our Cassiopeia algorithms (Cassiopeia-ILP, -Greedy, and -Hybrid) as well as an alternative maximum-parsimony algorithm, Camin-Sokal (previously used in lineage tracing applications [38, 42]), and the distance-based algorithm Neighbor-Joining. We assess performance using a combinatoric metric, “Triplets Correct” (Figure S7, see Methods), which compares the proportion of cell triplets that are ordered correctly in the tree. Importantly, this statistic is a weighted-average of the triplets, stratified by the depth of the triplet (measured by the distance from the root to the Latest Common Ancestor (LCA); see Methods). As opposed to other tree comparison metrics, such as Robinson-Foulds [43], we reason that combinatoric metrics [10] more explicitly address the needs of fundamental downstream analyses, namely determining evolutionary relationships between cells.

Overall, our simulations demonstrate the strong performance and efficiency of Cassiopeia. Specifically, we see that the Cassiopeia suite of algorithms consistently finds more accurate trees as compared to both Camin-Sokal and Neighbor-Joining (Figure 2a-e, Figure S8a-e). Furthermore, not only are trees produced with Cassiopeia more accurate than existing methods, but also more parsimonious across all parameter ranges - serving as an indication that the trees reach a more optimal objective solution (Figure S9). Importantly, we observe that Cassiopeia-Hybrid and - Greedy are more effective than Neighbor-Joining in moderately large sample regimes (Figure S10). Notably, Cassiopeia-Greedy and -Hybrid both scale to especially large regimes (of up to 50,000 cells) without substantial compromise in accuracy (S12). In contrast, Camin-Sokal and Cassiopeia-ILP could not scale to such input sizes (Figure S5).

These simulations additionally grant insight into critical design parameters for lineage recording technology. Firstly, we observe that the “information capacity” (i.e. number of characters and possible indels, or states) of a recorder confers an increase in accuracy for Cassiopeia’s modules but not necessarily Camin-Sokal and Neighbor-Joining (Figures 2a,d). This is likely because the greater size of the search space negatively affects the performance of these two algorithms (in other contexts referred to as the “curse of dimensionality” [52]). In addition to the information capacity, we find that indel distributions that tend towards a uniform distribution (thus have higher entropy) allow for more accurate reconstructions especially when the number of states is small or the number of samples is large (Figure S11). Unsurprisingly, the proportion of missing data causes a precipitous decrease in performance (Figure 2e). Furthermore, in longer experiments where the observed cell population is sampled from a larger pool of cells, we find that the problem tends to become more difficult (2c).

Furthermore, these results grant further insight into how Cassiopeia-Greedy is affected in regimes where parallel evolution is likely: such as in low information capacity regimes (e.g. where the number of possible indels is less than 10, Figure 2d), or with high mutation rates (Figure 2b). To strengthen our previous theoretical results suggesting that indels observed in more cells are more likely to occur fewer times and earlier in the phylogeny (Figure S2), we explored how parallel evolution affects Cassiopeia-Greedy empirically with simulation. Specifically, we simulated trees with varying numbers of parallel evolution events at various depths and find overall that while performance decreases with the number of these events, the closer these events occur to the leaves, the smaller the effect (Figure S4). Furthermore, we find that under the “default” simulation parameters (as determined by the experimental data; Figures S6 and 3), Cassiopeia-Greedy consistently makes accurate choices of the first indel event by which cells are divided into clades (Figure S3b).

Practically, the issue of parallel evolution can be addressed to some extent by incorporating state priors (i.e. probabilities of Cas9-induced indel formation). Ideally, Cassiopeia-Greedy would use these priors to select mutations that are low-probability, but observed at high frequency. Theoretically, this would be advantageous as low-probability indels are expected to occur fewer times in the tree (Eq. 1); thus if they appear at high frequency at the leaves, it is especially likely that these occurred earlier in the phylogeny. Furthermore, our precision-analysis indicates that Cassiopeia-Greedy’s decisions are especially precise if it chooses an indel with a low prior (Figure S3). To incorporate these priors in practice, we selected a link function (i.e. one translating observed frequency and prior probability to priority) that maximized performance for Cassiopeia-Greedy (Figure S13; see Methods). After finding an effective approach for integrating prior probabilities, we performed the same stress tests, and found that in cases of likely parallel evolution the priors confer an increase in accuracy (e.g. with high mutation rates; Figure S14), especially in larger regimes (Figure S12).

Here, we have introduced a flexible simulator that is capable of fitting real data, and thus can be used for future benchmarking of algorithms. Using this simulator and a wide range of parameters, we have demonstrated that Cassiopeia performs substantially better than traditional methods. Furthermore, these simulations grant insight into how Cassiopeia’s performance is modulated by various experimental parameters, suggesting design principles that can be optimized to bolster reconstruction accuracy.

### An *In Vitro* Reference Experiment Allows Evaluation of Approaches on Empirical Data

Existing experimental lineage tracing datasets lack a defined ground truth to test against, thus making it difficult to assess phylogenetic accuracy in practice. To address this, we performed an *in vitro* experiment tracking the clonal expansion of a human cell line engineered with a previously described lineage tracing technology [7]. Here, we tracked the growth of 11 clones (each with non-overlapping target site sets for deconvolving clonal populations) over the course of 21 days (approx. 15 generations), randomly splitting the pool of cells into two plates every 7 days (Figure 3a; see Methods). At the end of the experiment, we sampled approximately 10, 000 cells from the four final plates. This randomized plate splitting strategy establishes a course-grained ground truth of how cells are related to each other. Here, cells within the same plate can be arbitrarily distant in their lineage, however there is only a lower bound on lineage dissimilarity between cells in different plates (since they are by definition at least separated by the number of mutations that have occurred since the last split). Thus, it is expected to see some cells more closely related across plates than within (Figure 3a, right), and indels relating these cells across plates are likely to have occurred before the split. However, on average we expect cells within the same plate to be closer to each other in the phylogeny than cells from different plates.

**Figure 3:**
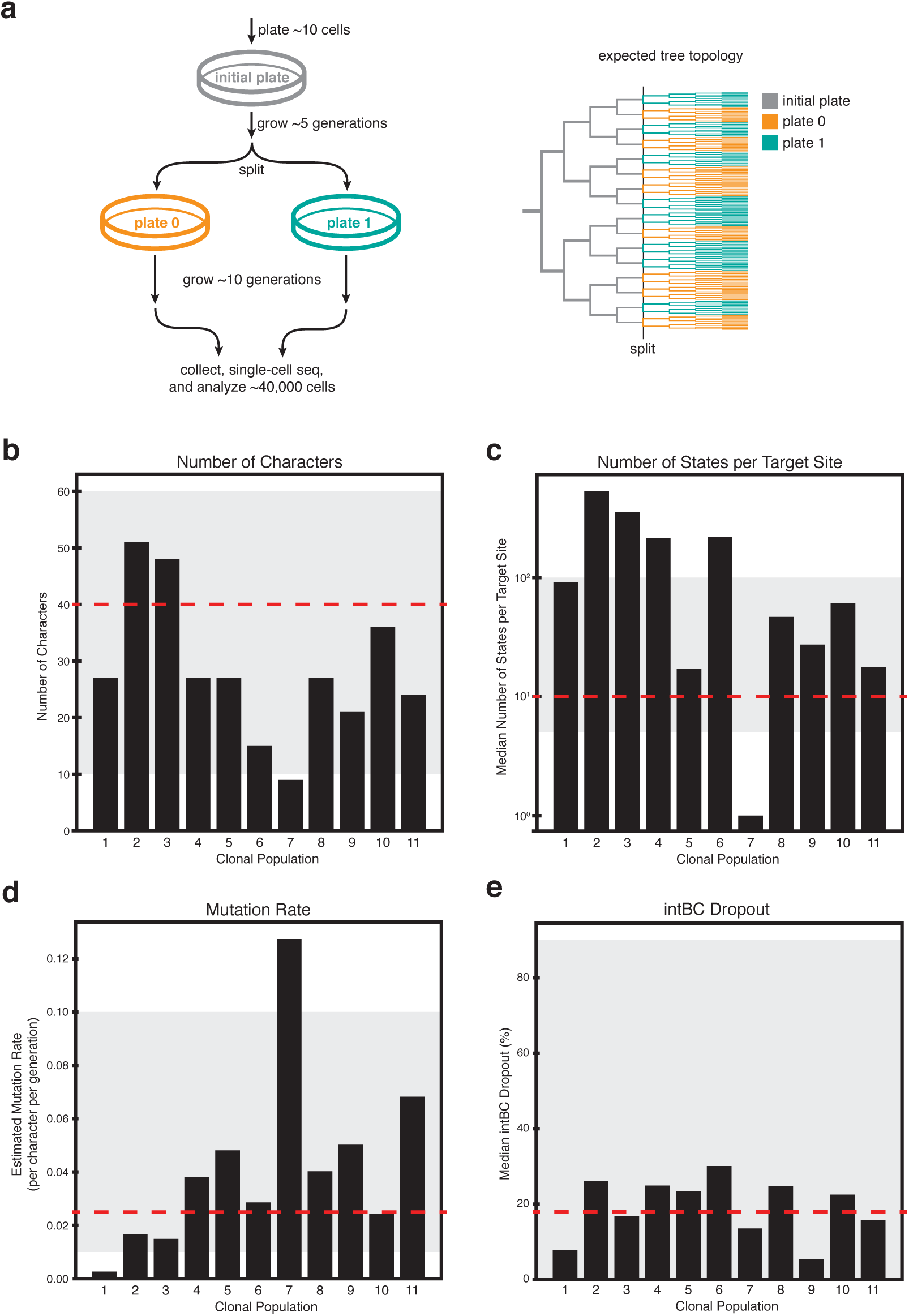
An *in vitro* reference experiment. (a) A reference lineage tracing dataset was generated using the technology proposed in Chan et al. [7] to human cells cultured in vitro for ~ 15 generations. A total of 34, 557 cells were analyzed after filtering and error correction. (b-e) Summary of relevant lineage tracing parameters for each clonal population in the experiment: (b) the number of characters per clone; (c) number of states per target site; (d) the estimated mutation rate per target site; and (e) median dropout per target site. Gray shading denotes parameter regimes tested in simulations and red-dashed lines denote the default values for each synthetic benchmarks.

Our lineage recorder is based on a constitutively expressed target sequence consisting of three evenly spaced cut sites (each cut site corresponding to a character) and a unique integration barcode (“intBC”) which we use to distinguish between target sites and thus more accurately relate character states across cells (Figure S15a). The target sites are randomly integrated into the genomes of founder cells at high copy number (on average 10 targets per cell or a total of 30 independently evolving characters; Figure 3b, S16c). We built upon the processing pipeline in our previous work [7] to obtain confident indel information from scRNA-seq reads (see Methods, Figure S16, Figure S15). In addition, we have added modules for the detection of cell doublets using the sets of intBCs in each clone, and have determined an effective detection strategy using simulations (see Methods, Figure S17). Additionally, we take a data-driven approach for estimating the prior probabilities of indels (see Methods; Figure S18) as other approaches recently proposed [33, 9] may be affected by cell-type and sequence context.

After quality control, error-correction, and filtering we proceeded with analyzing a total of 34, 557 cells across 11 clones. This diverse set of clonal populations represent various levels of indel diversity (i.e. number of possible states, Figure 3c), size of intBC sets (i.e. number of characters, Figure 3b and Figure S16c), character mutation rates (Figure 3d, see Methods), and proportion of missing data (Figure 3e, see Methods). Most importantly, this dataset represents a significant improvement in lineage tracing experiments: it is the longest and most complex dataset to date in which the large majority of cells are uniquely marked (71%), indicating a rich character state complexity for tree building.

We next reconstructed trees for each clone (excluding two which were removed through quality-control filters; see Methods) with our suite of algorithms, as well as Neighbor-Joining and Camin-Sokal (when computationally feasible). For both Cassiopeia-Greedy and Cassiopeia-Hybrid methods, we also compared tree reconstruction accuracy with or without prior probabilities. The tree for Clone 3, consisting of 7,289 cells, along with its character matrix and first split annotations (i.e. whether cells were initially split into plate 0 or plate 1, denoted as the plate ID), is presented in Figure 4. Interestingly, we find that certain indels indeed span the different plates, thus indicating that Cassiopeia-Greedy chooses early indels to serve as splits. Moreover, the character matrix and the nested dissection of the tree demonstrate the abundant lineage information encoded in this clone – 96% of the 7,289 observed cells are marked uniquely.

**Figure 4:**
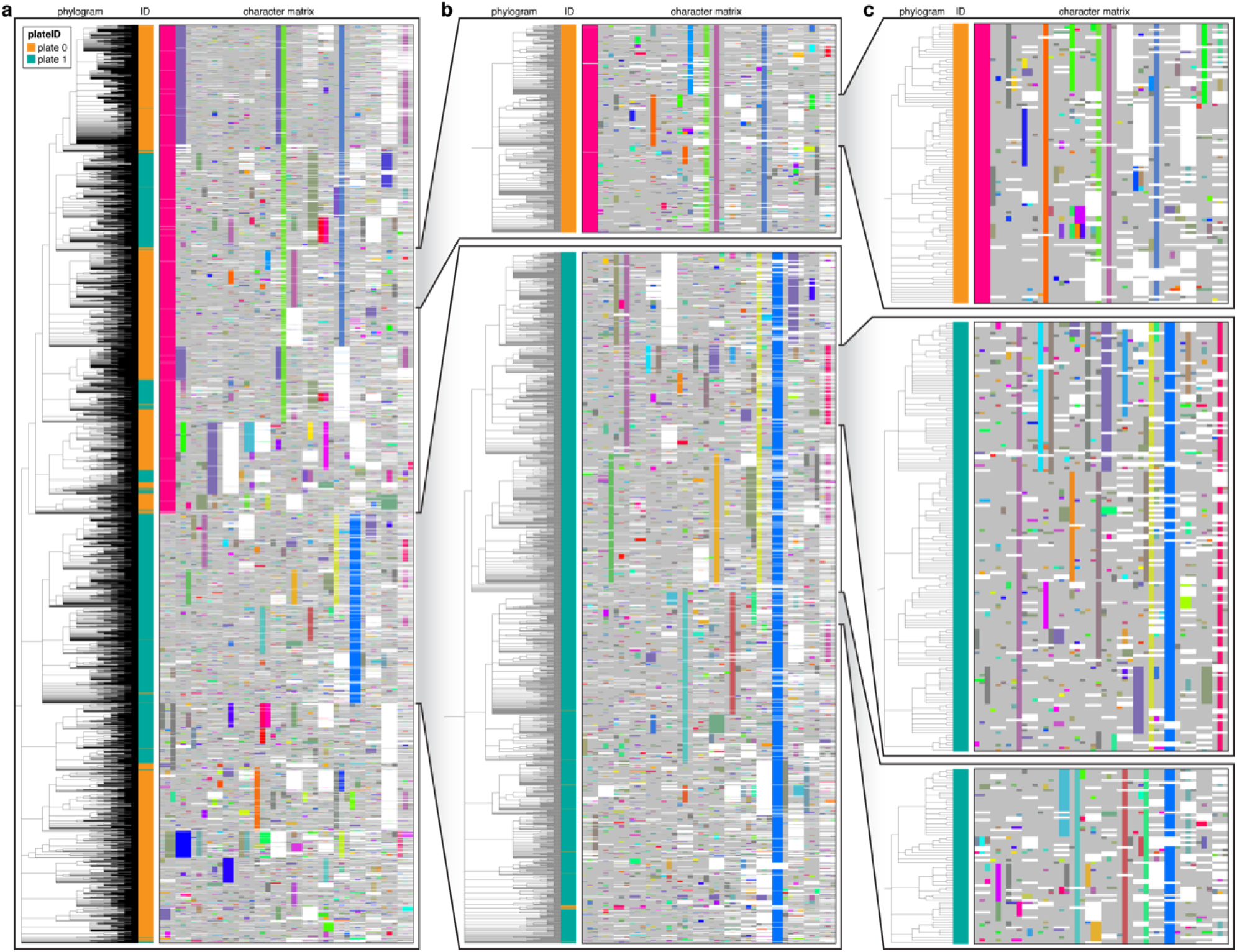
Cassiopeia can reconstruct high-resolution phylogenetic trees from empirical lineage tracing data. The full phylogenetic tree for Clone 3 (a), consisting of 7,289 cells, was reconstructed using Cassiopeia-Hybrid (with priors), and is displayed. The phylogram represents cell-cell relationships, and each cell is colored by sample ID at the first split (plate 0 or 1). The character matrix is displayed with each unique character state (or “indel”) represented by distinct colors. (Light gray represents uncut sites; white represents missing values.) Of these 7,289 cells, 96% were uniquely tagged by their character states. (b-c) Nested, expanded views of the phylogram and character matrices. As expected, Cassiopeia correctly relates cells with similar character states, and closely related cells are found within the same culture plate.

By keeping track of which plate each cell came from we are able to evaluate how well the distances in a computationally-reconstructed tree reflect the distances in the experimental tree. Thus, we test the reconstruction ability of an algorithm using two metrics for measuring the association between plate ID and substructure – “Meta Purity” and “Mean Majority Vote” (see Methods). Both are predicated on the assumption that, just as in the real experiment, as one descends the reconstructed tree, one would expect to find cells more closely related to one another. In this sense, we utilize these two metrics for testing homogeneous cell labels below a certain internal node in a tree, which we refer to as a “clade”.

We use these statistics to evaluate reconstruction accuracy for Clone 3 with respect to the first split labels (i.e. plate 0 or 1, Figure 5). In doing so, we find that Cassiopeia-Greedy and -Hybrid consistently outperform Neighbor-Joining. We find overall consistent results for the remainder of clones reconstructed (Figure S19, Figure S20), although Cassiopeia’s modules have the greatest advantage in larger reconstructions.

**Figure 5:**
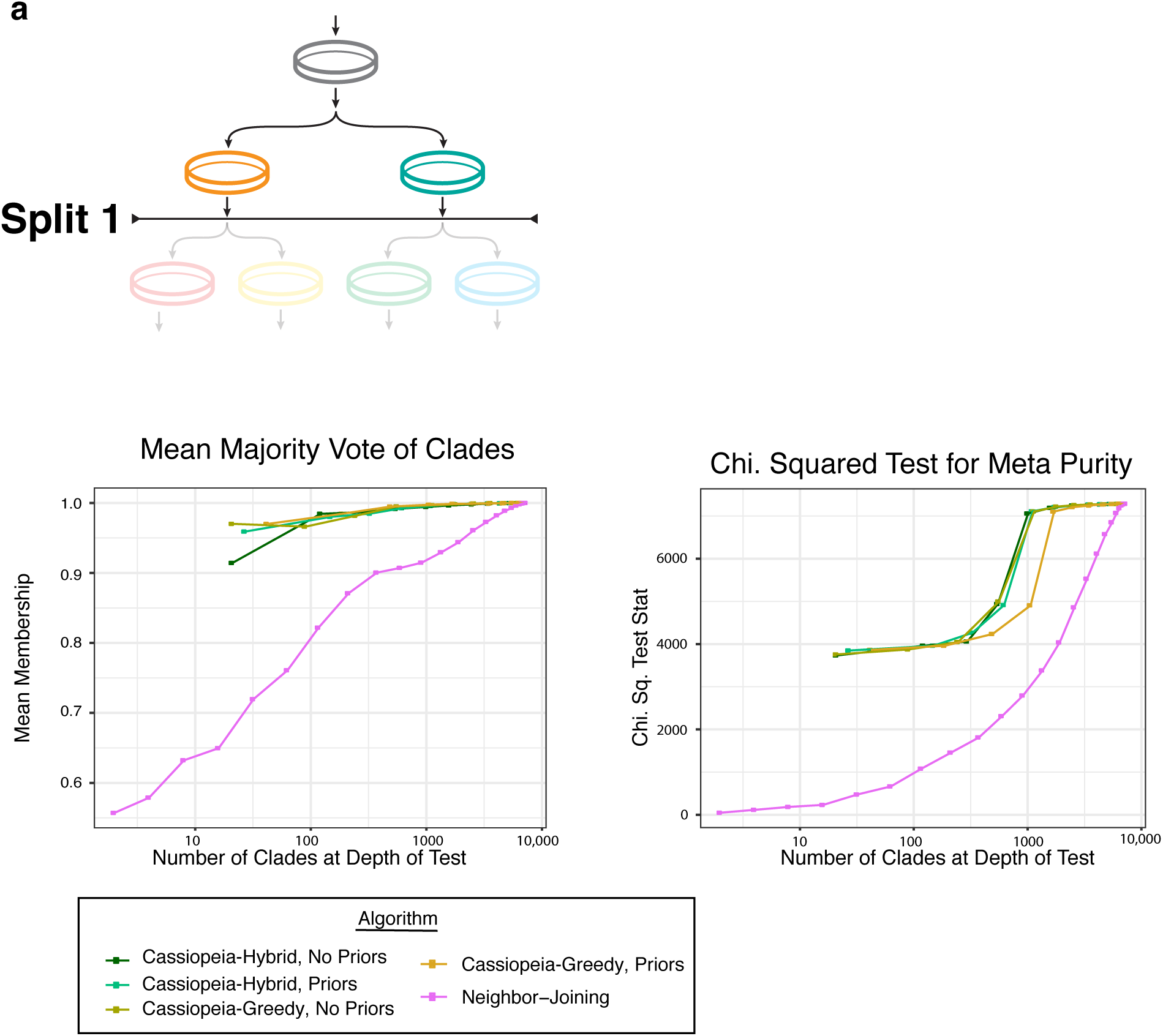
Cassiopeia builds highly accurate trees from large empirical datasets. The consistency between tree reconstructions are evaluated with respect to the first split, represented in (a). The Mean Majority Vote (b) and the Meta Purity test (c) were used for Cassiopeia-Hybrid and -Greedy (both with or without priors) and Neighbor-Joining. The statistics are plotted as a function of the number of clades at the depth of the test (i.e. the number of clades created by a horizontal cut at a given depth). All Cassiopeia approaches consistently outperform Neighbor-Joining by both metrics.

Overall, we anticipate that this *in vitro* dataset will serve as a valuable empirical benchmark for future algorithm development. Specifically, we have demonstrated how this dataset can be used to evaluate the accuracy of inferred phylogenies and illustrate that Cassiopeia outperforms Neighbor-Joining. Moreover, we demonstrate Cassiopeia’s scalability for reconstructing trees that are beyond the abilities of other maximum parsimony-based methods like Camin-Sokal as they currently have been implemented.

### Generalizing Cassiopeia to Alternative & Future Technologies

While previous single-cell lineage tracing applications have proposed methods for phylogenetic reconstruction, they have been custom-tailored to the experimental system, requiring one to filter out common indels [47] or provide indel likelihoods [7]. We thus investigated how well Cassiopeia generalizes to other technologies with reconstructions of data generated with the GESTALT technology [38, 42] (Figure 6a, Figure S21). Comparing Cassiopeia’s algorithms to Neighbor-Joining and Camin-Sokal (as applied in these previous studies [38, 42]), we find that Cassiopeia-ILP consistently finds the most parsimonious solution. While parsimony is not a direct measure of tree accuracy, it is a direct measure of solution optimality and clearly demonstrates Cassiopeia’s effectiveness for existing alternative lineage tracing technologies.

**Figure 6:**
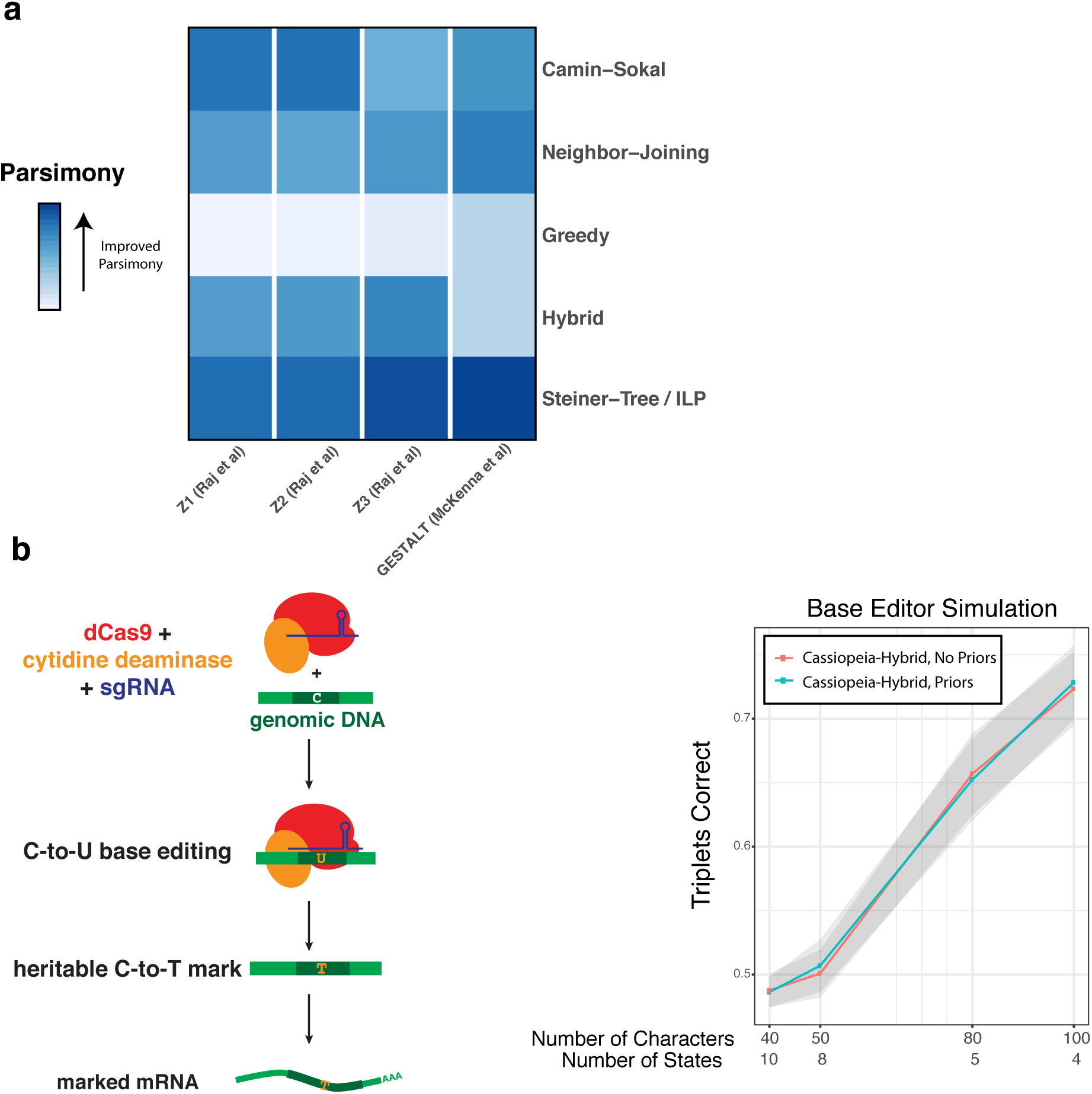
Generalizing Cassiopeia & future design principles of CRISPR-enabled lineage tracers. (a) Cassiopeia generalizes to alternative lineage tracing methods, as illustrated with the analysis of data from GESTALT technology [38, 42]). In a comparison of parsimony across Camin-Sokal, Neighbor-Joining, and Cassiopeia’s methods, the Steiner-Tree approach consistently finds more parsimonious (i.e more optimal) solutions. (b) Exploring information capacity of recorders with base-editors. A theoretical base-editor was simulated for 400 cells and reconstructions with Cassiopeia-Hybrid, with and without priors. We compared the accuracy of the reconstructions to the simulated tree using the triplets correct statistic. We describe the performance of Cassiopeia-Hybrid as the number of characters was increased (and consequently number of states was decreased).

After establishing Cassiopeia’s generalizability, we turned to investigating plausible next-generation lineage tracers. Recently, base-editing systems (Figure 6b) have been proposed to precisely edit *A* > *G* [19], *C* > *T* [36, 20] or possibly *C* > *N* (*N* being any base as in [26]). The promise of base-editing lineage recorders is three-fold: first, a base editor would increase the number of editable sites (as compared to the ones that rely on Cas9-induced double-strand breaks [7, 38, 47]) although at the expense of number of states (at best 4, corresponding to A, C, T, and G). Second, a base-editing system would theoretically result in less dropout, since target site resection via Cas9-induced double-strand breaks is far less likely [36]. Third, it is hypothesized that base-editors would be less cytotoxic as it does not depend on inducing double strand breaks on DNA (although this relies on effective strategies for limiting off-target base-editing of DNA and RNA [56]). To evaluate the application of base editors for lineage tracing, we tested the performance of Cassiopeia in high-character, low-state regimes as would be the case in base editing (Figure 6b, see Methods). Using simulations with parameters deduced by a recent base editor application [26], we demonstrate that there appears to be an advantage of having more characters than states (Figure 6b). This suggests that base-editors may be a promising future direction for lineage tracing from a theoretical perspective.

Another potentially promising design consideration concerns the range of character mutation rates and their variability across different target sites – a parameter that can be precisely engineered [30]. In this design, one would expect the variability to help distinguish between early and late branching points and consequently achieve better resolution of the underlying phylogeny [51, 31, 32]. We simulated “Phased Recorders” (Figure S22) with varying levels of target-site cutting variability and observe that this design allows for better inference when the distributions of mutation probabilities are more dispersed (Figure S22b). This becomes particularly useful when one can integrate accurate indel priors into Cassiopeia.

Overall, these results serve to illustrate how Cassiopeia and the simulation framework can be used to explore experimental designs. While there inevitably will be challenges in new implementations, these analyses demonstrate theoretically how design parameters can be optimized for downstream tree inference. In this way, the combination of our algorithms and simulations enables others to explore not only new algorithmic approaches to phylogenetic reconstruction but also new experimental approaches for recording lineage information.

## Discussion

In this study, we have presented three resources supporting future single-cell lineage tracing technology development and applications. Firstly, we described Cassiopeia, a scalable and accurate maximum parsimony framework for inferring high-resolution phylogenies in single-cell lineage tracing experiments. Next, we introduced a simulation framework for benchmarking reconstruction methods and investigating novel experimental designs. Finally, we generated the largest and most diverse empirical lineage tracing experiment to date, which we present as a reference for the systematic evaluation of phylogeny inference on real lineage tracing data. With the combination of these three resources, we have demonstrated the improved scalability and accuracy of Cassiopeia over traditional approaches for single-cell lineage tracing data and have explored design principles for more accurate tracing. To ensure broad use, we have made a complete software package, including the algorithms, simulation framework, and a processing pipeline for raw data, all publicly available at www.github.com/YosefLab/Cassiopeia.

Though we illustrate that Cassiopeia provides the computational foundation necessary for future large-scale lineage tracing experiments, there are several opportunities for future improvement. Firstly, the inclusion of prior probabilities increases Cassiopeia’s performance only when parallel evolution is likely (e.g. with a high per-character mutation rate or in low character-state regimes). While maximum parsimony methods are attractive due to their non-parametric nature, future studies may build on our work here by developing more powerful approaches for integrating prior mutation rates into maximum likelihood [14, 41] or Bayesian inference [28] frameworks, perhaps relying on recent literature that seeks to predict indel formation probabilities [8, 2]. Secondly, confidence values for internal branches are not provided in this algorithm primarily due to the infeasibility of sampling enough trees from these large tree state-spaces (naturally, we hypothesize that this will also limit existing likelihood phylogeny approaches). Recent work adapting traditional bootstrapping to large trees [15]) or quantifying how well edges impose compatibility of characters [3] may be useful in determining confidence values. Thirdly, there exists a promising opportunity in developing new approaches for dealing with the amount of missing data. Determining a model which explicitly distinguishes between stochastic and heritable (e.g. from Cas9 cuts that remove the entire target site) missing data may increase tree accuracy. Alternatively, adapting supertree methods (such as the Triple MaxCut algorithm [46]) for lineage tracing data may be an interesting direction as they have been effective for dealing with missing data (but only when this missing data is randomly distributed [55]). Fourthly, while we provide theoretical and empirical evidence for our greedy heuristic, we note that there are opportunities for developing other heuristics - for example, by considering mutations in many characters rather than a single mutation as we do or using a distance-based heuristic.

The ultimate goal of using single-cell lineage tracers to create precise and quantitative cell fate maps will require sampling tens of thousands of cells (or more), possibly tracing over several months, and effectively inferring the resulting phylogenies. While recent studies [45] have highlighted the challenges in creating accurate CRISPR-recorders, our results suggest that with adequate technological components and computational approaches complex biological phenomena can be dissected with single-cell lineage tracing methods. Specifically, we show that Cassiopeia and the benchmarking resources presented here meet many of these challenges. Not only does Cassiopeia provide a scalable and accurate inference approach, but also our benchmarking resources enable the systematic exploration of stronger algorithms as well as more robust single-cell lineage tracing technologies. Taken together, this work forms the foundation for future efforts in building detailed cell fate maps in a variety of biological applications.

## Acknowledgments

The authors would like to thank the members of the Yosef and Weissman labs for their helpful discussions in the development of this project. For sequencing, the authors thank Eric Chow and the UCSF CAT for their help. This work was funded by NIH-NIAID grant U19 AI090023 (N.Y.), NIH R01 DA036858 and 1RM1 HG009490-01 (J.S.W.), F32 GM125247 (J.J.Q), and Chan-Zuckerberg Initiative 2018-184034. J.S.W. is a Howard Hughes Medical Institute Investigator. J.A.H. is the Rebecca Ridley Kry Fellow of the Damon Runyon Cancer Research Foundation (DRG-2262-16). M.M.C. is a Gordon and Betty Moore fellow of the Life Sciences Research Foundation.

## Author Contributions

M.G.J., A.K., J.J.Q, N.Y, and J.S.W. contributed to the design of the algorithm, interpretation of benchmarking results, and writing of the manuscript. A.K., C.X., and N.Y. conceived of the multi-state greedy algorithm and Steiner-Tree adaptation for the phylogeny inference problem. A.K. and M.G.J. implemented the algorithms and all code relevant to the project. M.G.J. and A.K. conducted all stress tests on synthetic datasets. R.W. and A.K. conducted experiments and theoretical work regarding the greedy heuristics robustness in lineage tracing experiments. J.J.Q. generated the *in vitro* reference dataset. M.G.J., J.J.Q, M.C, and J.A.H. designed the processing pipeline for empirical lineage tracing data. M.G.J. and J.J.Q processed the reference dataset and M.G.J. reconstructed trees.

## Author Information

Data will be deposited into GEO prior to publication. All software (including processing scripts) is available on our public github repository: www.github.com/YosefLab/Cassiopeia. The authors declare no competing interests.

## Methods

### *In vitro* lineage tracing experiment

#### Plasmid design and cloning

The Cas9-mCherry lentivector, PCTXX (to be added to Addgene), was designed for stable, constitutive expression of enzymatically active Cas9, driven by the viral SFFV promoter, insulated with a minimal universal chromatin opening element (minUCOE), and tagged with C-terminal, self-cleaving P2A-mCherry. PCTXX is derived from pMH0001 (Addgene Cat#85969, active Cas9) with the BFP tag exchanged with mCherry. The P2A-mCherry tag was PCR amplified from pHR-SFFV-KRAB-dCas9-P2A-mCherry (Addgene Cat #60954; forward: GAGCAACG-GCAGCAGCGGATCCGGAGCTACTAACTTCAG; reverse: ATATCAAGCTTGCATGCCTGCAGGTC-GACTTACTACTTGTACAGCTCGTCCATGC) and inserted using Gibson Assembly (NEB) into SbfI/BamHI-digested pMH0001 (active Cas9). Resulting plasmid was used for lentiviral production as described below.

The Target Site lentivector, PCT48 (to be added to Addgene), was derived from the reverse lentivector PCT5 (to be added to Addgene) containing GFP driven by the EF1a promoter. The sequence of the 10X amplicon with most common polyA location is the following:

> AATCCAGCTAGCTGTGCAGCNNNNNNNNNNNNNNATTCAACTGCAGTAATGCTACCTCGTACTCACGCTTTCCAAGTGCTTGGCGTCGCATCTCGGTCCTTTGTACGCCGAAAAATGGCCTGACAACTAAGCTACGGCACGCTGCCATGTTGGGTCATAACGATATCTCTGGTTCATCCGTGACCGAACATGTCATGGAGTAGCAGGAGCTATTAATTCGCGGAGGACAATGCGGTTCGTAGTCACTGTCTTCCGCAATCGTCCATCGCTCCTGCAGGTGGCCTAGAGGGCCCGTTTAAACCCGCTGATCAGCCTCGACTGTGCCTTCTAGTTGCCAGCCATCTGTTGTTTGCCCCTCCCCCGTGCCTTCCTTGACCCTGGAAGGTGCCACTCCCACTGTCCTTTCCTAATAAAAAAAAAAAAAAAAAAAAAAA

where *N* denotes our 14bp random integration barcode. PCT5 was digested with SfiI and EcoRI within the 3’UTR of GFP. The Target Site sequence was ordered as a DNA fragment (gBlock, IDT DNA) containing three Cas9 cut-sites and a high diversity, 14-basepair randomer (integration barcode, or intBC). The fragment was PCR amplified with primers containing Gibson assembly arms compatible with SfiI/EcoRI-digested PCT5 (forward: GATGAGCTCTACAAATAATTAAT-TAAGAATTCGTCACGAATCCAGCTAGCTGT; reverse: GGTTTAAACGGGCCCTCTAGGC-CACCTGCAGGAGCGATGG). The amplified Target Site fragment was inserted into the digested PCT5 backbone using Gibson Assembly. The assembled lentivector library was transformed into MegaX competent bacterial cells (Thermo Fisher) and grown in 1L of LB with carbenicillin at 100 *µ*g/mL. Lentivector plasmid was recovered and purified by GigaPrep (Qiagen), and used for high-diversity lentiviral production as described below.

The triple-sgRNA-BFP-PuroR lentivector, PCT61 (to be added to Addgene), is derived from pBA392 (to be added to Addgene) as previously described [1, 29] containing three sgRNA cassettes driven by distinct U6 promoters and constitutive BFP and puromycin-resistance markers for selection. Importantly, the three PCT61 sgRNAs are complementary to the three cut-sites in the PCT48 Target Site. To slow the cutting kinetics of the sgRNAs to best match the timescale involved in the *in vitro* lineage tracing experiments [7], the sgRNAs contain precise single-basepair mismatches that decrease their avidity for the cognate cut-sites [21]. The triple-sgRNA lentivector was cloned using four-way Gibson assembly as described in [29]. Resulting plasmid was used for lentiviral production as described below.

#### Cell culture, DNA transfections, viral preparation, and cell line engineering

A549 cells (human lung adenocarcinoma line, ATCC CCL-185) and HEK293T were maintained in Dulbecco’s modified eagle medium (DMEM, Gibco) supplemented with 10% FBS (VWR Life Science Seradigm), 2 mM glutamine, 100 units/mL penicillin, and 100 *µ*g/mL streptomycin. Lentivirus was produced by transfecting HEK293T cells with standard packaging vectors and TransIT-LTI transfection reagent (Mirus) as described in ([1]). Target Site (PCT48) lentiviral preparations were concentrated 10-fold using Lenti-X Concentrator (Takara Bio). Viral preparations were frozen prior to infection. Triple-sgRNA lentiviral preparations were titered and diluted to a concentration to yield approximately 50% infection rate.

To construct the lineage tracing-competent cell line, A549 cells were transduced by serial lentiviral infection with the three lineage tracing components: (1) Cas9, (2) Target Site, and (3) triple-sgRNAs. First, A549 cells were transduced by Cas9 (mCherry) lentivirus and mCherry+ cells were selected to purity by fluorescence-activated cell sorting on the BD FACS Aria II. Second, A549-Cas9 cells were transduced by concentrated Target Site (GFP) lentivirus and GFP+ cells were selected by FACS; after sorting, Target Site infection and sorting were repeated two more times for a total of three serial lentiviral transfections, sorting for cells with progressively higher GFP signal after each infection. This strategy of serial transfection with concentrated lentivirus yielded cells with high copy numbers of the Target Site, which were confirmed by quantitative PCR. Third, A549 cells with Cas9 and Target Site were transduced by titered triple-sgRNA (BFP-PuroR) lentivirus and selected as described below.

#### *In vitro* lineage tracing experiment, single-cell RNA-seq library preparation, and sequencing

One day following triple-sgRNA infection, cells were trypsinized to a single-cell suspension and counted using an Accuri cytometer (BD Biosciences). Approximately 25 cells were plated in a single well of a 96-well plate. Seven days post-infection, cells were trypsinized and split evenly into two wells of a 96-well plate. Cells stably transduced by triple-sgRNA lentivirus were selected by adding puromycin at 1.5 *µ*g/mL on days 9 and 11 post-infection; puromycin-killed cells were removed by washing the plate with fresh medium. After 14 days, cells were trypsinized and split evenly for a second time into four wells of a 6-well plate. Finally, after 21 days in total, cells from the four wells were trypisinized to a single-cell suspension and collected.

Cells were washed with PBS with 0.04% w/v bovine serum albumin (BSA, New England Bio-labs), filtered through 40 *µ*m FlowMi filter tips filter tips (Bel-Art), and counted according to the 10x Genomics protocol. Approximately 14,000 cells per sample were loaded (expected yield: approximately 10,000 cells per sample) into the 10x Genomics Chromium Single Cell 3’ Library and Gel Bead Kit v2, and cDNA was reverse-transcribed, amplified, and purified according to the manufacturer’s protocol. Resulting cDNA libraries were quantified by BioAnalyzer, yielding the expected size distribution described in the manufacturer’s protocol.

To prepare the Target Site amplicon sequencing library, resulting amplified cDNA libraries were further amplified with custom, Target Site-specific primers containing P5/P7 Illumina adapters and sample indices (forward: CAAGCAGAAGACGGCATACGAGATXXXXXXXXGTCTCGTGGGCTCG-GAGATGTGTATAAGAGACAGAATCCAGCTAGCTGTGCAGC; reverse: CAAGCAGAAGACG-GCATACGAGATXXXXXXXXGTCTCGTGGGCTCGGAGATGTGTATAAGAGACAGGCATG-GACGAGCTGTACAAGT; “X” denotes sample indices). PCR amplification was performed using Kapa HiFi HotStart ReadyMix, as in [1], according to the following program: melting at 95°C for 3 minutes, then 14 cycles at 98°C for 15 seconds and 70°C for 20 seconds. Approximately 12 fmol of template cDNA were used per reaction; amplification was performed in quadruplicate to avoid PCR-induced library biases, such as jack-potting. PCR products were re-pooled and purified by SPRI bead selection at 0.9x ratio and quantified by BioAnalyzer.

Target Site amplicon libraries were sequenced on the Illumina NovaSeq S2 platform. Due to the low sequence complexity for the Target Site library, a phiX genomic DNA library was spiked in at approximately 50% for increased sequence diversity. The 10x cell barcode and unique molecular identifier (UMI) sequences were read first (R1: 26 cycles) and the Target Site sequence was read second (R2: 300 cycles); sample identities were read as indices (I1 and I2: 8 cycles, each). Over 550M sequencing clusters passed filter and were processed as described below.

### Processing Pipeline

#### Read Processing

Each target site was sequenced using the Illumina Nova-seq platform, producing 300bp long-read sequences. The Fastq’s obtained were quantitated using 10x’s cellranger suite, which simultaneously corrects cell barcodes by comparing against a whitelist of 10x’s approved cell barcodes.

For each cell, a consensus sequence for each unique molecule identifier (UMI) was produced by collapsing similar sequences, defined by those sequences differing by at most one Levenshtein distance. A directed graph is constructed, where sequences with identical UMI’s are connected to one another if the sequences themselves differ by at most one Levenshtein distance. Then, UMI’s in this network are collapsed onto UMI’s that have greater than or equal number of reads. This produces a collection of sequences indexed by the cell barcode and UMI information (i.e. there is a unique sequence associated with each UMI).

Before aligning all sequences to the reference, preliminary quality control is performed. Specifically, in cases where UMI’s in a given cell still have not been assigned a consensus sequence, the sequence with the greatest number of reads is chosen. Cells are then filtered based on the number of reads and UMIs observed, and finally a filtered file in Fastq format returned.

#### Allele Calling

Alignment is performed with Emboss’s Water local alignment algorithm. Optimal parameters were found by performing a grid search of gap open and gap extend parameters on a set of 1,000 simulated sequences, comparing a global and local alignment strategy. We found a gap open penalty of 20.0 and a gap extension penalty of 1.0 produced optimal alignments. The “indels” (insertions and deletions resulting from the Cas9 induced double-strand break) at each cut site in the sequences are obtained by parsing the cigar string from the alignments. To resolve possible redundancies in indels resulting from Cas9 cutting, the 5’ and 3’ flanking 5-nucleotide context is reported for each indel.

#### UMI Error Correction

To correct errors in the UMI sequence either introduced during sequencing, PCR preparation, or data processing, we leverage the allele information. UMIs are corrected within groups of identical cell barcode-integration barcode pairs (i.e. we assume that only UMIs encoding for the same intBC in a given cell can be corrected). We reason that ideally, for a given integration barcodes, a cell will only report one sequence, or allele. Within these “equivalence classes,” UMIs that differ by at most 1 Levenshtein distance (although this number can be user-defined) are corrected towards the UMI with a greater number of reads.

#### Cell-based Filtering

With the UMI corrected and indels calculated, the new “molecule table” is subjected to further quality control. Specifically, UMIs are filtered based on the number of reads, integration barcodes (denoting a particular integration site) are error corrected based on a minimum hamming distance and identical indels (referred to as alleles), and in the case where multiple alleles are associated with a given integration barcode a single allele is chosen based on the number of UMIs associated with it.

#### Calling Independent Clones

Collections of cells part of the same clonal population, are identified by the set of integration barcodes each cell contains. Because all cells in the same clone are clonal, we reasoned that cells in the same clone should all share the same set of integration barcodes that the progenitor cell contained. Because of both technical artifacts (e.g. sequencing errors, PCR amplification errors) and biological artifacts (e.g. bursty expression, silenced regions) however, rather than looking for sets of non-overlapping sets, we perform an iterative clustering procedure. We begin by selecting the intBC that is shared amongst the most cells and assign any cell that contains this barcode to a cluster and remove these cells from the pool of unassigned cells. We perform this iteratively until at most *k* percent (in our case defined as .5% of cells are unassigned, which we assign to a “junk” clone.

Using the set of integration barcodes for each clone, we are able to identify doublets that consist of cells from different clones. Finally, after identifying doublets, to further filter out low quality integration barcodes, for each clone integration barcodes that are not shared by at least 10% of cells in a given clone are filtered out, producing the final allele table.

#### Filtering of clones for Reconstruction

We filtered out clones upon two criteria: firstly, we removed clone 1 as we deduced that it had two defective guides; secondly, we removed lineages that reported fewer than 10% unique cells (thus removing clone 7). The remainder of clones were reconstructed.

#### Estimation of Per Character Mutation Rates

To estimate mutation rates per clone, we assume that every target site was mutated at the same rate and independently of one another across 15 generations. Assuming some mutation rate, *p*, per character, we know that the probability of not observing a mutation in *d* generations is (1 − *p*)^*d*^ in a given character and that the probability of observing at least 1 mutation in that character is 1 − (1− *p*)^*d*^. Then, giving this probability 1 − (1− *p*)^*d*^ = *m* can be used as a probability of observing a mutated character in a cell and model the number of times a character appears mutated in a cell as a binomial distribution where the expectation is simply *nm* where *n* is the number of characters. Said simply, given this model, one would expect to see *nm* characters mutated in a cell). In this case, the empirical expectation is the mean number of times a given character appeared mutated in a cell (averaged across all cells), which we denote as *K* and propose that

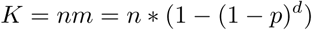

and thus *p*, the mutation rate, is

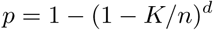

### Bulk Cutting Experiment to Determine Prior Probabilities of Indel Formation

Two and four days following triple-sgRNA (PCT61) infection, infected cells were selected by adding puromycin at 1.5 *µ*g/mL; puromycin-killed cells were removed by washing the plate with fresh medium. Cells were split every other day, and 500k cells were collected on days 7, 14, and 28. Frozen cell pellets were lysed and the genomic DNA was extracted and purified by ethanol precipitation. The PCT48 Target Site locus was PCR amplified from genomic DNA samples (forward: TCGTCGGCAGCGTCAGATGTGTATAAGAGACAGAATCCAGCTAGCT-GTGCAGC; reverse: GTCTCGTGGGCTCGGAGATGTGTATAAGAGACAGTCGAGGCTGATCAGCG) and further amplified to incorporate Illumina adapters and sample indices (forward: AATGAT-ACGGCGACCACCGAGATCTACACXXXXXXXXTCGTCGGCAGCGTCAG; reverse: CAAGCA-GAAGACGGCATACGAGATXXXXXXXXGTCTCGTGGGCTCGGAG; “X” denotes sample indices). The subsequent amplicon libraries were sequenced on an Illumina MiSeq (paired end, 300 cycles each). Sequencing data was analyzed as described below.

### Determining Prior Probabilities of Indel Formation

To determine the prior probabilities of edits, we leverage the fact that we have access to a large set of target sites (or intBCs) with a similar sequence (apart from the random barcode at the 5’ end); namely, a total of 117 intBC across the 11 clones. To compute the prior probability for a given indel, we compute the empirical frequency of observing this mutation out of all unique edits observed. Specifically, we compute the prior probability of a given indel *s, q*_*s*_ as the following:

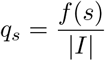

where *f* (*s*) is the number of intBC’s that had *s* in at least one cell and |*I*| is the number of intBCs that are present in the dataset.

As further support for this method, we used the bulk experiment consisting of many separately engineered A549 cells, as described in the previous section. The advantage of the bulk experiment is that we have access to substantially more intBCs (> 10*k*), thus providing a more robust estimation of *q*_*s*_. We therefore employed the same approach to estimate indel formation rates from the bulk data and find that the resulting rates correlate well with the indel rates estimated from the single cell lineage tracing experiment (Figure S18).

### Doublet Detection

#### Methods to Detect Doublets

We hypothesized that doublets could come in two forms and that we could use various components of the intBC data structure to identify them. Namely, doublets could be of cells from the identical clone, here dubbed “intra-doublets”, or doublets could be of cells from separate clones, here dubbed “inter-doublets.”

In the case of “intra-doublets”, we can utilize the fact that these cells will have a large overlap in their set of intBCs but will report “conflicting” alleles for each of these intBCs. Thus, to identify these doublets, we calculate the percentage of UMIs that are conflicting in each cell. Explicitly, for each cell we iterate over all intBCs and sum up the number of UMIs that correspond to an allele that conflicts with the more abundant allele for a given intBC; we then use the percentage of these UMIs to identify doublets. We perform this after all UMI and intBC correction in hopes of calling legitimate conflicts.

To deal with “inter-doublets”, we developed a classifier that leverages the fact that cells from different clones should have non-overlapping intBC sets. While this is the ideal scenario, often times intBCs are shared between clones for one of two reasons (1) the clustering assignments are noisy or (2) the transfections of intBCs resulted in two cells receiving the same intBC, even though cells are supposed to be progenitors of separate clones. Our strategy is thus: for each cell *c*_*i*_ ∈ *C* calculate a “membership statistic”, *m*_*i,k*_ for each clone *l*_*k*_ ∈ *L*. The membership statistic is defined as so:

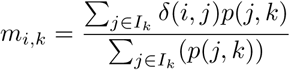

where *I*_*k*_ is the set of intBCs for the clone *l*_*k*_ and *p*(*j, k*) is the prevalence rate of the intBC *j* in *l*_*k*_. We use *δ*(*i, j*) as an indicator function for whether or not we observed the intBC *j* in the cell *c*_*i*_. Intuitively, this membership statistic is a weighted similarity for how well the cell fits into each clone, where we are weighting by how much we are able to trust the intBC that is observed in the cell. To put all on the same scale, we normalize by total membership per cell, resulting in our final statistic, 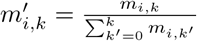 We then filter out doublets whose *m′* for their classified clone falls below a certain threshold.

#### Simulation of Doublets

We simulated two datasets to test our methods for identifying doublets and to find the optimal criterion on which to filter out doublets. To test this strategy, we took a single clone from our final Allele Table (the table relating all cells and their UMIs to clones) and formed 200 doublets by combining the UMIs from two cells. We generated 20 of these datasets, and noted which cells were artificially introduced doublets.

Contrary to the strategy for simulating doublets from the same clone, we created artificial “inter” doublets from the final Allele Table by combining doublets from two different clones. Similarly, we generated 20 synthetic datasets each with 200 of these artificial doublets.

#### Identification of Decision Rule

To identify the optimal decision rule for calling both types of doublets, we tested decision rules ranging from 0 to 1.0 at 0.05 intervals and calculated the precision and recall at each of these rules. Taking these results altogether, we provide an optimal decision rule where the F-measure (or the weighted harmonic mean of the precision and recall) of these tests is maximal.

### Algorithmic Approaches For Phylogenetic Reconstruction

One way to approach the phylogenetic inference problem is to view each target site as a “character” that can take on many different possible “states” (each state corresponding to an indel pattern induced by a CRISPR/Cas9 edit at the target site). Formally, these observations can be summarized in a “character matrix”, *M* ∈ *R*^*n,m*^, which relates the *n* cells by a set of characters *χ* = {*χ*_1_, …, *χ*_*m*_} where each character *χ*_*i*_ can take on some *k*_*i*_ possible states. Here, each sample, or cell, can be described as a concatenation of all of their states over characters in a “character string”. From this character matrix, the goal is to infer a tree (or phylogeny), where leaf nodes represent the observed cells, internal nodes represent ancestral cells, and edges represent a mutation event.

We first propose an adaption of a slow, but accurate, Steiner-Tree algorithm via Integer Lineage Programming (ILP) to the lineage tracing phylogeny problem. Then, we propose a fast, heuristic-based greedy algorithm which simultaneously draws motivation from classical perfect phylogeny algorithms, and the fact that mutations can only occur unidirectionaly from the unmutated, or *s*_0_ state. Lastly, we combine these two methods and present a hybrid method, which presents better results than our greedy approach, yet remains feasible to run over tens of thousands of cells.

#### Adaptation to Steiner Tree Problem

Steiner Trees are a general problem for solving for the minimum weight tree connecting a set of target nodes. For example, if given a graph *G* = (*V, E*) over some *V* vertices and *E* edges, finding the Steiner-Tree over all *v* ∈ *V* would amount to solving for the minimum spanning tree (MST) of *G*. While there exist polynomial time algorithms for the minimum-spanning tree, the general Steiner Tree problem, where the set of targets *T* ⊆ *V* is designated, is NP-hard.

Previously, Steiner-Trees have been suggested to solve for the maximum parsimony solution to the phylogeny problem. Here, the graph would consist of all possible cells (both observed and unobserved) and each edge would consist of a possible evolutionary event connecting two states (e.g. a mutation). Generally, given a set of length-*l* binary “character-strings” (recall that these are the concatenation of all character states for a given sample), we can solve for the maximum parsimony solution by finding the optimal Steiner Tree over the 2^*l*^ hypercube (i.e. graph). As a result, by converting our multi-state characters to binary characters via one hot encoding, theoretically, we should be able to compute the most parsimonious tree which best explains the observed data. However, in practice this method turns out to be infeasible, as we deal with hypercubes of size *O*(2^*mn*^), where *m* is the number of characters, and *n* is the number of states. In the following, we will propose a method for estimating the underlying search space, providing us with a feasible solvable instance and a formulation of an Integer-Linear Programming (ILP) problem to solve for the optimal Steiner-Tree.

##### Approximation of Potential Graph

We first begin by constructing a directed acyclic graph (DAG) *G*, where nodes represent cells. We then take the source nodes, or nodes with in-degree 0, of *G*, and for each pair of source nodes, consider the latest common ancestor (LCA) they could have had. This LCA has an unmutated state for character *χ*_*i*_ if they disagree across two source nodes, and the same state as the two source nodes if they agree in value. If the edit distance between these two cells is below a certain threshold *d*, we add the LCA to *G*, along with directed edges to the two source nodes, weighted by the edit distance between the parent and the source. We repeat this process until only one node remains as a source: the root.

One may think that this step explodes with *O*(*n*^2^) complexity at each stage, where n is the number of source nodes in each prior stage, as we consider all pairs of source nodes. However, we note that the number of mutations per latest common ancestor is always less than both children, and therefore, we eventually converge to the root. Therefore, when dealing with several hundred cells, the potential graph is feasible to calculate.

Furthermore, to add scalability to the approximation of the Potential Graph, we allow the user to provide a “maximum neighborhood size” which will be used to dynamically solve for the optimal LCA distance threshold *d* to use. One may think of this as the maximum memory or time allowed for optimizing a particular problem. Since the size of the Potential Graph can grow quite large in regards to the number of nodes, we iteratively create potential graphs for various threshold *d* and at each step ensure that the number of nodes in the network does not exceed the maximum neighborhood size provided. If at any point the number of nodes does exceed this maximum size, we return the potential graph inferred for an LCA threshold of *d* − 1.

##### Formulation of Integer Linear Programming Problem

Given our initial cells, *S*, the underlying potential graph drawn from such cells, *G*, and the final source node, or root, *r* from *G*, we are interested in solving for *𝒯* = *SteinerTree*(*r, S, G*). We apply an integer linear programming (ILP) formulation of Steiner Tree, formulated in terms of network flows, with each demand being met by a flow from source to target. Below we present the Integer Linear Programming formulation for Steiner Tree. We use Gurobi [23], a standard ILP solver package

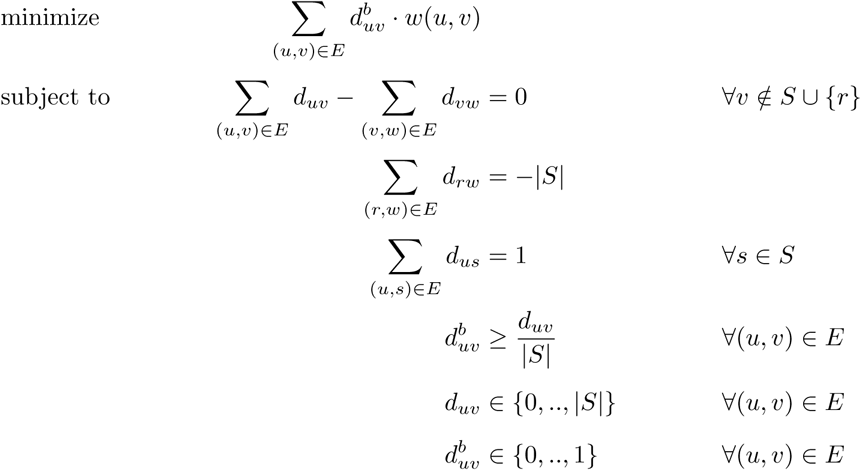

Each variable *d*_*uv*_ denotes the flow through edge (*u, v*), if it exists; each variable 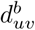 denotes whether (*u, v*) is ultimately in the chosen solution sub-graph. The first constraint enforces flow conservation, and hence that the demands are satisfied, at all nodes and all conditions. The second constraint requires |*S*| units of flow come out from the *root*. The third constraint requires that each target absorb exactly one unit of flow. The fourth constraint ensures that if an edge is used at any condition, it is chosen as part of the solution.

Below we explicitly define the algorithm in pseudocode.

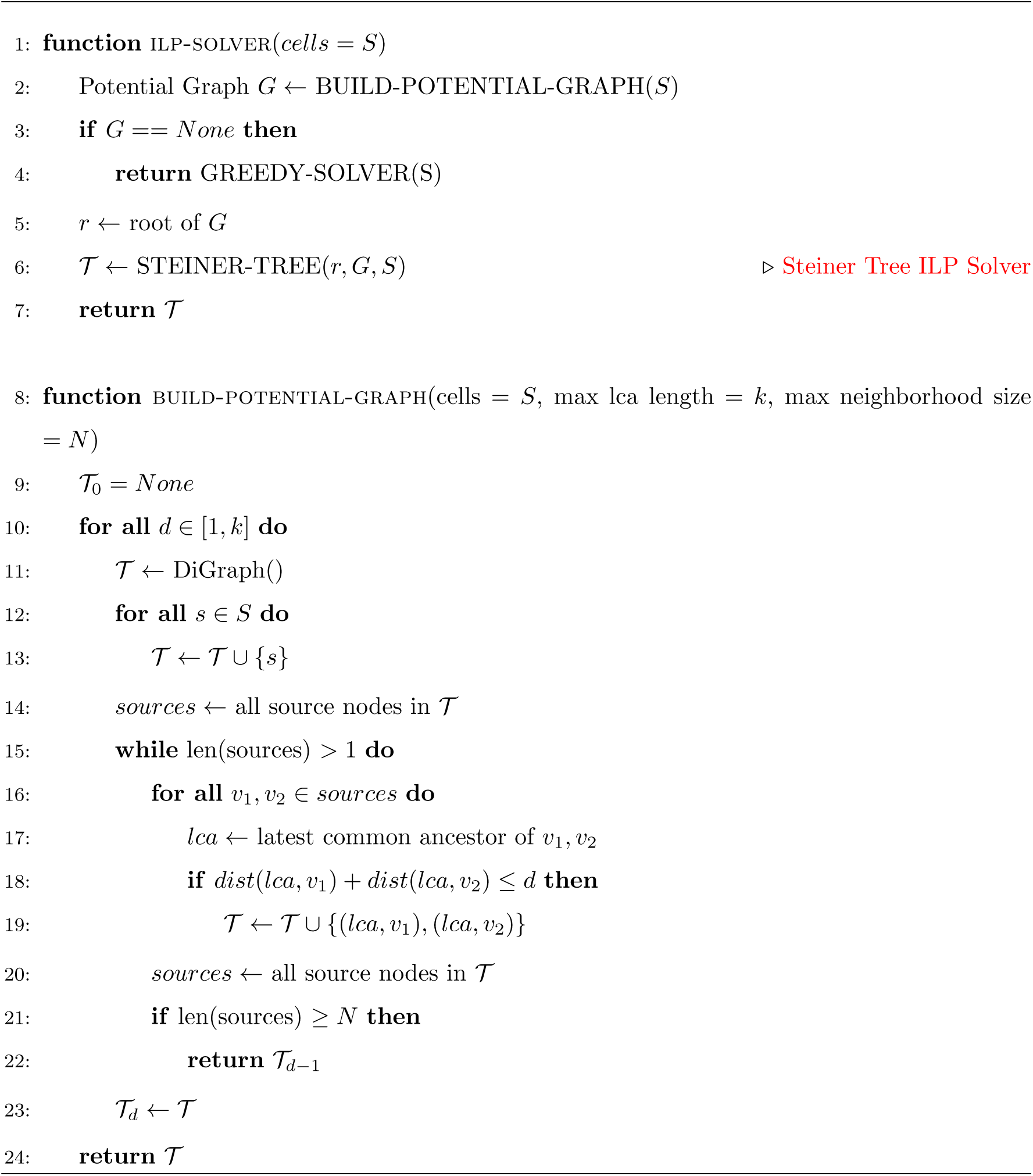

#### Stability Analysis of the Maximum Neighborhood Size Parameter

To evaluate the stability of the user-defined maximum neighborhood size parameter, we assessed the accuracy of the reconstructions for parameters varying from 800 to 14, 000. We used trees simulated under default conditions (400 samples, 40 characters, 40 states per character, 11 generations, 2.5% mutation rate per character, and a mean dropout rate of 17%). The accuracy of trees were compared to the tree generated with a parameter of 14, 000 using the triplets correct statistic. We used 10 replicates to provide a sense for how stable a given accuracy is.

In addition to providing measures of accuracy, we also provide the optimal LCA threshold *d* found for a given maximum neighborhood size during the inference of these potential graphs. Using these analysis, we found that a maximum neighborhood size of 10, 000 nodes seemed to be an ideal tradeoff between scalability and accuracy (as it is in the regime where accuracy saturates) for our default simulations. This corresponded to a mean LCA threshold, *d*, of approximately 5.

#### Heuristic-Based Greedy Method

##### On Perfect Phylogeny & Single Cell Lineage Tracing

In the simplest case of phylogenetics, each character is binary (i.e. *k*_*i*_ = 2, ∀*i* ∈ *m*) and can mutate at most once. This case is known as “perfect phylogeny” and there exist algorithms (e.g. a greedy algorithm by Dan Gusfield [24]) for identifying if a perfect phylogeny exists over such cells, and if so find one efficiently in time *O*(*mn*), where *m* is the number of characters and *n* are the number of cells. However, several limitations exist with methods such as Gusfield’s algorithm. One potential problem in using existing greedy perfect phylogeny algorithms for lineage tracing is that they require the characters to be binary. Indeed, if the characters are allowed to take any arbitrary number of states, the perfect phylogeny problem becomes NP-hard. However, while the number of states (CRISPR/Cas9-induced indels at a certain target site) in lineage tracing data can be large, these data benefit from an additional restriction that makes it more amenable for analysis with a greedy algorithm. Below, we show that because the founder cell (root of the phylogeny) is unedited (i.e. includes only uncut target sites) and that the mutational process is irreversible, we are able to theoretically reduce the multi-state instance (as observed in lineage tracing) to a binary one so that it can be resolved using a greedy algorithm.

A second remaining problem in using these perfect phylogeny approaches is that we cannot necessarily expect every mutation to occur exactly once. In theory, it may happen that the same indel pattern is induced in exactly the same target site on two separate occasions throughout a lineage tracing experiment, especially if a large number of cell cycles takes place. A final complicating factor is that these existing greedy algorithms often assume that all character-states are known, whereas lineage tracing data is generated by single-cell sequencing, which often suffers from limited sensitivity and an abundance of “dropout” (stochastic missing data) events.

##### The Greedy Algorithm

We suggest a simple heuristic for a greedy method to solve the maximum parsimony phylogeny problem, motivated by the classical solution to the perfect phylogeny problem and irreversibility of mutation. Namely, we consider the following method for building the phylogeny: Given a set of cells, build a tree top-down by splitting the cells into two subsets over the most frequent mutation. Repeat this process recursively on both subsets until only one sample remains.

Formally, we choose to split the dataset into two subsets, *O*_*i,j*_ and *Ō*_*i,j*_, such that *O*_*i,j*_ contains cells carrying mutation *s*_*j*_ in *χ*_*i*_, and *Ō*_*i,j*_ contains cells without *s*_*j*_ in *χ*_*i*_. We choose *i, j* based on the following criteria:

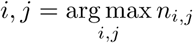

where *n*_*i,j*_ is the number of cells that carry mutation *s*_*j*_ in character *χ*_*i*_. We continue this process recursively until only one sample exists in each subset. We note that this method operates over cells with non-binary states, solving the first of problems addressed earlier.

A major caveat exists with methods such as the greedy method proposed by Gusfield, as well as the one proposed by us thus far: namely, they assume all character states are known (i.e. no dropout). However, in our practice, we often encounter dropout as a consequence of Cas9 cutting or stochastic, technical dropout due to the droplet-based scRNA-seq platform. To address this problem in our greedy approach, during the split stage, these cells are not initially assigned to either of the two subsets, *O*_*i,j*_ or *Ō*_*i,j*_. Instead, for each individual sample which contains a dropped out value for chosen split character *χ*_*i*_, we calculate the average percentage of mutated states shared with all other cells in *O*_*i,j*_ and *Ō*_*i,j*_ respectively, and assign the sample to the subset with greater average value.

Appending the dropout resolution stage with the initial split stage, we present our greedy algorithm below in its entirety.

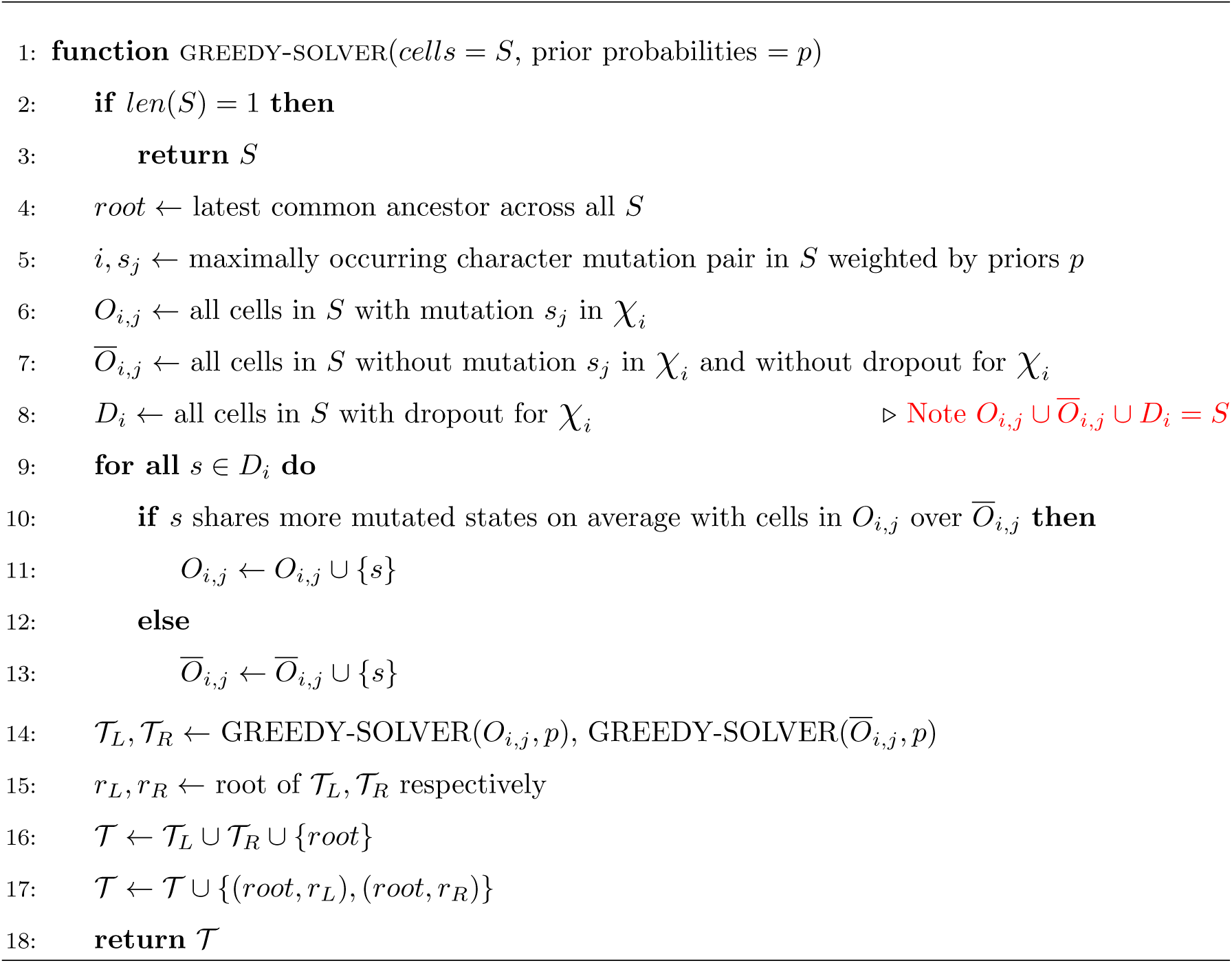

Overall, this method is very efficient, and scales well into tens of thousands of cells. Below, we show via proof below that this algorithm can find perfect phylogeny if one exists.

##### Cassiopeia-Greedy Algorithm Can Solve Multi-State Perfect Phylogeny

Here we show that while not required, Cassiopeia can solve the multi-state perfect phylogeny problem optimally. Importantly, however, Cassiopeia’s effectiveness makes no assumption about perfect phylogeny existing in the dataset but rather leverages this concept to provide a heuristic for scaling into larger datasets.

To show how Cassiopeia’s greedy method can solve perfect phylogeny optimally, we begin by introducing a few clarifying definitions prior to the main theorem. We define *M* as the original *n* cells by *n* character *k*-state matrix (i.e. entries ∈ {*s*_0_, …, *s*_*k*−1_}). We say *M* has a zero root perfect phylogeny if there exists a tree *𝒯* over its elements and character extensions such that the state of the root is all zeros and every character state are mutated into at most once. In addition, we assume that all non-leaf nodes of *𝒯* have at least two children (i.e. if they only have one child, collapse two nodes into one node). Finally, we offer a definition for *character compatibility*:

###### Definition 1.

*(Character Compatibility). For a pair of binary characters*, (*χ*_1_, *χ*_2_), *where the sets* (*O*_1_, *O*_2_) *contain the sets of cells mutated for χ*_1_ *and χ*_2_, *respectively, we say that they are compatible if one of the following is true:*

- *O*_1_ ⊆ *O*_2_
- *O*_2_ ⊆ *O*_1_
- *O*_1_ ∩ *O*_2_ = Ø

This definition extends to multi-state characters as well, assuming they can be binarized.

Before proving the main theorem, we first prove the following lemma:

###### Lemma 1.

*If M has a perfect phylogeny, then the most frequent character, mutation pair appears on an edge from the root to a direct child node.*

*Proof.* WLOG let *χ*_*i*_ : *s*_0_ → *s*_*j*_ denote the maximally occurring character, mutation pair within *M*. Suppose by contradiction that this mutation does not appear on an edge directly from root to a child, but rather on some edge (*u, v*) that is part of a sub-tree whose root *r*\*, is a direct child of the root. As *r*\* has at least two children, this implies that the mutation captured from the root to *r*\* must be shared by strictly more cells than *χ*_*i*_ : *s*_0_ → *s*_*j*_, thereby reaching a contradiction on *χ*_*i*_ : *s*_0_ → *s*_*j*_ being the maximally occurring mutation. □

###### Theorem 1.

*The greedy algorithm accurately constructs a perfect phylogeny over M if one exists.*

*Proof.* We approach via proof by induction. As a base case, a single is trivially a perfect phylogeny over itself.

Now suppose by induction that for up to *n* − 1 cells, if there exists a perfect phylogeny *T* over such cells, then the greedy algorithm correctly returns the perfect phylogeny. Consider the case of *n* cells. By the above lemma, we know we can separate these *n* cells into two subsets based on the most frequent character, mutation pair *χ*_*i*_ : *s*_0_ → *s*_*j*_, *O*_*i,j*_ and *Ō*_*i,j*_, where *O*_*i,j*_ contains cells with mutation *s*_*j*_ over *χ*_*i*_, and *Ō*_*i,j*_ = *M* − *O*_*i,j*_. By induction, the greedy algorithm correctly returns two perfect phylogenies over *O*_*i,j*_ and *Ō*_*i,j*_, which we can merge at the root, giving us a perfect phylogeny over n cells.

#### Accounting for Prior Probability of Mutations

In most situations, the probability of mutation to each distinct state may not be uniform (i.e. character *χ*_1_ mutating from the unmutated state *s*_0_ to state *s*_4_ may be twice as likely as mutating to state *s*_6_). Therefore, we incorporate this information into choosing which character and mutation to split over based on the following criteria:

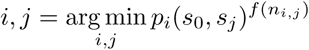

where *p*_*i*_(*s*_0_, *s*_*j*_) is the probability that character *χ*_*i*_ mutates from the unmutated state *s*_0_ to *s*_*j*_ and *f* (*n*_*i,j*_) is some transformation of the number of cells that report mutation *j* in character *i* that is supposed to reflect the future penalty (number of independent mutations of character *i* to state *j*) we will have to include in the tree if we do not pick *i, j* as our next split. After a comparison of 5 different transformations (Supp Figure 4), we find that *f* (*n*_*i,j*_) = *n*_*i,j*_ gives the best performance, leaving us with the following criteria for splittings:

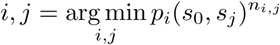

#### A Hybrid Method for Solving Single Cell Lineage Tracing Phylogenies

Due to the runtime constraints of the Steiner Tree Method, it is infeasible for such method to scale to tens of thousand of cells. Therefore, we build a simple hybrid method which takes advantage of the heuristic proposed in the greedy algorithm and the theoretical optimality of the Steiner Tree method.

Recall that in the greedy method, we continued to choose splits recursively until only one sample was left per subset. In this method, rather than follow the same process, we choose a cutoff for each subset (e.g. 200 cells). Once a subset has reached a size lower than said cutoff, we feed each individual subset into the Potential Graph Builder and Steiner Tree solver, which compute an optimal phylogeny for the subset of cells. After an optimal subtree is found, we merge it back into the greedy tree. Therefore, we build a graph whose initial mutations are chosen from the greedy method, and whose latter mutations are chosen more precisely via the Steiner Tree approach.

Below we present a pseudo-code algorithm for the hybrid method. We note the slight difference in greedy from before. Namely, greedy additionally accepts a cutoff parameter, and in addition to returning a network built up to that cutoff, returns all subsets that are still needed to be solved.

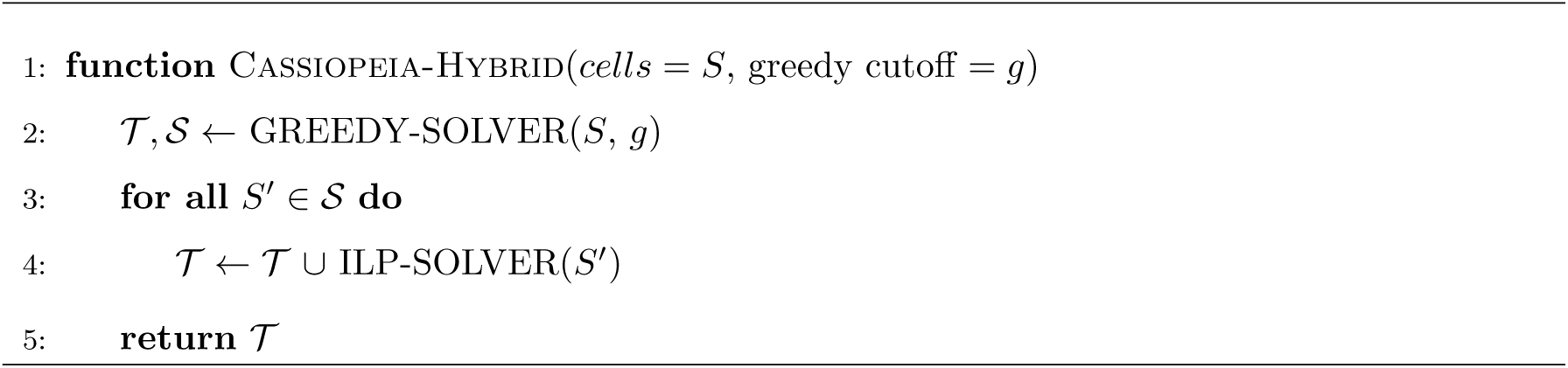

This approach scales well when each instance of Steiner Tree is ran on an individual thread, and thus often takes only a few hours to run on several thousand cells.

### Theoretical Analysis of Parallel Evolution

#### Estimating First and Second Moments of Double Mutations

##### Expected Number of Double Mutations

Under the framework of our simulation, we assume that each at each generation, every cell divides, and then each character of each cell undergoes random mutation independently. Let *p* be the probability that a particular character mutates, and *q* be the probability the character took on a particular mutated state given that it mutated. Let *T* be the true phylogenetic tree over the samples. According to our model, *T* must be a full binary tree, and the samples are leaves of *T*. Let *X* be the total number of times a particular mutation occurred in the *T*. Let *X*_*u,v*_ be an indicator variable for edge (*u, v*) such that:

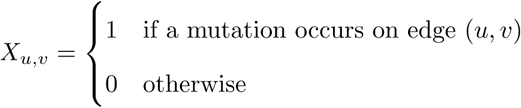

Let *h* be the height of the *T*, which is equalled to the number of generations. If *v* is at depth *d* in *T*, then the probability that a mutation occurs at (*u, v*) is *pq*(1 − *p*)^*d*−1^. Since there are 2^*d*^ nodes at depth *d*, we have:

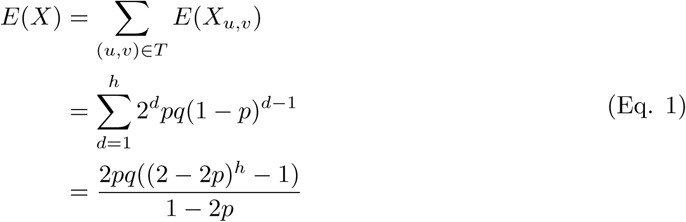

Let *n* = 2^*h*^ is the number of cells in our sample. If *p >* 0.5, *E*(*X*) ≤ 2*pq/*(2*p* −1), if *p* = 0.5, *E*(*X*) = 2*pqh* = *O*(log *n*), and if 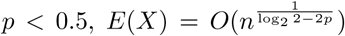. Moreover, for fixed *h, E*(*X*) has a single peak for *p* ∈ [0, 1], meaning that it increases with *p* for sufficiently small values of *p*, and always increases with *q*. Intuitively, this is because *E*(*X*) is small if 1) *p* is small enough that the character never mutates much throughout the experiment or 2) *p* is large enough that most mutations occur near the top of the tree, resulting in the extinction of unmutated cells early in the experiment. While *E*(*X*) peaks for values of *p* in between, it is always directly proportional to *q* because *X* is simply equalled to *q* time the number of times the character mutated.

##### Variance of Double Mutations

We can compute the variance as:

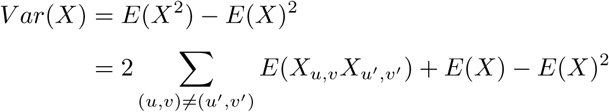

To compute *E*(*X*_*u,v*_*X*_*u*′,*v*′_), we note that for a given pair of edges (*u, v*) and (*u*′, *v*′), such that *LCA*(*u, u*′) is at depth *d, u* is at depth *d* + *l*, and *u*′ is at depth *l* + *k*, the probability that a mutation occurred on both edges is *p*^2^*q*^2^(1 − *p*)^*d*+*l*+*k*^. Thus, we have:

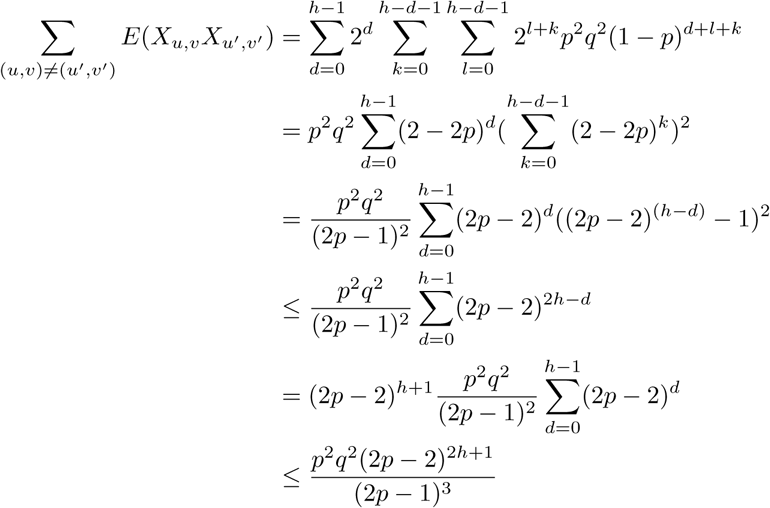

Thus, we can bound the variance as follows:

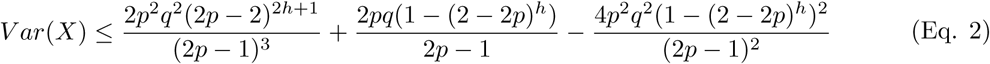

This means that in the case that *p >* 0.5:

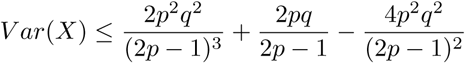

In the case that *p* = 0.5:

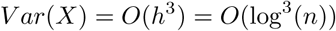

In the case that *p <* 0.5:

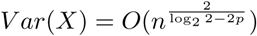

#### Least Squares Linear Estimate & Negative Correlation Between Frequency and Number of Double Mutations

To justify the greedy, we must show that if a mutation occurs frequently, then it is likely to have occurred less times throughout the experiment. Let *Y* be the frequency of a particular mutation in the samples. We estimate *X* given *Y* using the least squares linear estimate (LLSE) as follows:

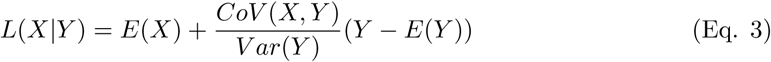

Since *CoV* (*X, Y*) = *E*(*XY*) − *E*(*X*)*E*(*Y*), we need only to compute *E*(*XY*), which we do by expressing *X* and *Y* in terms of the same indicators:

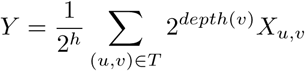

As a sanity check, it can easily be verified that *E*(*Y*) = *q*(1 − (1 − *p*)^*h*^) by computing *E*(*Y*) using these indicators:

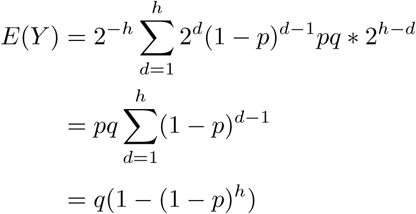

Thus, we can compute *E*(*XY*) similar to how we computed *E*(*X*^2^) for Variance.

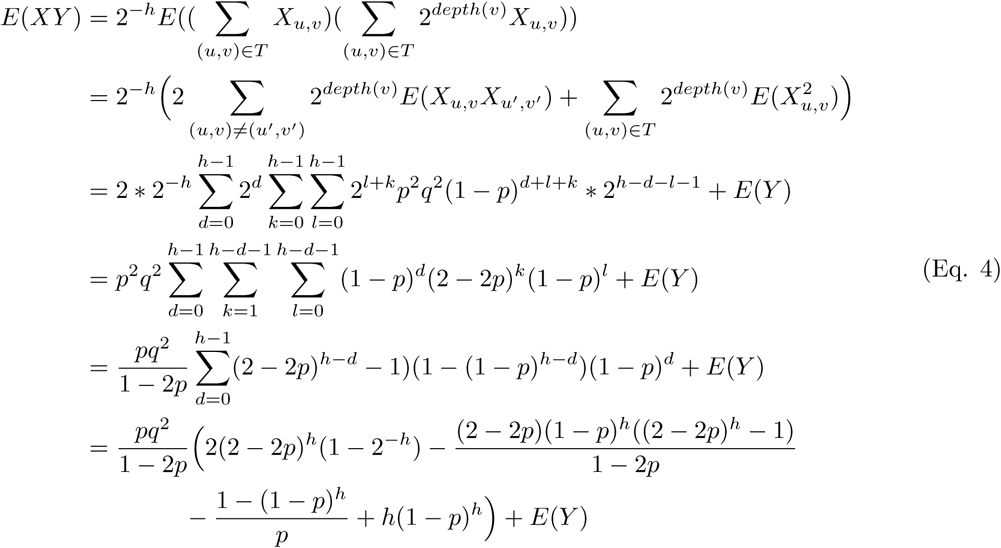

Assuming that is 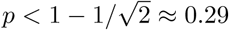 (based on our estimation of Cas9-cutting rates, this seems to be a biologically relevant probability), we have:

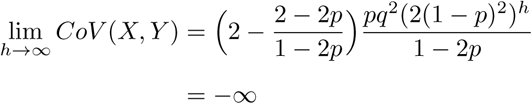

since 2 *<* (2 − 2*p*)*/*(1 − 2*p*) when *p <* 0.5.

*V ar*(*Y*) can be computed using the same indicators:

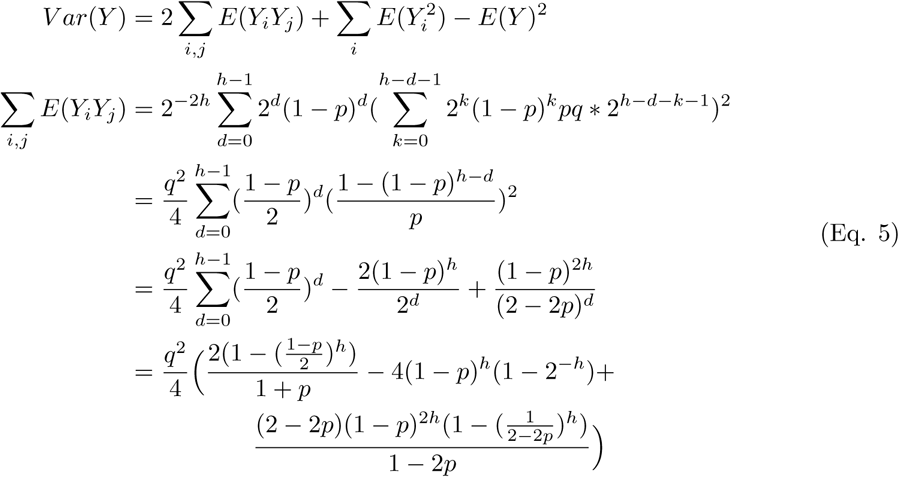

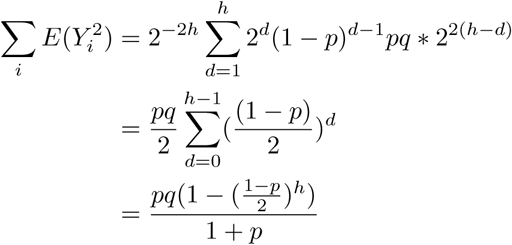

Note that if *p <* 0.5, every term in *V ar*(*Y*) converges to a constant as *h* → *∞*. Thus, if (1 − *p*)^2^ *>* 0.5, then as the depth increases, *X* and *Y* become exponentially more negatively correlated. This means that for biologically relevant values of *p*, the frequency of a mutation in the samples is negatively correlated with number of times the mutation occurred, thus justifying the rationale of splitting the sample on more frequently occurring mutations.

#### Simulation For Tracking the Evolution of a Particular Mutation

To more efficiently simulate the number of occurrences of a particular mutation, we define {*N*_1_, *N*_2_, …*N*_*h*_} as a Markov chain, where *N*_*t*_ is the number of unmutated cells at generation *t*, and *N*_1_ = 1. Let *A*_*t*_ ~ *Bin*(2*N*_*t*_, *p*) be the number of cells that mutates at generation *t*, and *B*_*t*_ ~ *Bin*(*A*_*t*_, *a*) be the number of mutated cell that took on the particular state in question. The Markov chain evolves as *N*_*t*+1_ = 2*N*_*t*_ − *A*_*t*_. Note that we assume, in this model, that mutation can only occur after cell division. Thus we have 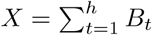 and 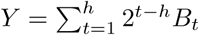.

### Assessing the Precision of Greedy Splits

To assess the precision of greedy splits, we first simulated 100 true phylogenies of 400 cells (without dropout) for all pairs of parameters in *num_states* = {2, 10, 40} and *p*_*c*_*ut* = {0.025, 0.1, 0.4}. For each network, we assessed the precision of the greedy split as follows:

1. We used the criteria *i, j* = arg max*i,j n*_*i,j*_ to select the character *χ*_*i*_ and state *j* to split on (as Cassiopeia-Greedy would do). This group of cells that have a mutation *j* in character *χ*_*i*_ is called *G*.
2. For define the a set of *n* subsets corresponding to cells that inherited the (character, state) pair (*i, j*) independently using the true phylogenies, and call this set *S* = (*s*_1_, *s*_2_, …, *s*_*n*_) (this corresponds to there being *n* parallel evolution events for the (character, state) pair (*i, j*).
3. We presume that the largest group of cells in *S* is the “true positive” set (let this be defined as *s*′ = arg max*s* |*s*_*i*_|. We then define the precision *P* as the proportion of true positives in the set *G* – i.e. 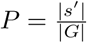.

### Statistics for IVLT Analysis

#### Meta Purity Statistic

To calculate the agreement between clades (i.e. the leaves below a certain internal node of the tree) and some meta-value, such as the experimental plate from which a sample came from, we can employ a Chi-Squared test. Specifically, we can compute the following statistic: considering some *M* clades at an arbitrary depth *d*, we find the count of meta values associated with each leaf in each clade, resulting in a vector of values *m*_*i*_ comprised of these meta-counts for each clade *i*. We can form a contingency table summarizing these results, *T*, where each internal value is exactly *m*_*i,j*_ - the counts of the meta item *j* in clade *i*. A Chi-Squared test statistic can be computed from this table.

To compare across different trees solved with different methods, we report the Chi-Squared Test Statistic as a function of the number of clades, or degrees of freedom of the test.

#### Mean Majority Vote Statistic

The Mean Majority Vote statistic seeks to quantify how coherent each clade is with respect to its majority vote sample at a give depth. For a given clade with leaves *L*_*i*_ where |*L*_*i*_| = *n*, where every leaf *l*_*i,j*_ corresponds to cell *j* in clade *i* has some meta label *m*_*j*_, the majority vote of the clade is *v* = *argmax*_*m*′∈*M*_ Σ_*j*∈*n*_ *δ*(*j, m*′). Here *M* is the full set of possible meta values and *δ*(*m*_*j*_, *m*′) is an indicator function evaluating to 1 iff *m*_*j*_ = *m*′. The membership of this clade is then simply 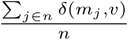. Then, the mean membership is the mean of these membership statistics for all clades at a certain depth (i.e. if the tree were cut at a depth of *d*, the clades considered here are all the internal nodes at depth *d* from the root). By definition, this value ranges from 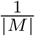 to 1.0.

As above, to compare across different trees solved with various methods, we report this mean membership statistic as a function of the number of clades.

### Triplets Correct Statistic

To compare the similarity of simulated trees to reconstructed trees, we take an approach which compares the sub-trees formed between triplets of the terminal states across the two trees. To do this, we sample ~ 10, 000 triplets from our simulated tree and compare the relative orderings of each triplet to the reconstructed tree. We say a triplet is “correct” if the orderings of the three terminal states are conserved across both trees. This approach is different from other tree comparison statistics, such as Robinson-Foulds [43], which measures the number of edges that are similar between two trees.

To mitigate the effect of disproportionately sampling triplets relatively close to the root of the tree, we calculate the percentage of triplets correct across each depth within the tree independently (depth measured by the distance from the root to the Latest Common Ancestor (LCA) of the triplet). We then take the **average** of the percentage triplets correct across all depths. To further reduce the bias towards the few triplets that are sampled at levels of the tree with very few cells (i.e. few possible triplets), we modify this statistic to only take into account depths where there at least 20 cells. We report these statistics without this depth threshold in Figure S8.

### Application of Camin-Sokal

We applied Camin-Sokal using the “mix” program in PHYLIP [13] as done for reconstructions for McKenna et al [38] and Raj et al [42]. To use “mix” we first factorized the characters into binary ones (thus ending up with Σ_*i*_ *s*_*i*_ binary characters total, where *s*_*i*_ is the number of states that character *i* presented). Then, we one-hot encoded the states into this binary representation where every position in the binary string represented a unique state at that character. We thus encoded every cell as having a 1 in the position of each binary factorization corresponding to the state observed at that character. If the cell was missing a value for character *i*, the binary factorization of the character was a series of ‘?’ values (which represent missing values in PHYLIP “mix”) of length *s*_*i*_. Before performing tree inference, we weighted every character based on the frequency of non-zero (and non-missing values) observed in the character matrix. After PHYLIP “mix” found a series of candidate trees, we applied PHYLIP “consense” to calculate a consensus tree to then use downstream.

### Application of Neighbor-Joining

We used Biopython’s Neighbor-Joining procedure to perform all neighbor joining in this manuscript. We begun similarly to the Camin-Sokal workflow, first factorizing all of the characters into a binary representation. Then, we applied the Neighbor-Joining procedure using the “identity” option as our similarity map.

### Application of Cassiopeia

#### Reconstruction of simulated data

We used Cassiopeia-ILP with a maximum neighborhood size of 10, 000 and time to converge of 12, 600s. Cassiopeia-Hybrid used a greedy cutoff of 200, a maximum neighborhood size of 6000 and 5000s to converge. Cassiopeia-Greedy required no additional hyperparameters. Simulations with priors applied the exact prior probabilities used to generate the simulated trees.

#### Reconstruction of *in vitro* clones

For both Cassiopeia-Hybrid with and without priors, we used a cutoff of 200 cells and each instance of Cassiopeia-ILP was allowed 12, 600s to converge on a maximum neighborhood size of 10, 000. Cassiopeia-ILP was applied with a maximum neighborhood size of 10, 000 and a time to converge of 12, 600s.

#### Simulation of Target Site Sequences for Alignment Benchmarking

To determine an optimal alignment strategy and parameters for our target site sequence processing pipeline, we simulated sequences and performed a grid search using Emboss’s Water algorithm (a local alignment strategy). We simulated 5, 000 sequences. For each sequence, we begun with the reference sequence and subjected it to multiple rounds of mutagenesis determined by a Poisson distribution with *λ* = 3, and a maximum of 5 cuts. During each “cutting” event, we determined the outcomes as follows:

1. Determine the number of Cas9 proteins localizing to the target site in this iteration, where *n*_*cas*9_ ~ *min*(3, *Pois*(*λ* = 0.4)).
2. Determine the site(s) to be cut by choosing available sites randomly, where the probability of being chosen is 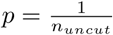 and *n*_*uncut*_ is the number of sites uncut on that sequence.
3. If *n*_*cas*9_ = 1, we determined the type of the indel by drawing from a Bernoulli distribution with a probability of success of 0.75 (in our case, a “success” meant a deletion and a “failure” meant an insertion). We then determined by drawing from a Negative Binomial Distribution as so: *s* ~ *min*(30, *max*(1, *NB*(0.5, 0.1))). In the case of an insertion, we added random nucleotides of size *s* to the cut site, else we removed *s* nucleotides.
4. In the case of *n*_*cas*9_ *≥* 2, we performed a resection event where all nucleotides between the two cut sites selected were removed.
5. After a cut event, we appended the result of the Cas9 interaction to a corresponding CIGAR string

Our Water simulations were exactly 300bp, possibly extending past the Poly-A signal, as would be the case reading off a Nova-seq sequencer.

Upon simulating our ground truth dataset, we performed our grid search by constructing alignments with Water with a combination of gap open and gap extension penalties. We varied the gap open penalties between 5 and 50 and gap extension penalties between 0.02 and 2.02.

To score resulting alignments, we compared the resulting CIGAR string to our ground truth CIGAR string for each simulated sequence. To do so, we first split each cigar string into “chunks”, corresponding to the individual deletions or insertions called. For example, for some CIGAR string 40*M* 2*I*3*D*10*M*, the chunks would be 40*M*, 2*I*, 3*D*, and 10*M*. Then, beginning with a max score of 1, we first deducted the difference between the number of chunks in the ground truth and the alignment. Then, in the case where the number of chunks were equal between ground truth and alignment, we deducted the percent nucleotides that differed between CIGARs. For example, if the ground truth was 100*M* and the alignment gave 95*M*, the penalty would be 0.05.

To find the optimal set of parameters, we selected a parameter pair that not only scored very well, but also located in the parameter space where small perturbations in gap open and gap extension had little effect.

#### Simulation of Lineages for Algorithm Benchmarking

We simulated lineages using the following parameters:

1. The number of characters to consider, *C*
2. The number of states per character, *S*
3. The dropout per characters, *d*_*c*_ ∀*c* ∈ *C*
4. The depth of the tree (i.e. the number of binary cell division), *D*
5. The probability that a site can be mutated, *p*. This is a general probability of cutting
6. The rate at which to subsample the data at the end of the experiment, *M*

To simulate the tree, we begin by first generating the probability of each character mutating to a state, here represented as *p*_*c*_(0, *s*), ∀*s* ∈ *S*. In order to do this, we fit a spline function to the inferred prior probabilities from the lineage tracing experiment. (refer to the section entitled “Determining Prior Probabilities of Indel Formation” for information on how we infer prior probabilities). We then draw *S* values from this interpolated distribution. We then normalize these mutation rates to sum to *p*, therefore allowing in general a *p* probability of mutating a character and 1 − *p* probability of remaining uncut. In the case of the “State Distribution” simulations (Figure S11), we say that *p*_*c*_ is distributed as:

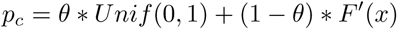

where *F*′(*x*) is the interpolated empirical distribution and *θ* is the mixture component.

Then, we simulate *D* cell divisions, where each cell division consists of allowing a mutation to take place at each character with probability *p*. In the case a mutation takes place, we choose a state to mutate to according to their respective probabilities. Importantly, once a character has been mutated in a cell, that character cannot mutate again.

At the end of the experiment, we sample *M* percent of the cells resulting in 2^*D*^ * *M* cells in the final lineage.

We find that this method for simulating lineages (in particular the method for generating a set of priors on how likely a given state is to form) is able to closely recapitulate observed lineages (Figure S6).

##### Metrics for Comparing Simulations to Empirical Data

We used three metrics of complexity to compare simulated clones to real clones:

- *Minimum Compatibility Distance*: For every pair of character, we define the Minimum Compatibility Distance as the minimum number of cells to be removed to obtain compatibility (Def. 1).
- *Number of Observable States per Cell* : The number of non-zero or non-missing values for each cell, across all characters (i.e. the amount of data that can be used for a reconstruction, per cell).
- *Number of Observable States per Character* : The number of non-zero or non-missing values across for each character, across all cells.

#### Parallel Evolution Simulations for Greedy Benchmarking

As shown above, our greedy approach should accurately reconstruct a lineage if a perfect phylogeny exists. In order to better quantify how much our greedy algorithm’s performance is affected by parallel mutations, we decided to simulate “near perfect phylogenies”, whereby we first began by simulating a perfect phylogeny, and afterwards introduced double mutated characters.

Specifically, we begin by simulating perfect phylogenies with 40 − *k* characters. We then fix a depth, *d*, and sample a node from said depth. We choose two grandchildren randomly from this node (one from each child) and introduce the same mutation on each of the edges from each child to grandchild, thereby violating the perfect phylogeny. We repeat this process *k* times. This thus creates an analysis, as presented in Figure S4, whereby accuracy can be evaluated as a function of both depth of parallel evolution, *d*, and the number of events that occurred, *k*.

#### Simulation of “Base Editor” Technologies

We used the simulation framework described above to simulate base-editor technologies. To explore the trade off between the number of states and the number of characters, we simulated trees with 40, 50, 80, and 100 characters while maintaining the product of characters and states equal at 400 (thus we had trees of 10, 8, 5, and 4 states per character, respectively). The dropout per character was set to 10%, the mutation rate per character was set to 1.04% (a previously observed mutation rate [26]), and a depth of 10 where 400 cells where sampled. For each character/state regime, we generated 10 trees for assessing the consistency of results. We use a negative binomial model (~ *NB*(5, 0.5)) as the editing outcome distribution (i.e. state distribution).

#### Simulation of “Phased Recorder” Technologies

To simulate the phased recorder, we used 5 different experiments varying mutation rates across 50 characters and 10 states per character. In each experiment, we chose a mutation rate for each character from one of 10 regimes, each differing in their relationship to the base mutation rate *p*_0_. To systematically implement this, mutation rate for *χ*_*i*_ is described as such:

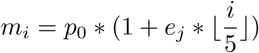

where *p*_0_ = 0.025 and *e*_*j*_ is a experiment scalar in **e** = {0, 0.05, 0.1, 0.25, 0.5}. This means that for characters 1 − 5, *m*_*i*_ = *p*_0_, for characters 6 − 10, *m*_*i*_ = *e*_*j*_*p*_0_, for characters 11 − 15, *m*_*i*_ = 2*e*_*j*_*p*_0_, etc. To summarize each experiment, we provide the ratio between the maximum and minimum mutation rates, which is by definition 1 + 10*r*_*j*_ (because we had 50 characters). We compare two models of indel formation rates - the first being a negative binomial model (~ *NB*(5, 0.5)), and the second being the spline distribution fit from empirical data.

We simulated 10 trees per regime and reconstructed trees with Cassiopeia with and without priors.

### Reconstructions of GESTALT Datasets

We downloaded data corresponding to the original GESTALT study [38] and the more recent scGESTALT study from https://datadryad.org/resource/doi:10.5061/dryad.478t9 and GSE105010, respectively. We created character matrices for input into Cassiopeia by creating pivot tables relating each cell the observed indel observed at each one of the 10 tandem sites on the GESTALT recorder. We then reconstructed trees from these character matrices using one of five algorithms: Camin-Sokal (used in the original studies), Neighbor-Joining, Cassiopeia’s greedy method, Cassiopeia’s Steiner Tree method, and Cassiopeia’s hybrid method.

For each reconstruction, we record the parsimony of the tree, corresponding to the number of mutations that are inferred along the reconstructed tree. We display these findings in Figure 6a, where we have Z-normalized the parsimonies across the methods for each dataset to enable easier visualization of relative performances.

### Visualization of Trees

To visualize trees we use the phytools R package. Colors in the heatmap denote a unique mutation, gray denotes an uncut site, and white denotes dropout.

---

**Figure S1:**
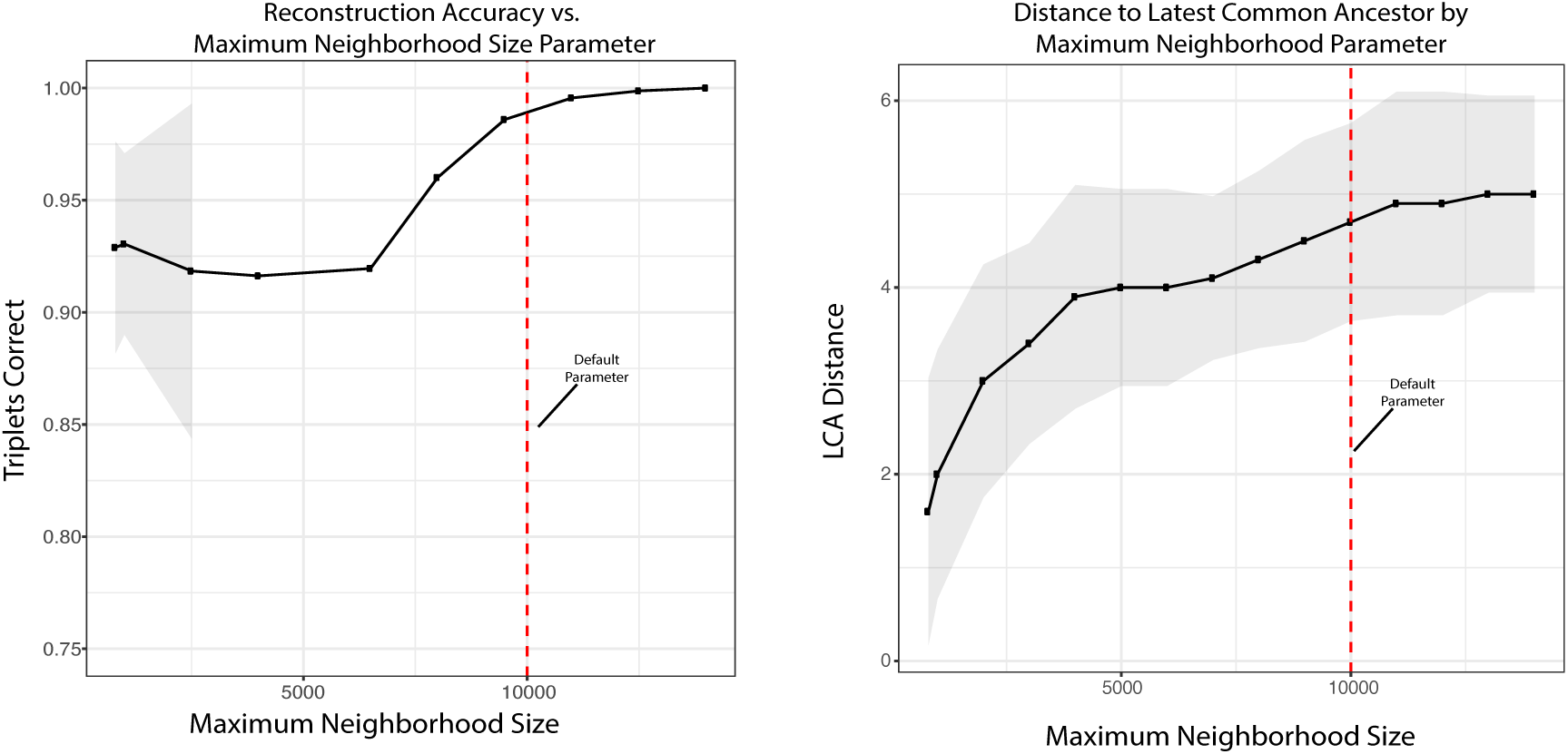
Stability analysis of maximum neighborhood size parameter for Steiner Tree approach. The maximum neighborhood size is a central parameter provided by the user when inferring the potential graph necessary as input to the Steiner-Tree solver (see methods). Here, we bench-mark the stability of solutions with respect to several maximum neighborhood sizes using 10 trees with default parameters (40 characters, 40 states, 2.5% per-character mutation rate, depth of 11, and an average dropout rate of 17% per character). We quantify both the reconstruction accuracy with respect to the reconstructions found with the largest maximum neighborhood size (14, 000 nodes) which displays a saturation at around 9, 000 nodes. To provide intuition for the accuracy of the potential graph (represented as the maximum distance to the ‘latest common ancestor’ (LCA) which is dynamically solved for, given a maximum neighborhood size) we display the LCA allowed for each maximum neighborhood size parameter. In both figures, we display lines connecting the mean values; shaded regions are the standard deviation of the measurements across the 10 replicates.

**Figure S2:**
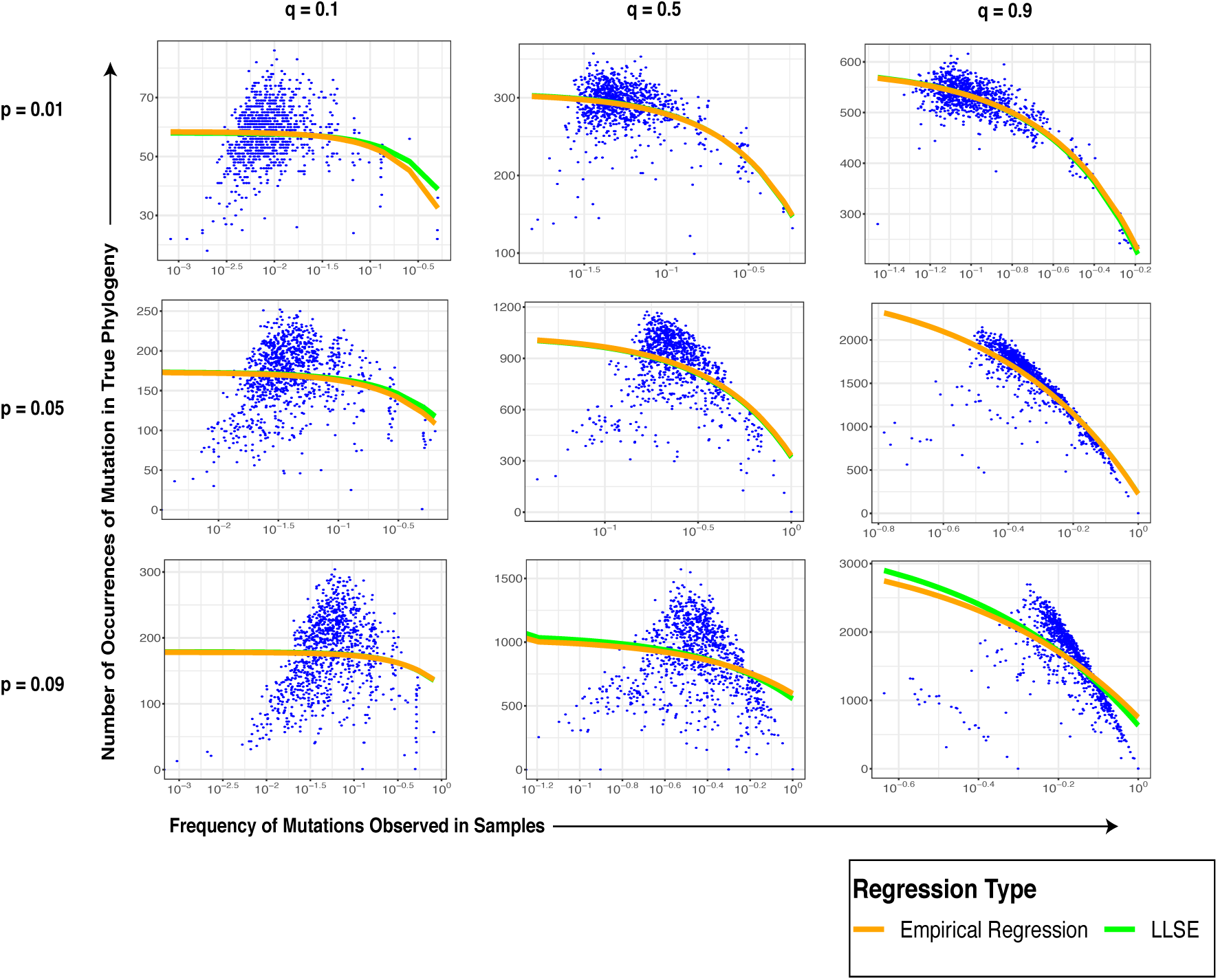
Observed Frequency of Mutations is Measure of True Mutation Count. The true number of occurrences of a mutation is estimated well by the observed frequency at leaves. We use a Linear Least Squares Estimate to quantify the relationship between the expected number of times a mutation occurred given the observed frequency at the leaves (Eq. 3). Using various rates for character and indel mutation rates (*p* and *q*, respectively) we show that this relationship is negative (i.e. greater observed frequencies tend to correspond to mutations that occurred few times near the top of the phylogeny) for a range of biologically-relevant values.

**Figure S3:**
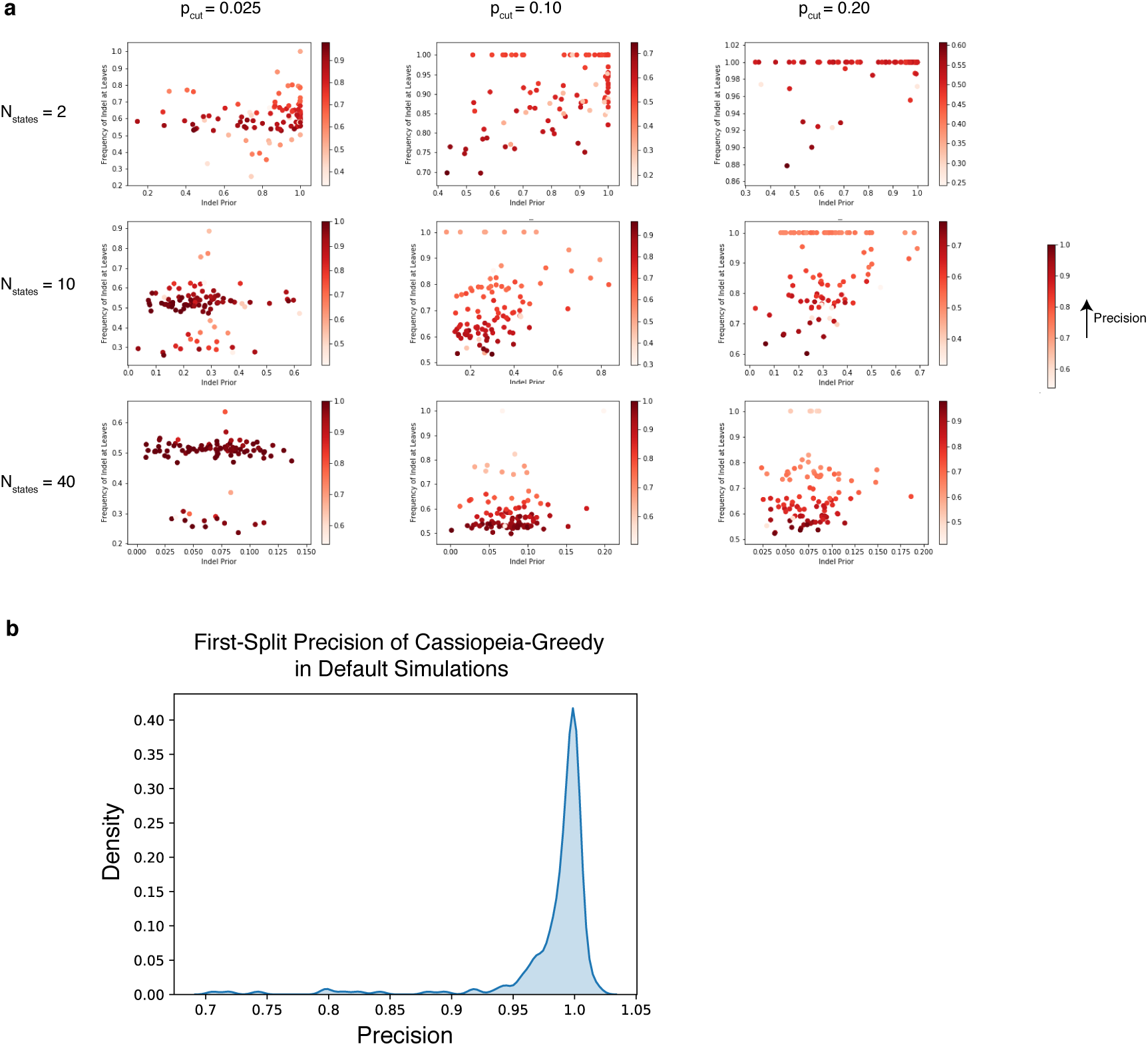
Precision of Cassiopeia-Greedy First Split. (a) The precision of greedy splits of 400 cells was measured with varying mutation rates and states per character, wihtout dropout. For each pair of parameters (number of states and mutation rate), we measure precision as a function of the conditional probability of the selected (character, state) pair and the frequency of that mutation observed in the 400 cells. (The conditional probability for state *j, q*(*j*) is defined as *Pr*(*χ* → *j*|*χ mutates*)). Precision was defined as the proportion of true positives in the greedy split (see Methods). Each point indicates a replicate (100 per plot) and the heat represents the precision. (b) The density histogram (smoothed using a kernel density estimation procedure) of all first-split precision statistics from Cassiopeia-Greedy on default simulations (i.e. 40 characters, 40 states, 2.5% mutation rate, 11 generations, 400 cells, and 18% dropout rate). We measured a median precision of 0.99 across all default simulations.

**Figure S4:**
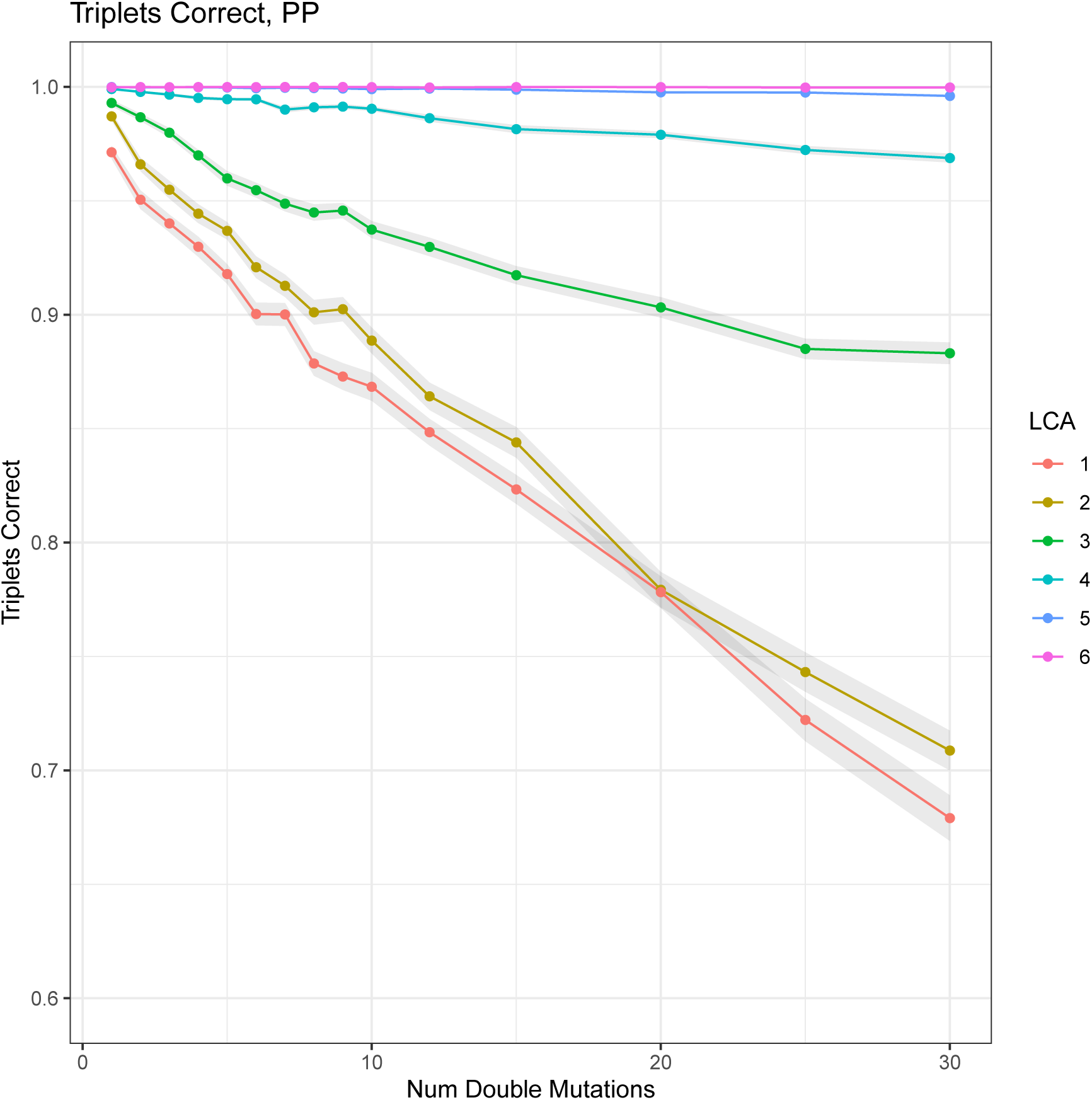
Benchmarking of parallel evolution on the greedy heuristic. The greedy heuristic, inspired by algorithms to solve the case of perfect phylogeny (see methods), is impacted by two factors: (1) the number of parallel evolution events (i.e. the same mutation occurs more than once in the experiment) and (2) the depth from the root these mutations occur at. Here, each line represents a series of experiments increasing the number of ‘double mutations’ (i.e. the simplest case of parallel evolution where a mutation occurs exactly twice) where the ‘latest common ancestor’ (LCA) is a set depth from the root.

**Figure S5:**
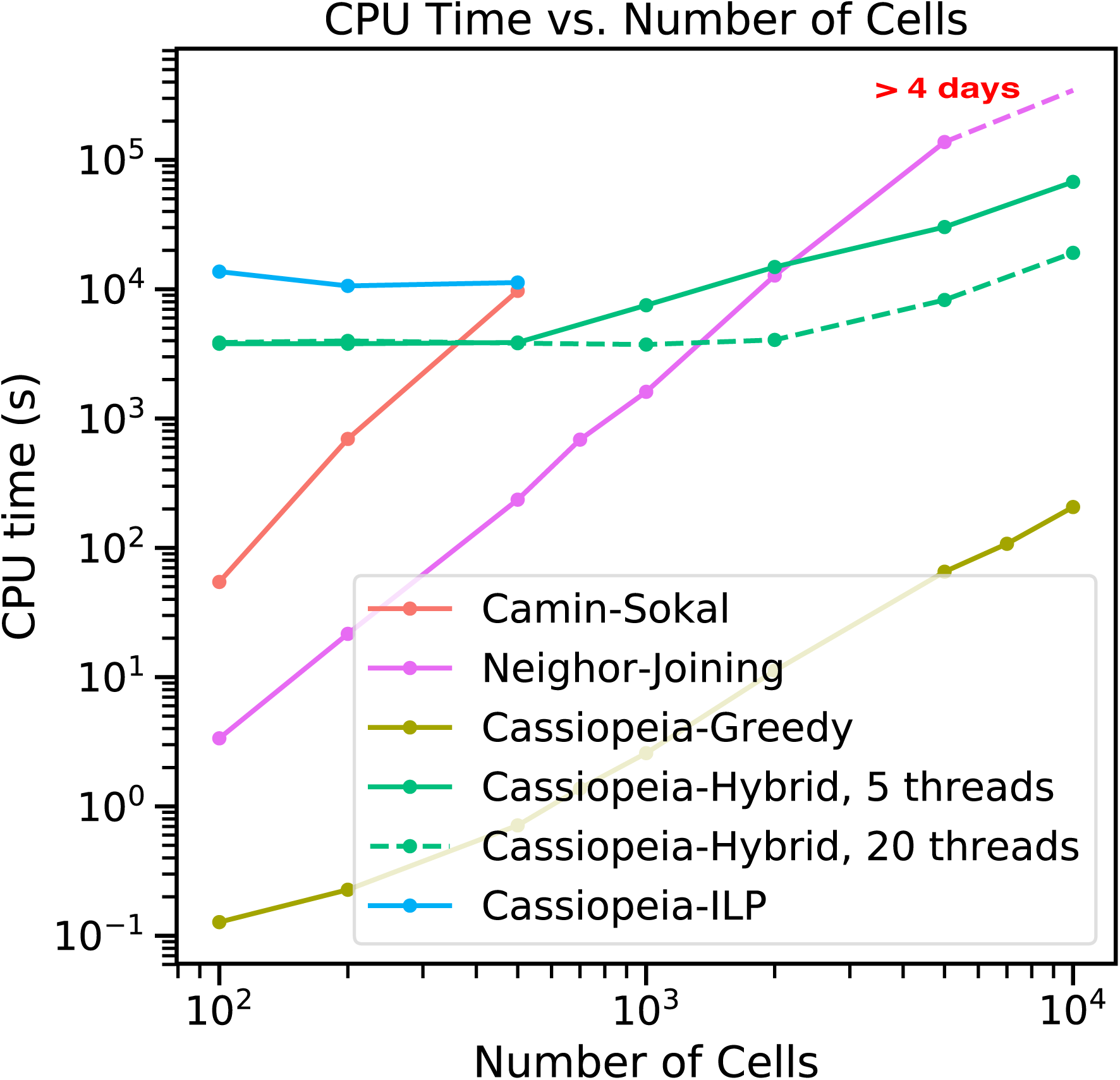
Time complexity of lineage reconstruction approaches. Time complexity, as measured in seconds, of each algorithm tested in this manuscript is compared using simulated datasets ranging from 100 cells to 10,000 cells. Default settings for the simulations were used (0.025 mutation rate, 40 characters, 10 states, and 0.18 median dropout rate). Cassiopeia was tested using default parameters of a maximum neighborhood size of 3000, time to converge of one hour, and a greedy cutoff of 200 cells. Cassiopeia was tested using 5 threads and 20 threads, illustrating the advantage of parallelizing the reconstruction algorithm. ILP, which was only run until 500 cells due to the infeasibility of running on larger datasets, was allowed 10000s to converge on a maximum neighborhood size of 20,000 (the default settings). Neighbor-Joining could not reconstruct a tree for 10,000 cells within 4 days when the reconstruction was terminated.

**Figure S6:**
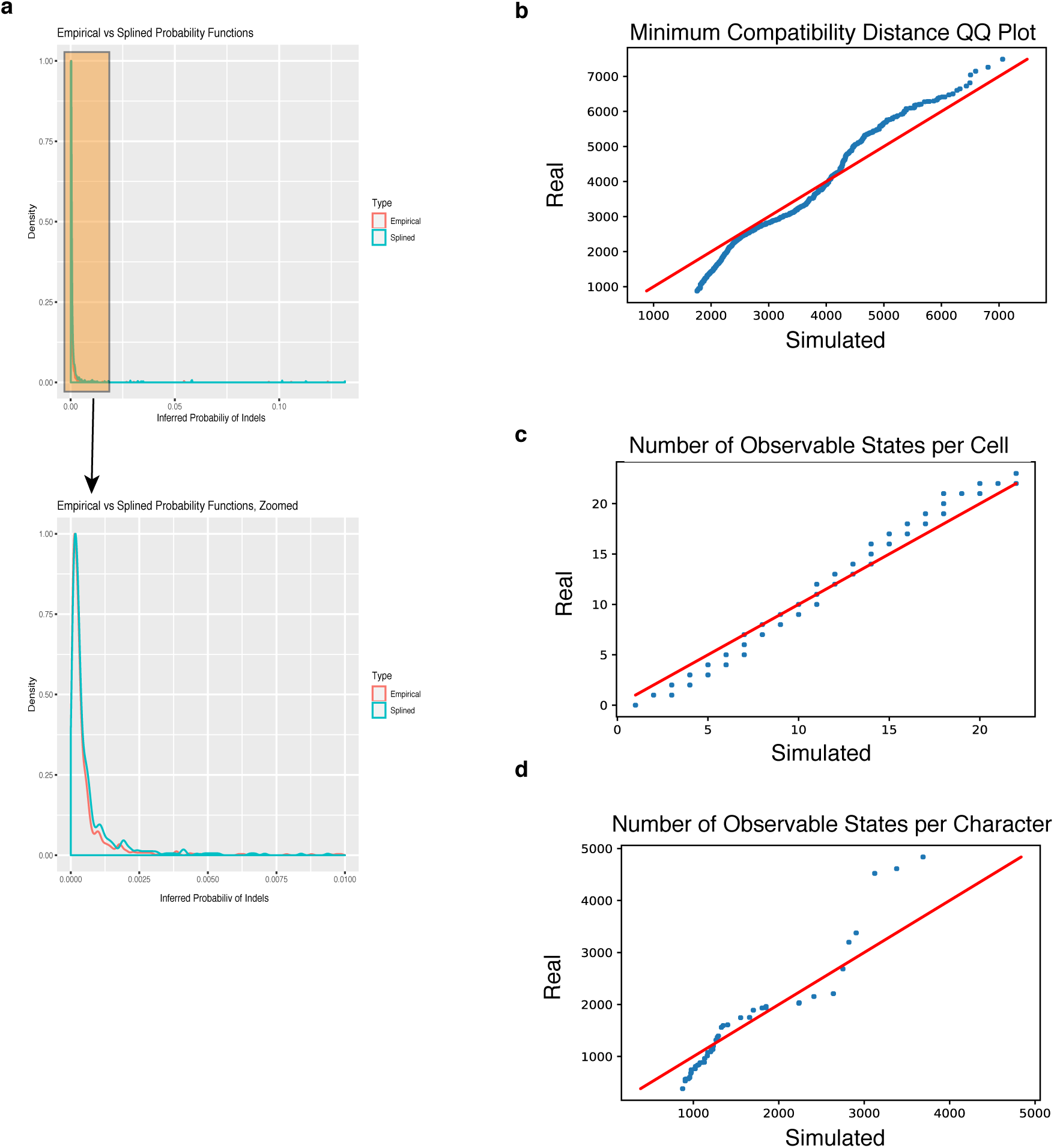
Determination of mutation rates used in simulation. We use an interpolation of the empirical indel distribution as input for the conditional probability of a state arising given a mutation. (a) A comparison of the empirical and ‘splined’ indel distributions; a zoomed in version is provided for comparison at low probabilities. (b-c) A comparison of three metrics between an observed clone (clone 3) and a simulated clone using inferred parameters. We used the number of character, states, per-character mutation rate, and dropout probabilities inferred from the empirical data; the indel formation rates were calculated using a polynomial spline function. (b) measures the ‘minimum compatibility distance’ for all pair-wise character combinations (see methods). (c) compares the number of observable states per cell. (d) compares the number of observable states per character.

**Figure S7:**
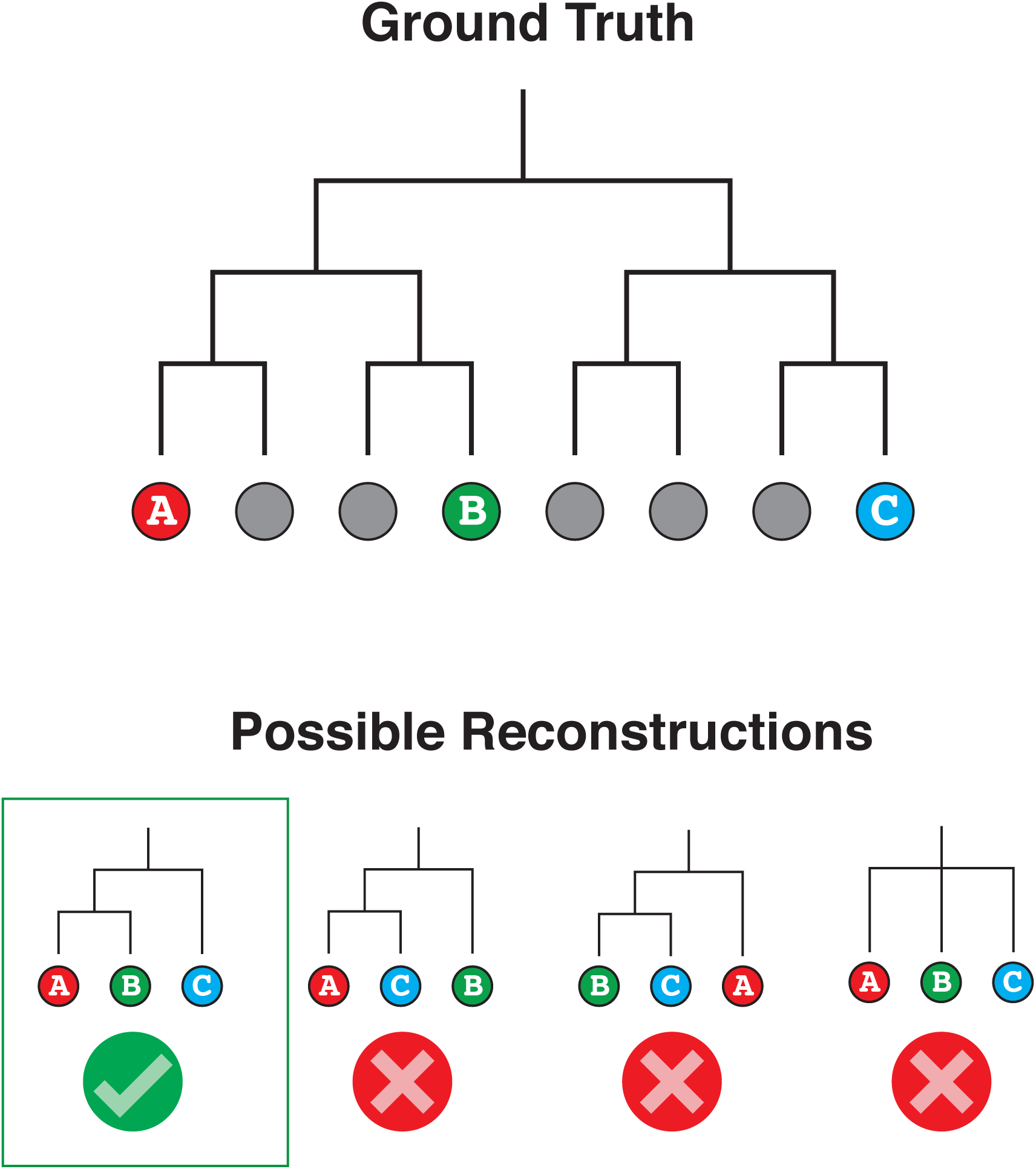
Triplets Correct Statistics. Schematic for the Triplets Correct statistic, the combinatorial metric used to compare between trees. In this metric, we compare the relative orderings of three leaves between two trees (e.g. the “Ground Truth” and a reconstruction). There are four possible ways that a triplet could be ordered here, based on the relationship between each leaf and the Latest Common Ancestor (LCA) of the triplet. The statistic tallies the number of correct triplets and reports this value weighted by the depth of the LCA from the root.

**Figure S8:**
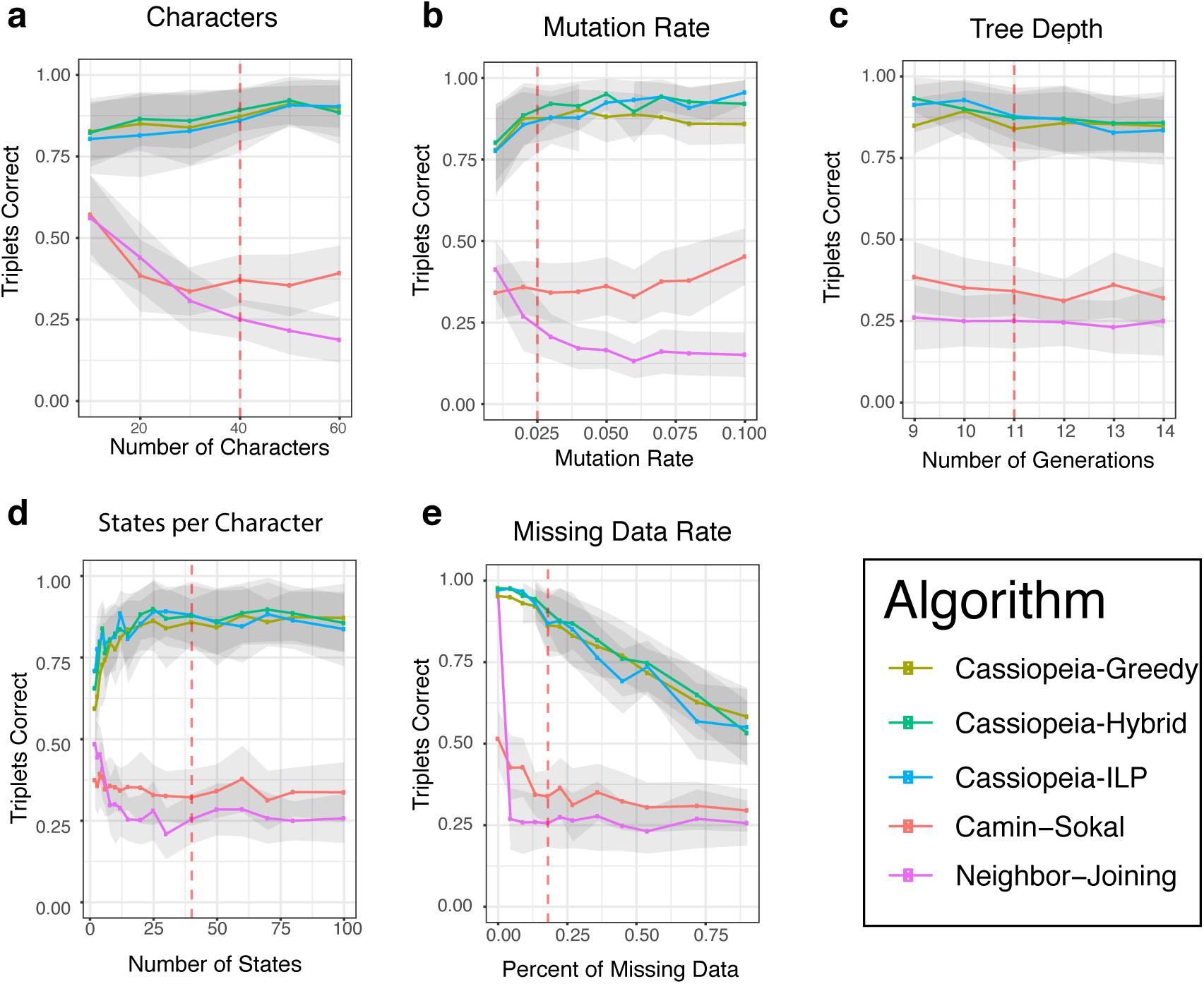
Unthresholded Triplets Correct Statistics. The Triplets Correct statistic reported for synthetic benchmarks presented in Figure 2 without removing triplets whose LCA-depth was sampled deeply enough (by default, a given triplet at depth *D* is only considered if a sufficient number of triplets at depth *D* is observed). Here, the effective threshold is 0.

**Figure S9:**
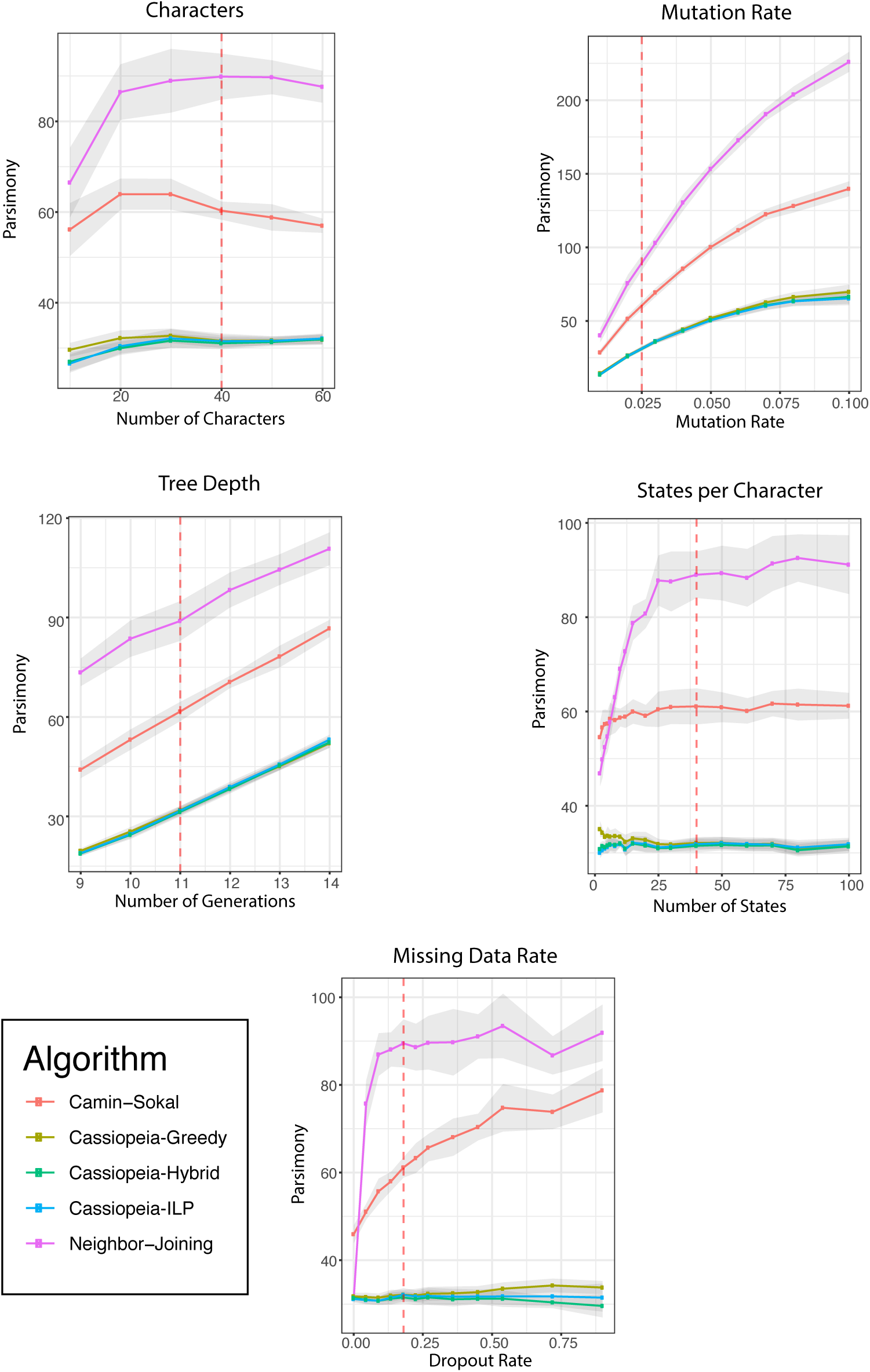
Parsimony of reconstructed trees of 400 cell simulated datasets. Parsimony scores (or number of evolutionary events) for each reconstructed network presented in Figure 2 were calculated and compared across phylogeny reconstruction methods. Results are presented for the number of characters, the mutation rate, tree depth, number of states and dropout rate for all five algorithms used in this study. Standard error is represented by shaded area.

**Figure S10:**
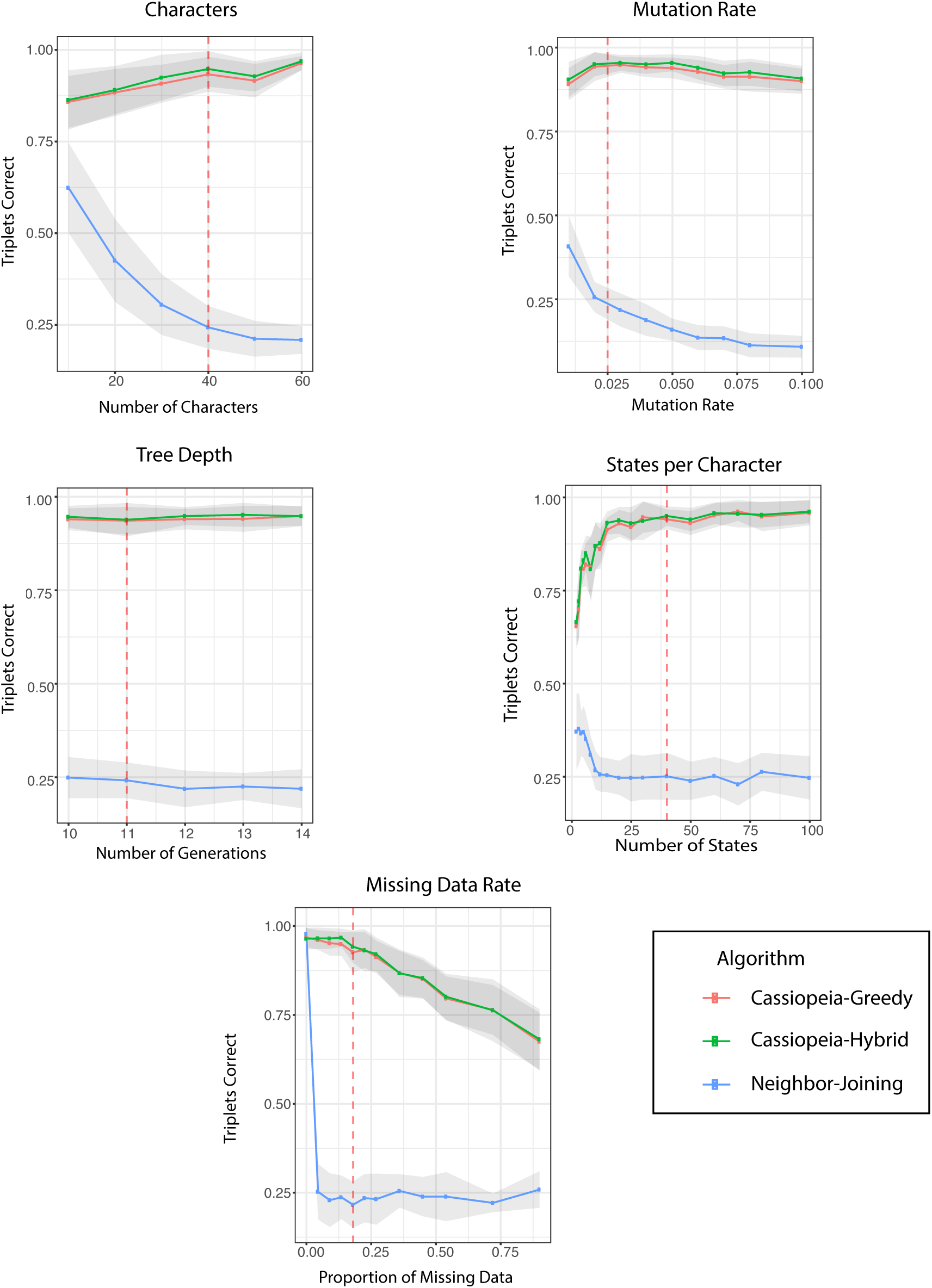
Benchmarking of lineage tracing algorithms on 1000 cell synthetic datasets. Phylogeny reconstruction algorithms were benchmarked on simulated trees consisting of 1,000 cells. The number of characters, character-wise mutation rate, length of experiment or tree depth, number of states, and dropout rate were tested. Due to scalability issues, only greedy, hybrid, and neighbor-joining were tested. Standard error is represented by shaded area.

**Figure S11:**
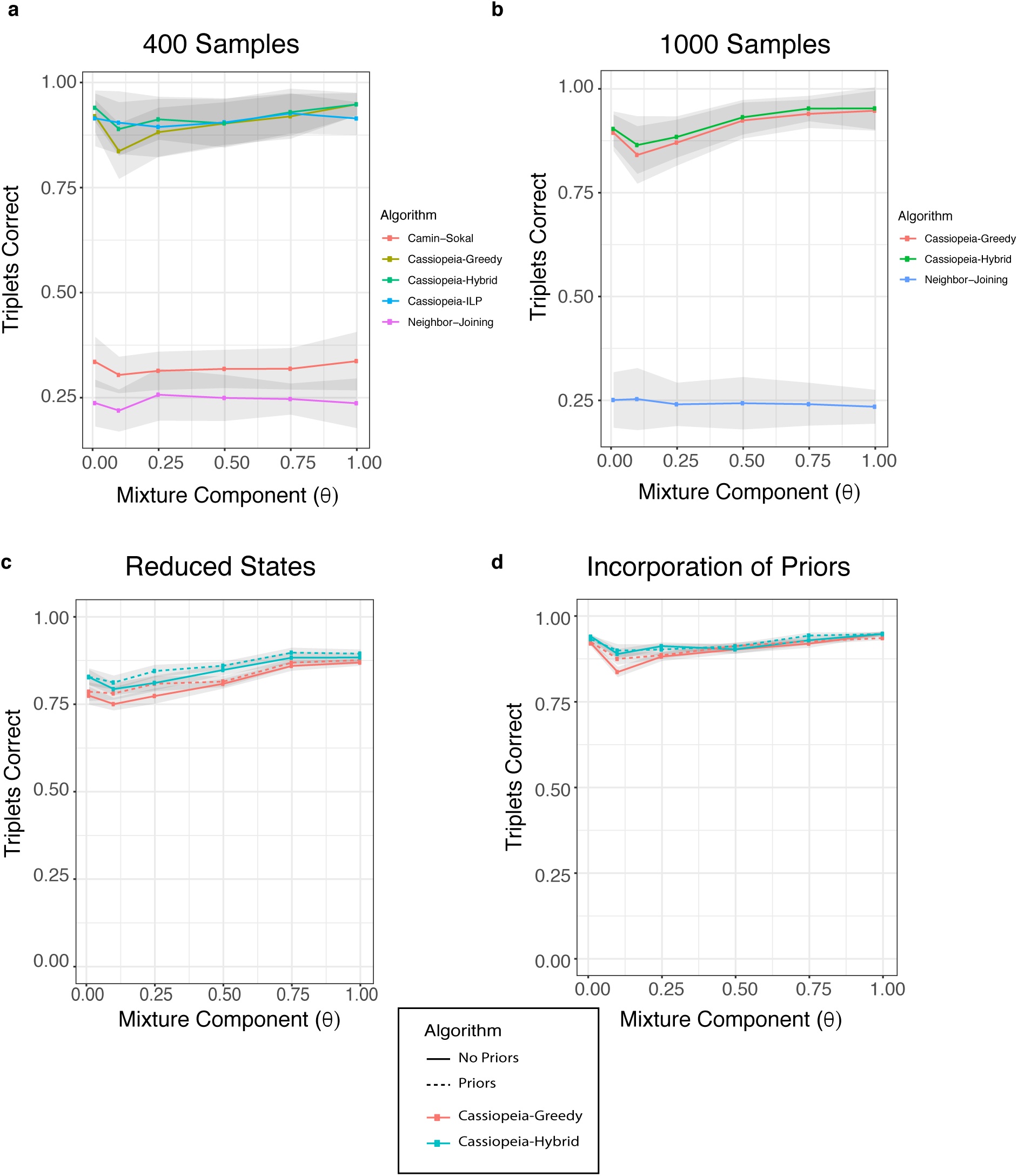
Reconstruction accuracy under over-dispersed state distributions. The effect of the indel distribution (i.e. the relative propensity for a given indel outcome) was explored in various regimes using a mixture model. Here, the mixture model consisted of mixing the inferred indel distribution with a uniform distribution between 0 and 1.0 with some probability *θ* (i.e. when *θ* = 1.0, the indel distribution was uniform). In all simulations, we used default parameters for the simulated trees unless stated otherwise (40 characters, 40 states, depth of 11, median dropout rate of 17%, and a character mutation rate of 2.5%). (a) displays the results of all five algorithms over 400 samples. (b) displays results for simulations over 1000 samples for hybrid, greedy, and neighbor-joining methods. (c) Simulations for 400 samples using 10 states rather than 40 states per character. Dashed lines represent reconstructions performed with priors. (d) Simulations over 400 samples and 40 states, comparing results with and without priors. Dashed lines represent reconstructions performed with priors.

**Figure S12:**
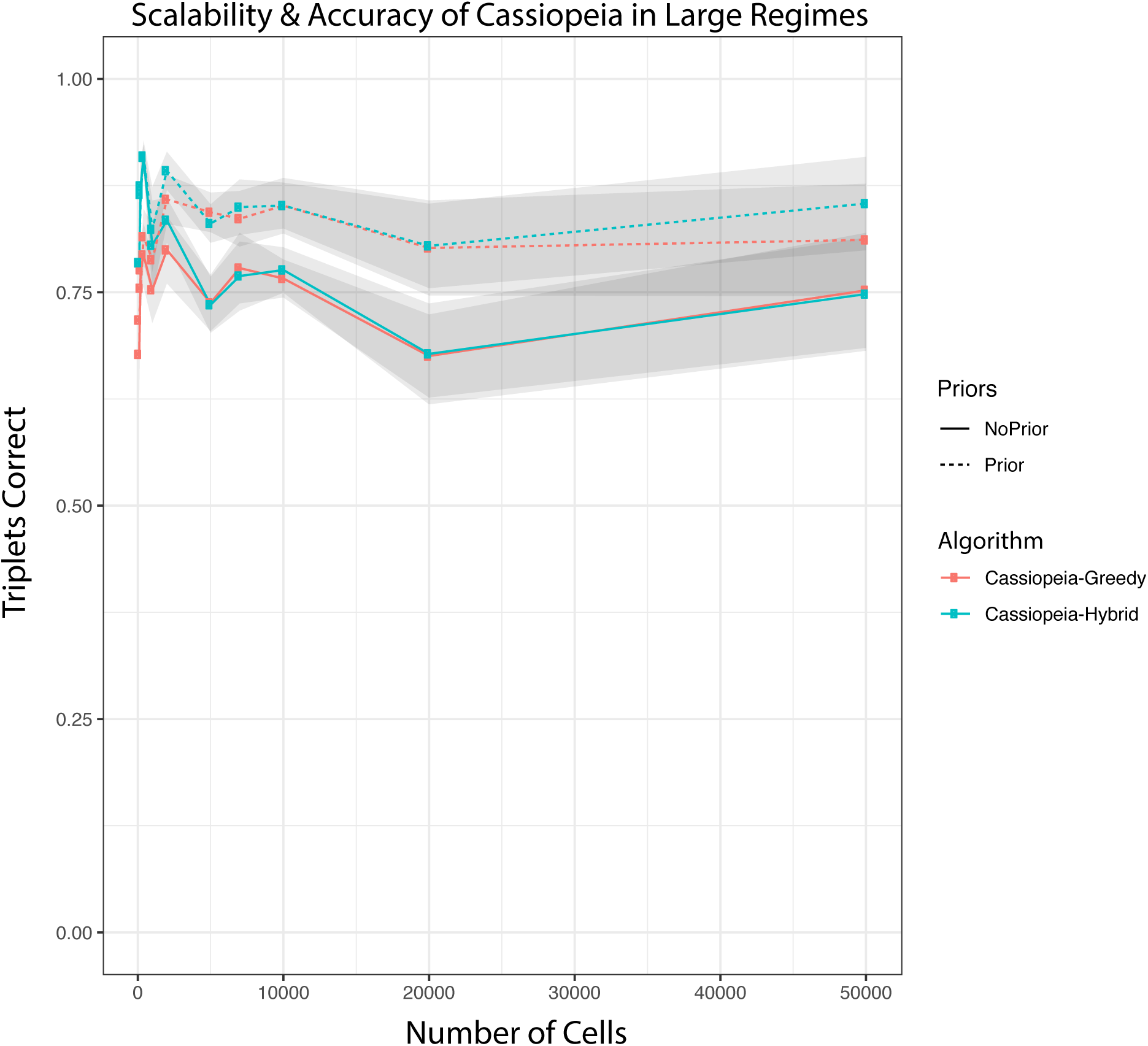
Benchmarking of greedy and hybrid algorithms on large experiments. Triplets correct is used to measure the accuracy of reconstructions using both hybrid and greedy algorithms on large trees (up to 50, 000 cells). Of note, hybrid and greedy have comparable results on larger trees, which remain accurate even in these massive regimes. In addition, the knowledge of prior probabilities of particular states confers a large increase in accuracy.

**Figure S13:**
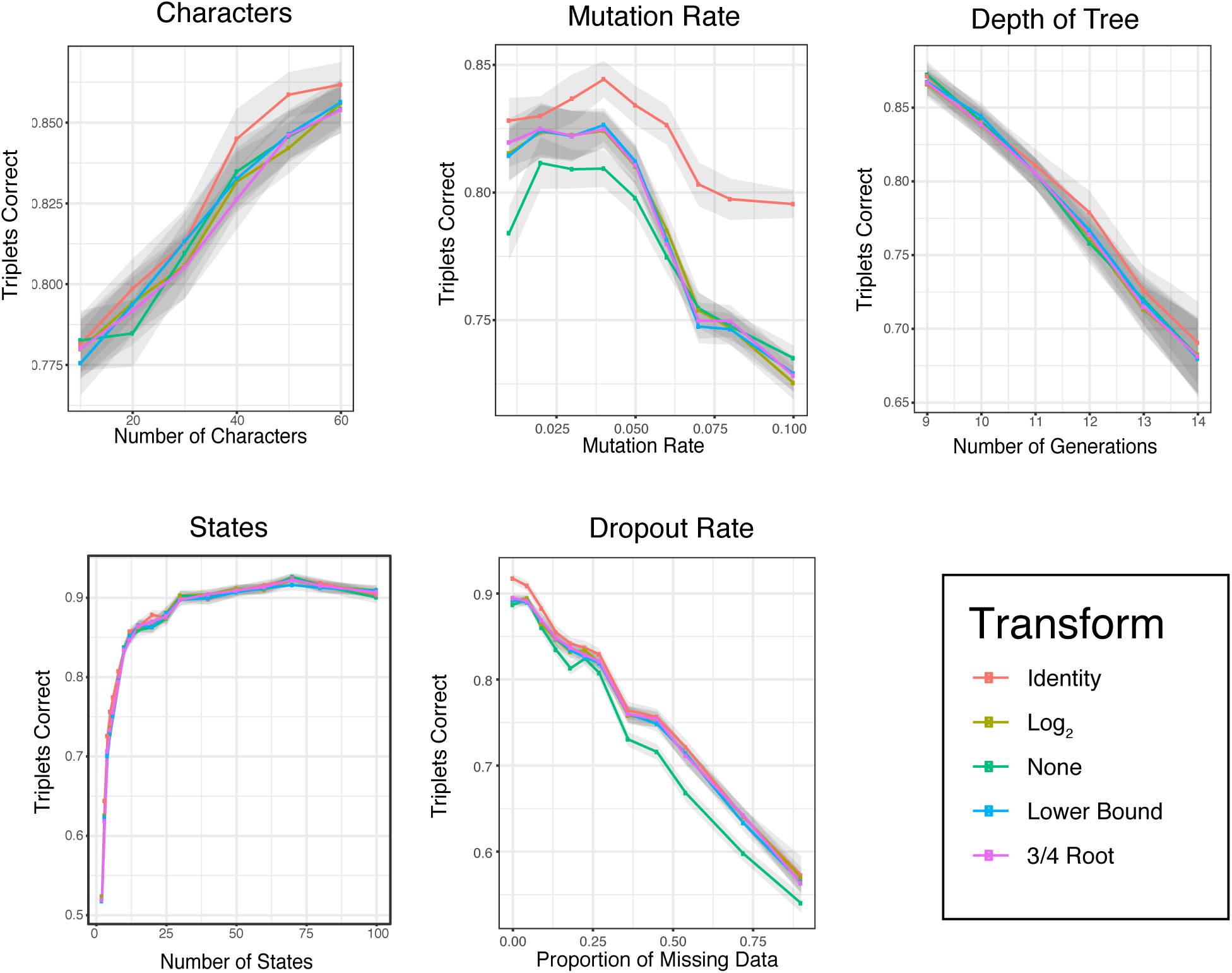
Determination of the indel prior transformation function. The effect of incorporating the prior probabilities of mutation events into the greedy algorithm is explored using synthetic datasets. The exact mutation probabilities used for simulations are used during reconstruction (i.e. the mutations drawn during simulation). Five possible transformations *f* (*n*_*i,j*_), representing an approximation of the future penalty of not choosing this mutation (see methods) were tested for incorporation with the priors. The transformations were: (i) Identity (*f* (*n*_*i,j*_) = *n*_*i,j*_), (ii) Log2 (*f* (*n*_*i,j*_) = *log*_2_(*n*_*i,j*_)), (iii) None (*f* (*n*_*i,j*_) = 1), (iv) Lower Bound 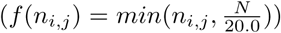, and (v) 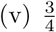 root 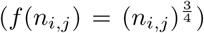. *n*_*i,j*_ denotes the number of cells which report the mutation *j* in character *i* and *N* is the total number of samples. To test these transformations, we evaluated the resulting tree accuracy via Triplets Correct. Standard error is represented by shaded area.

**Figure S14:**
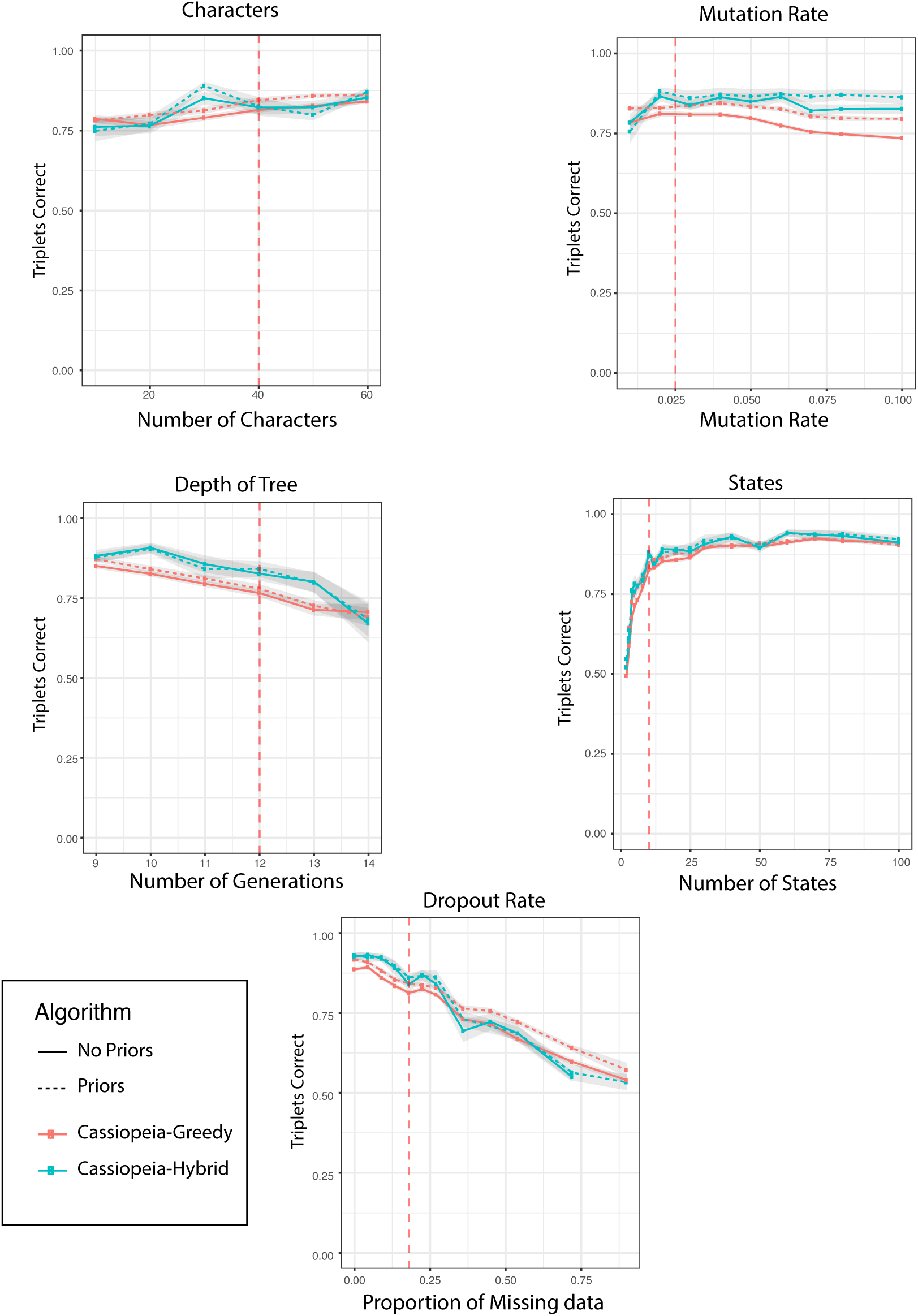
Incorporation of priors into Cassiopeia. A comparison of tree accuracy when using priors for both the greedy-only method and Cassiopeia. We compared performance as we varied the number of characters per cell, the mutation rate per character, the length of the experiment, the number of states per character, and the amount of missing data. Standard error is represented by shaded area.

**Figure S15:**
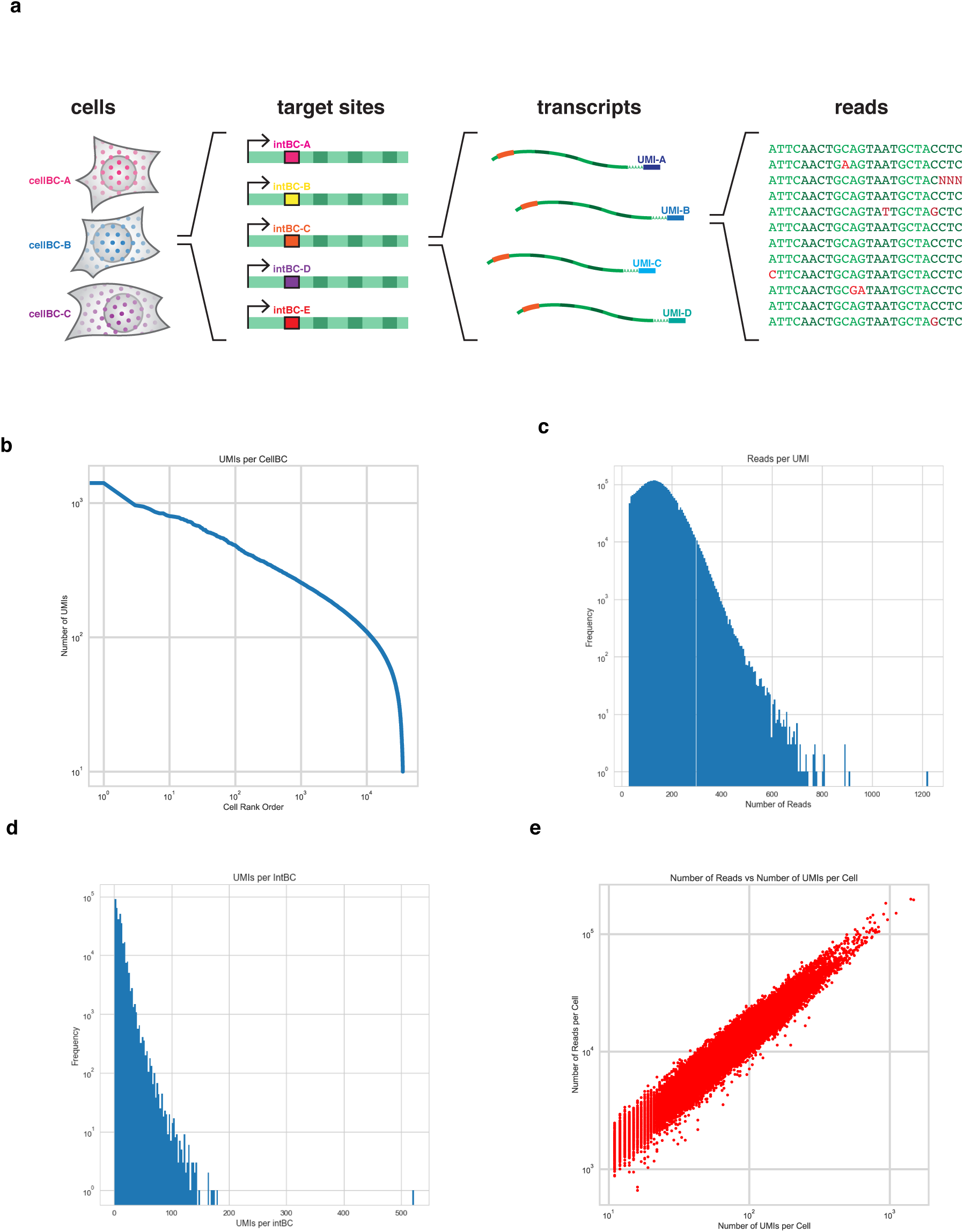
Quality control metrics for the target site sequencing library processing pipeline. (a) schematizes the target site library that is output from the lineage tracing experiment as described in [7]. Cells consist of multiple target sites, each of which contains 3 separate & independently targeted cut sites. Each target site is indexed by an integration barcode (intBC) for phasing of mutations. Each cell contains roughly 5-20 target sites (and on average 9), as determined by the number of unique intBCs observed after sequencing and processing. Target sites are read off of RNA transcripts where many RNA transcripts can correspond to a single target site, and each transcript is read several times. (b-e) present quality control metrics after the processing pipeline. Cells are ranked by the number of UMIs they contain and plotted in (b); (c) contains the number of reads per UMI; (d) contains the number of UMI per intBC; (e) is the concordance between reads per cellBC and UMIs per cellBC.

**Figure S16:**
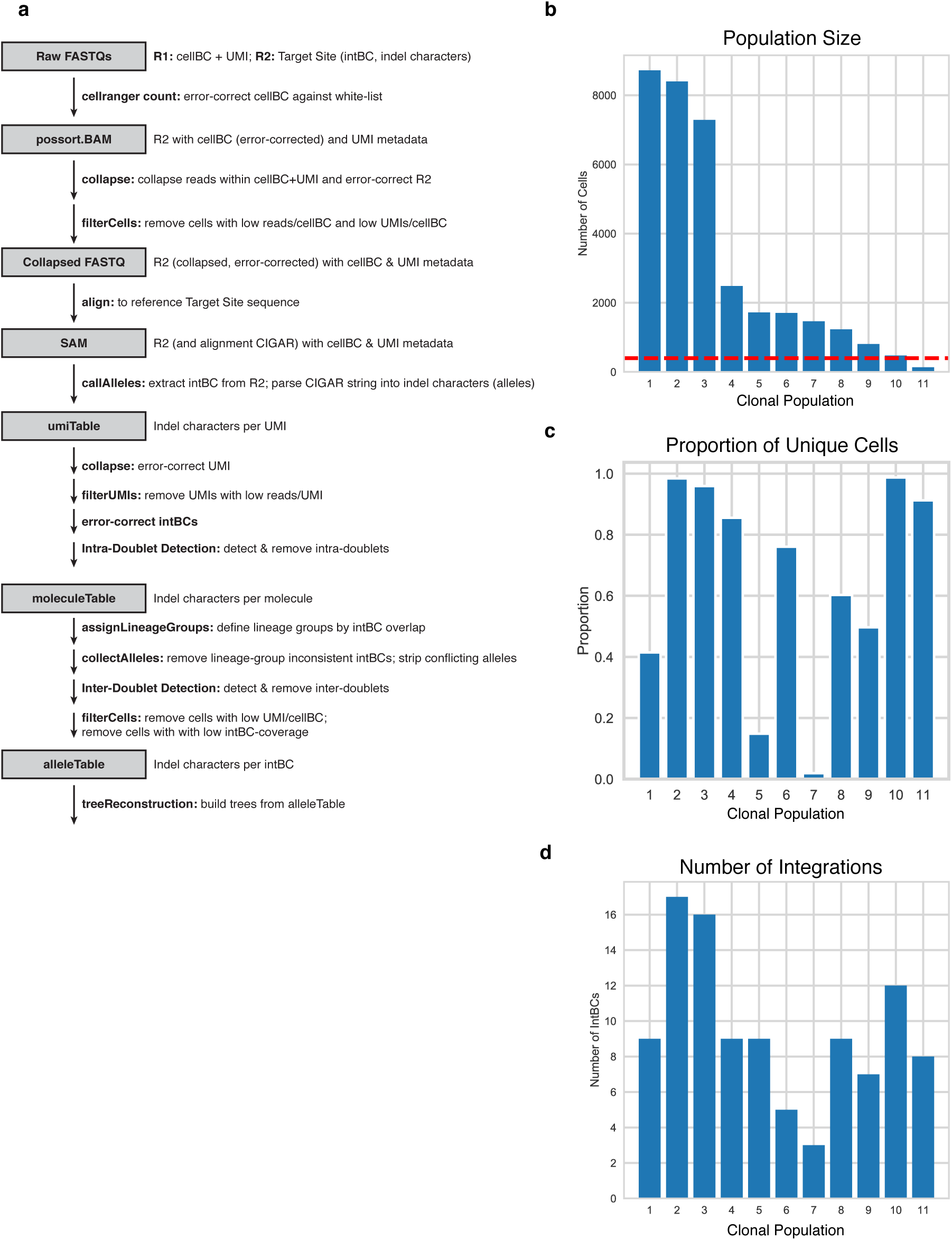
Processing pipeline for *in vitro* dataset. (a) describes a flowchart of the processing pipeline taking as input the raw FASTQs from a sequencing run and converting the observed reads into final trees. Cellranger “count” [58] is used to map reads to dummy transcriptome (junk sequence that nothing will align to), filter cells, and read off the 10x cell barcodes and UMIs. The resulting BAM file is then passed through a series of cell filtering, UMI error correction, and allele mapping before becoming the final allele table that can be converted to character matrices for clone reconstruction. See methods for more detailed information for each step. (b-c) present additional summary statistics for the final allele table. (b) displays the number of cells per clone; (c) shows the median number of intBCs observed in each clone.

**Figure S17:**
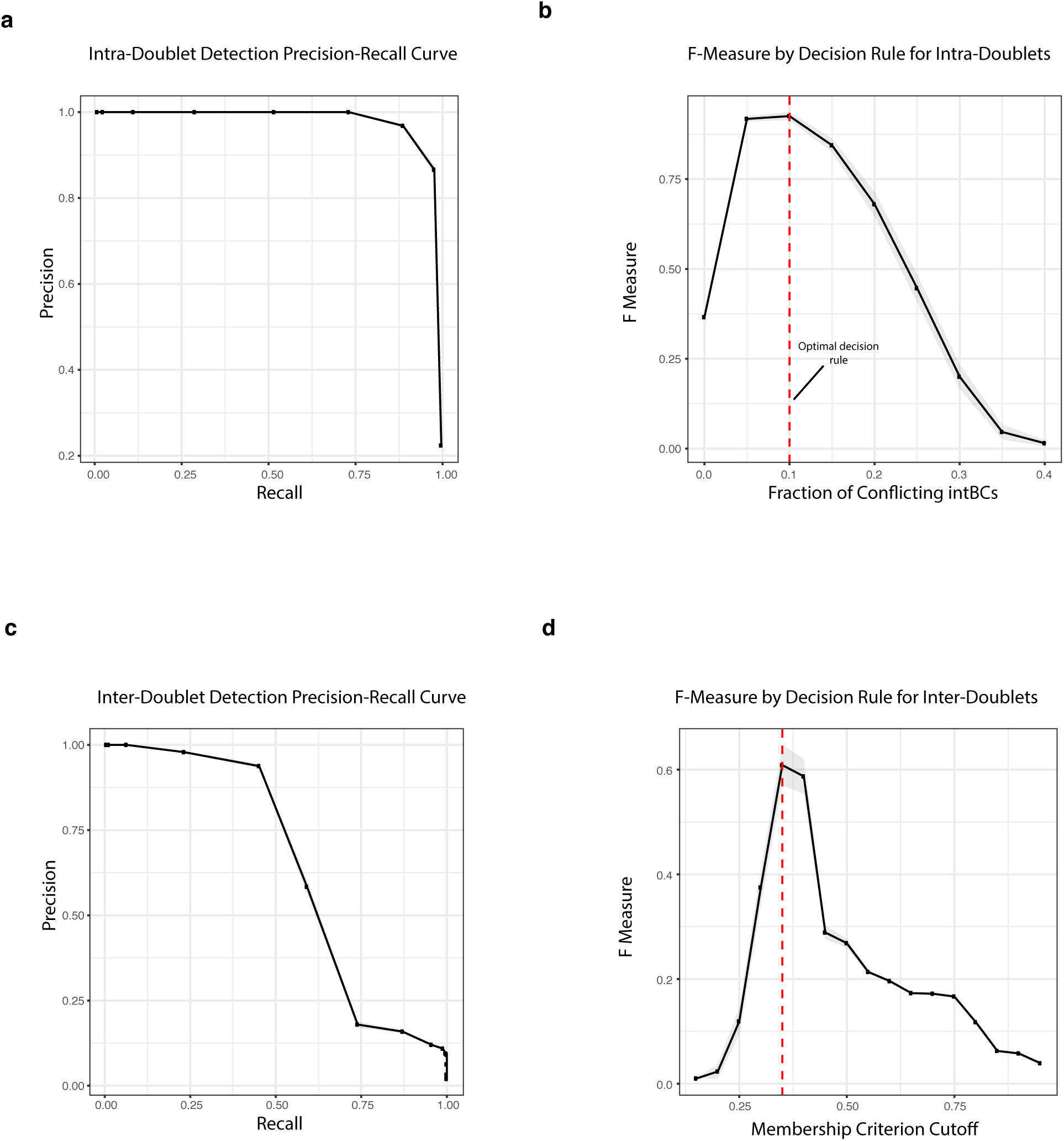
Identification of doublets using intBCs. IntBCs are used to identify doublets. (a-b) report the ability to identify doublets arising from the same clone, referred to as “intra”-doublets; (c-d) report the ability to identify doublets arising from different clones, referred to as “inter”-doublets. Doublets were simulated using the final allele table and 200 “intra”- and “inter”-doublets were created in each of 20 replicates. Precision-recall curves for intra- and inter-doublet detection methods are presented in (a) and (b), respectively. (c) and (d) present the F-measure (defined as the weighted harmonic mean between precision and recall) of detection methods for intra- and inter-doublets, respectively. Red-dashed lines denote the optimal decision rule for doublet detection. Standard error is represented by shaded area.

**Figure S18:**
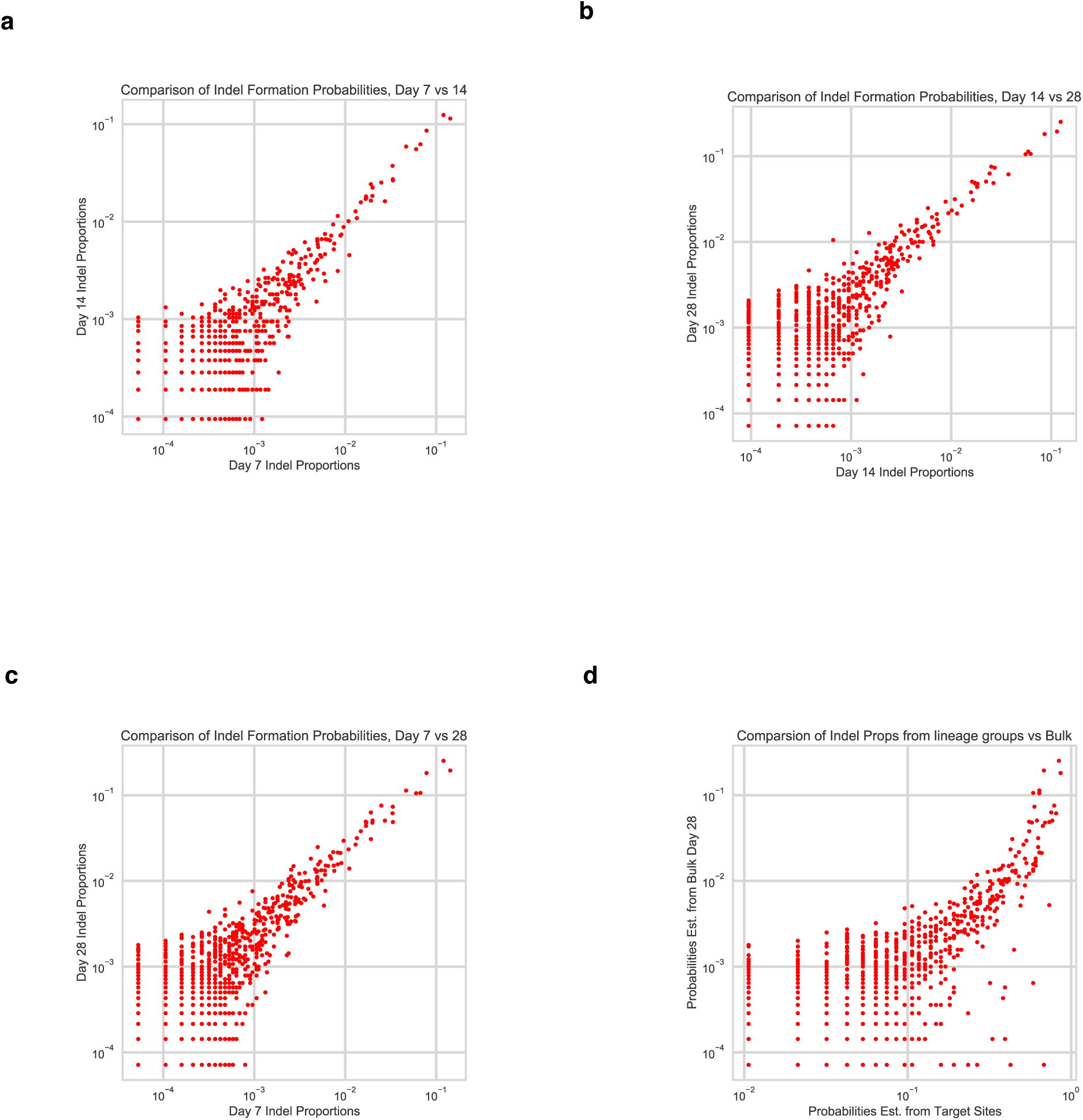
Estimation of Prior Probabilities for Tree Reconstruction. Prior probabilities to be used during tree reconstruction can be determined from both a bulk assay and independent clonal populations. Prior probabilities of mutations were determined by calculating the proportion of unique intBCs that report a particular indel (see methods). The bulk assay consisted of several independent clones with non-overlapping intBCs grown over the course of 28 days. (a-c) report the correlation of indel formation probabilities between various time points in the bulk experiment. A strong correlation is observed between all time points: 7 and 14 (a), 14 and 28 (b) and 7 and 28 (c). Indel formation probabilities can also be calculated using the int-BCs from each clone as independent measurements. Using this method, (d) reports the correlation between this lineage-group specific probability calculation and the last time point of the bulk assay.

**Figure S19:**
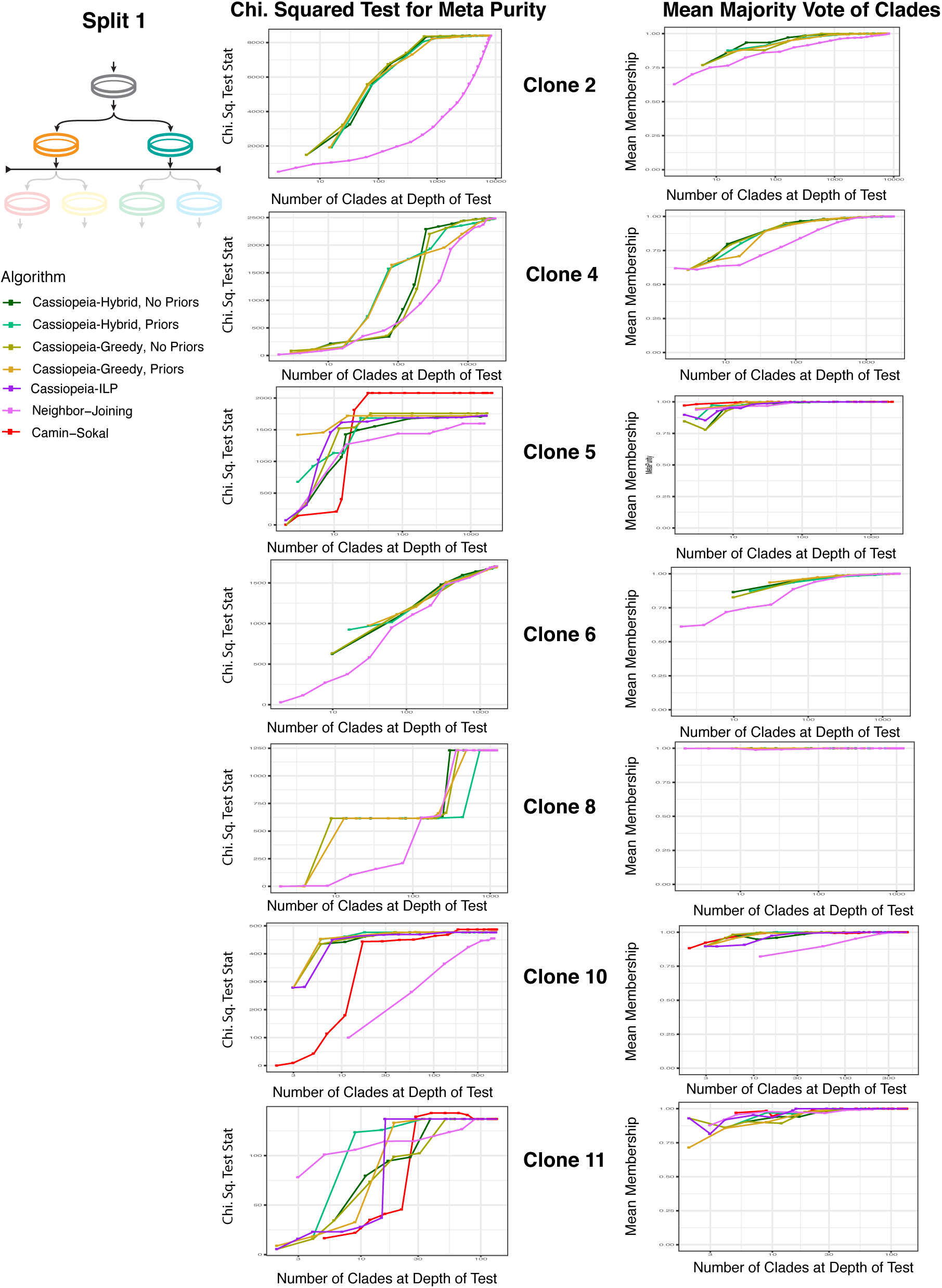
Evaluation of algorithms on *in vitro* lineage tracing clones, First Split. Trees were reconstructed for the remaining clones in the *in vitro* dataset that consisted of more than 500 unique cell states. LG2, LG4, LG6, and LG8 passed this threshold and were reconstructed with Cassiopeia (with and without priors), greedy-only (with and without priors) and Neighbor-Joining. The statistics provided were taken with respect to the first split ID (see methods). For both Cassiopeia with and without priors, we used a cutoff of 200 cells and each instance of the ILP was allowed 5000s to converge on a maximum neighborhood size of 6000.

**Figure S20:**
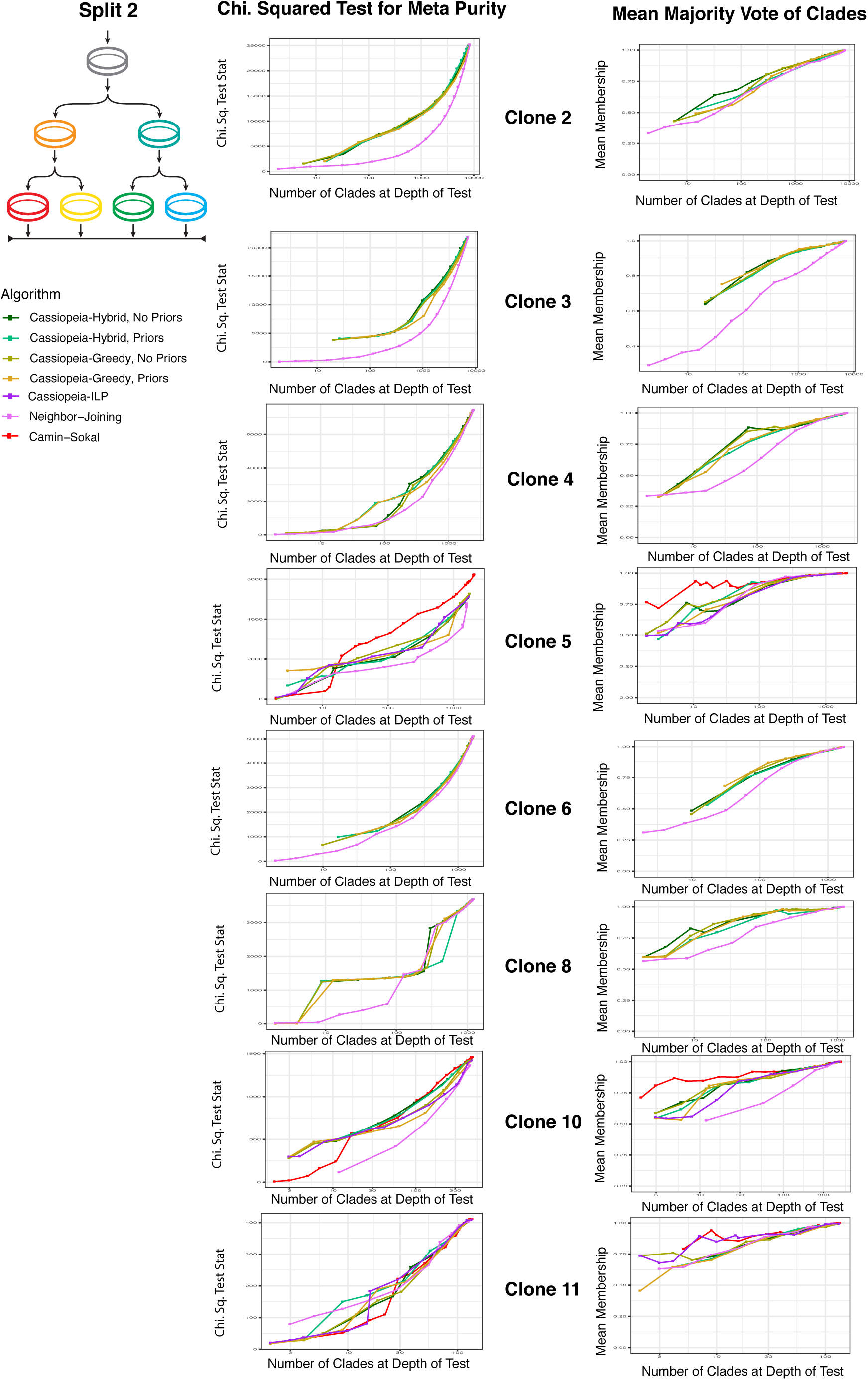
Evaluation of algorithms on *in vitro* lineage tracing clones, Second Split. Trees were reconstructed for the remaining clones in the *in vitro* dataset that consisted of more than 500 unique cell states. LG2, LG4, LG6, and LG8 passed this threshold and were reconstructed with Cassiopeia (with and without priors), greedy-only (with and without priors) and Neighbor-Joining. The statistics provided were taken with respect to the second split ID (see methods). For both Cassiopeia with and without priors, we used a cutoff of 200 cells and each instance of the ILP was allowed 5000s to converge on a maximum neighborhood size of 6000.

**Figure S21:**
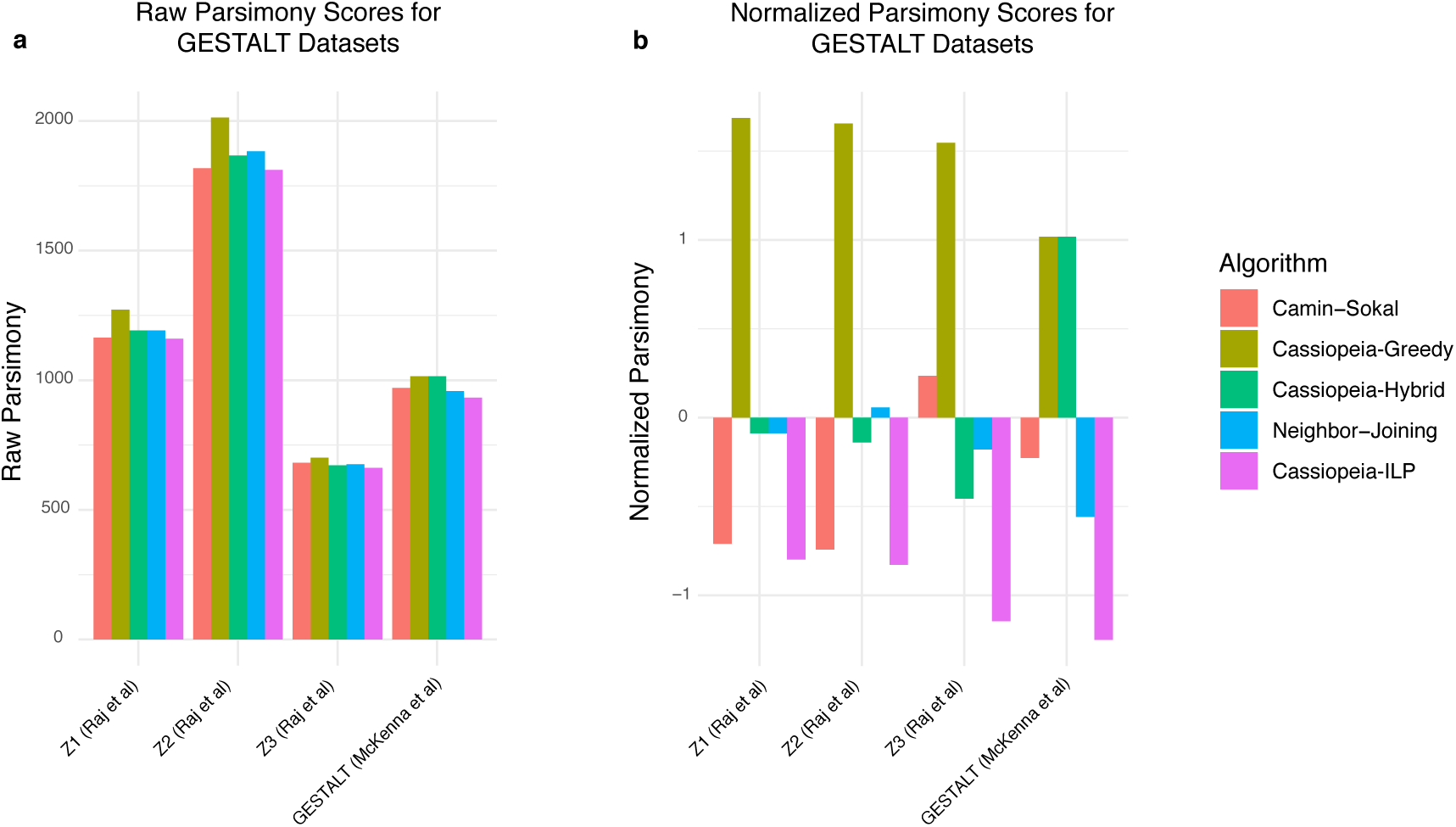
Parsimony Scores, Normalized and Raw, for GESTALT Reconstructions. (a) Raw and (b) normalized parsimony scores for the parsimony scores from the GESTALT datasets. Camin-Sokal, Neighbor-Joining, Cassipeia-Greedy, -Hybrid, and -ILP were run on datasets from Raj et al [42] and McKenna et al [38]. Raw parsimony scores are calculated as the number mutations present in a phylogeny (summing over the mutations along every edge of the tree). The normalized scores correspond to z-scores for each dataset.

**Figure S22:**
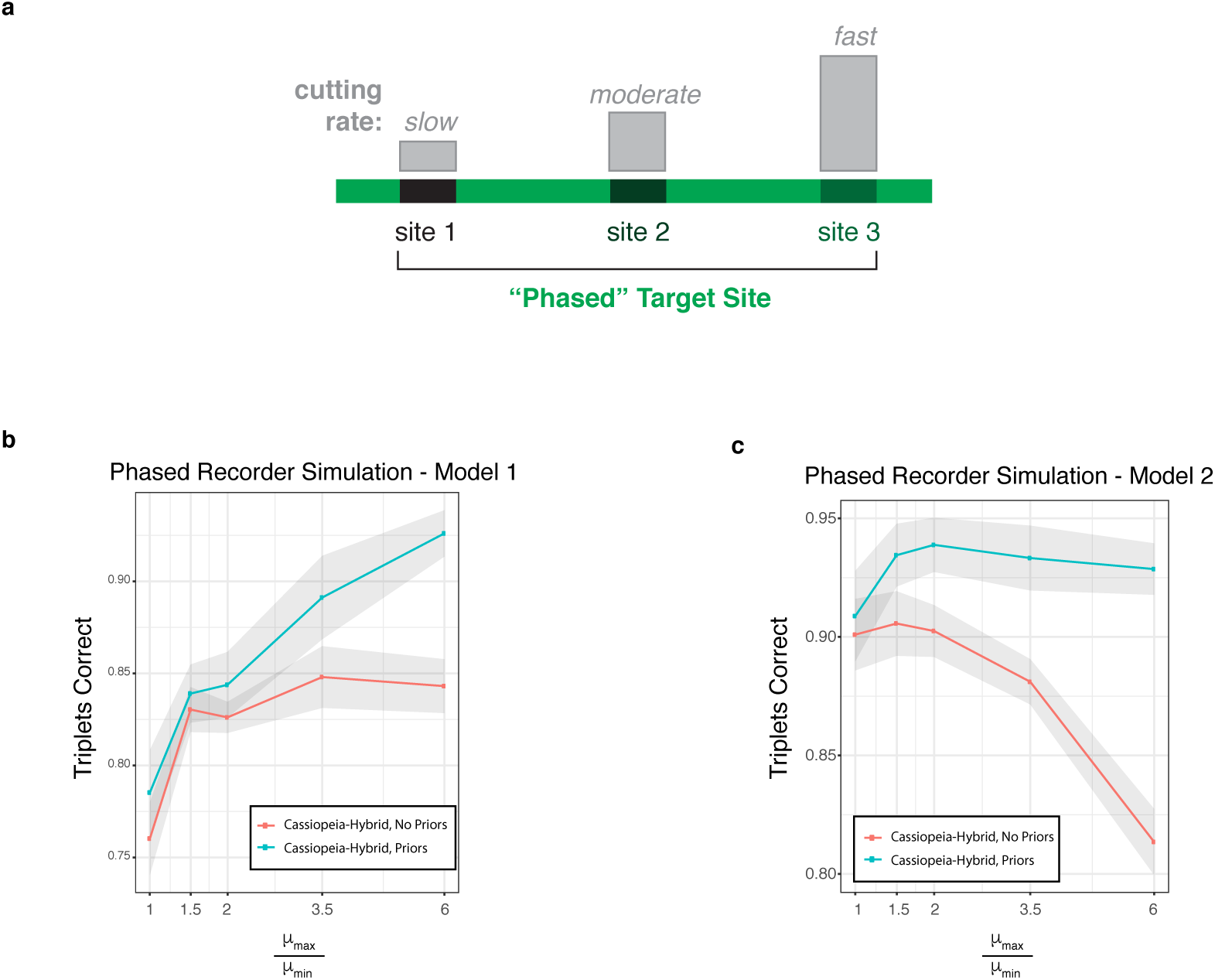
Simulations of Phased Recorder. (a) Design concept of the “Phased Recorder.” (a) We simulated a “phased” editor, where each character is mutated at variable rates. (b-c) We varied the amount each character could very across 5 different experiments and simulated using two different indel formation rate models. Each cell had 50 characters with 10 states per character and a mean dropout of 10%. The amount of mutation variability is described with the ratio between the maximum and minimum mutation rates 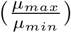. Standard error is represented by shaded area. (b) Model 1 consists of drawing indels from a negative binomial distribution *NB*(5, 0.5) where there are few “rare” indels. (c) Model 2 consists of drawing indels from the splined distribution of the empirical dataset’s indel formation rates, as used in other synthetic benchmarks.

